# High-Stability Polyimide-based Flexible Electrodes with IrO_x_ to Interface the Mouse Vagus Nerve

**DOI:** 10.1101/2021.05.10.442867

**Authors:** Tao Sun, Jessica Falcone, Christine Crosfield, Maria Fernanda Lopez, Joanne Peragine, Romil Modi, Rohit Sharma, Brian Baker, Gavin Anderson, Shubh Savani, Chunyan Li, Eric H. Chang, Harbaljit Sohal, Loren Rieth

## Abstract

**Objective:** We developed robust and cost-effective cuff *Flex* electrodes to facilitate bioelectronic medicine research in mouse models. They utilize polyimide (PI) as a dielectric insulation and iridium oxide (IrO_x_) for the electrodes, and are designed to interface small autonomic and somatic nerves (e.g. mouse vagus nerve).

**Approach:** *Flex* electrodes were made using micro-fabrication technology, and innovative integration processes were developed to enable reliable acute and chronic vagus nerve interfaces. The electrochemical properties of *Flex* electrodes were characterized. Moreover, accelerated aging at 57 °C and stimulation-stability (Stim-Stab) testing (10^9^ pulses at ∼ 1.59 mC/cm^2^/phase) were performed to evaluate the lifetime of the PI encapsulation and IrO_x_ electrodes, respectively. *Flex* electrodes efficacy was demonstrated by stimulating the mouse vagus nerve (∼100 µm) and measuring heart and respiratory rate changes as biomarkers.

**Results:** Cost effective and robust lead and connector integration strategies were demonstrated, including small helical leads that improved the lead elongation by > 7x. PI encapsulation had stable impedance spectra for at least 336 days for interdigitated electrodes. Stim-Stab testing using an aggressive paradigm and rigorous optical and electrical characterization, revealed that half of electrodes showed less than minor damage at the endpoints. A trend of decreasing respiratory rate with stimulation current reached statistical significance at 500 µA, demonstrating efficacy for *Flex* electrodes.

**Significance:** *Flex* electrodes offer demonstrated efficacy, low impedance (443 ± 32 Ω at 10^3^ Hz), excellent bench test stability, and cost-effective fabrication. Acute devices are easy to integrate, and mechanically robust chronic devices will be investigated *in vivo* in future studies. These characteristics make the electrodes well-positioned to advance bioelectronics medicine research by 1) enabling reliable studies with statistically relevant populations of acute mouse models, and 2) offering the potential for a technology that can be used in chronic studies, which scales to very small nerves.

## 1. Introduction

Bioelectronic medicine (BEM) is a rapidly growing field of healthcare, that endeavors to treat diseases and conditions through neuromodulation [1]. BEM continues to be explored for diseases and conditions including rheumatoid arthritis and inflammatory diseases [2], hypertension [3], pain [4], heart diseases [5], and metabolic diseases [6]. The vagus nerve is an important autonomic nerve for BEM, involving nerve recording, stimulation, and/or modulation to diagnose and/or treat disease. With the strong drive towards minimizing tissue damage of the delicate vagus nerve, unmet need to improve the electrode lifetime for chronic implantation in small animal models, and benefits for low electrode impedance to minimize required compliance voltage and improve stimulation safety, there is keen interest in developing long-lifetime, stable, low impedance, and flexible neural electrodes to interface the mouse vagus nerve.

Murine models, particularly mouse models, are critical for research in biomedical science, due to the availability of many disease models, genetic constructs, and pharmacological methods. However, the technology to interface delicate mouse vagus nerve for truly chronic endpoints (> 30 day) has been a challenge, despite being well-established in rat models. A wrappable platinum (Pt) microwire electrode was reported to chronically record from the cervical vagus nerve of awake murine models for 60 days [7]. Similarly, a chronic (day 36) rat vagus nerve stimulation (VNS) has been performed using a cuff electrode made from platinum iridium (Pt/Ir) wire threaded into a medical grade silicone tubing [8]. There are very few reports regarding chronic electrodes for VNS of mouse models, and fewer still involving flexible micro-fabricated thin-film electrodes. This gap is due to challenges that arise from the small 100 µm diameter of the nerve, including achieving stable low impedance materials for stimulation, making robust devices to interface the delicate nerve, and maintaining intimate contact with nerve for such a small nerve diameter [2], [9], [10].

Micro-fabrication allows cost-effective batch fabrication, scaling to small feature sizes through lithography, and the ability to use a sophisticated set of materials and structures to tune the properties of the neural interfaces. Micro-fabricated flexible split-ring electrodes enable the control of the flight direction for a forward-flying moth by minimally-invasive stimulation of the ventral nerve cord [11]. Micro-fabricated flexible transverse intrafascicular multichannel electrodes (TIMEs) and longitudinal intrafascicular nerve electrodes (LIFEs) are able to selectively activate subsets of axons within the nerve by transersely and longitudinally penetrating the peripheral nerves, respectively [12], [13]. Considering the delicate and fragile mouse vagus nerve, we choose to micro-fabricate a cuff electrode as a less invasive strategy.

Iridium oxide (IrO_x_) has a long history as an electrode material with high charge injection capacity (CIC), considerable stability, compatibility with microelectromechanical system (MEMS) processing, and can be synthesized on electrode sites by electroplating [14], electrochemical activation of iridium (AIROF) [15], and reactive sputtering in an oxidizing plasma (SIROF) [16]. However, there are only limited studies on its stability performance under aggressive stimulation conditions generated by pulsing the electrodes with high charge density (> 1.0 mC/cm^2^/phase), high frequency (∼10^3^ Hz), and a large number of stimulation cycles (∼10^9^), especially for IrO_x_ on the flexible polyimide (PI) substrate.

Leads to connect electrodes to electronics remain one of the dominate failure mechanisms for neural interface technologies, even for clinical devices such as deep brain stimulation implants [17]–[19]. The primary principles of electrode integration include minimization of tethering forces from the leads to the nerual interface, protecting the electrical connection from fatigue and breakage, ensuring the insulation of electrical components and minimizing the tissure responses [20]. In this study, a novel electrode integration using a small diameter helical lead was developed to enhance the mechanical elongation capability and bending fatigue tolerance for chronic implantation. Note that the bending fatigue studies have not yet been performed for these leads. Also studies have found that helical leads can scale up 100 channels, and significantly improved the lifetime of Utah slanted electrode arrays in peripheral nerves of human subjects [21]. Moreover, the cost of the integrated *Flex* eletrodes was estimated for both acute and chronic configurations, to demonstrate both can be cost-effective even when integrated at lab scale. Their cost effectiveness and ability to be integrated in a lab environment with good throughput enables the technology to support sufficient cohort sizes for robust statistical analysis.

Besides electrode failures caused by electrode integration or foreign body responses, insulating material failure is considered to be one of primary factors for neural electrode failure, accounting for 11.6% of electrode failure in a long-term non-human primates study [22]. Al_2_O_3_-parylene C insulated interdigitated electrodes (IDEs) exhibit a lifetime approximately equivalent to 40 months at 37 °C, three times longer than parylene insulated counterparts [23]. Although the stability of the bilayer structure was demonstrated to be 4.6 times greater than parylene alone for planar devices in another *in vitro* study [24], the results from the bilayer insulation testing on non-planar devices were less positive [25].

Polyimide (PI) is another popular polymeric material widely used for neural interfaces. However, there are only limited reports specifically regarding electrical characterization for the *in vivo* or *in vitro* lifetime of the PI insulation. Also, vastly different materials degradation rates have been observed based on the PI materials used and the associated processing conditions [26]. Alternative evidence based on physical characterization of implanted electrodes has shown reasonable stability for PI encapsulation during use in animal models, and recently with human subjects [27]. A study regarding the mechanical properties of PI, found that the Young’s modulus, stress and stain at breakage, and the stress at 10% strain for PI were stable over the soaking time of 20 months in PBS at 60°C [28]. Takmakov *et al*. developed a reactive accelerated aging test with hydrogen peroxide to evaluate the PI insulation for gold-coated tungsten microwire implants [29]. After a soaking time of 7 days at 87°C, the Pyre M.L PI (Industrial Summit Technology, Parlin, New Jersey) insulation loss on the rigid microwire electrodes was verified, using scanning electron microscopy (SEM) and electrochemical impedance spectroscopy (EIS). Details of the processing for the PI insulation used for the three dimentional electrodes are not known. One group did measure leakage impedance over time to generate lifetime curves to evaluate the effects of processing conditions. They determined that a lower curing temperature process (205°C) for PI bottom layer, followed by a normal high temperature curing process (300°C) for top PI layer, was found to yield a 7.5-fold improvement in lifetime of PI insulation for an electrode test structure [30]. Despite the advance to increase the lifetime of PI insulation, over 60% of channels from the electrode test structures failed after 30-day soak test in PBS at 40°C. Hence, from the perspective of the variation of electrochemical properties over time, the investigation into the lifetime of PI insulation still warrants further study for flexible neural electrodes.

In this study, a robust and cost-effective *Flex* thin film electrode using PI encapsulation with IrO_x_ electrode sites was fabricated via MEMS processes, and two integration strategies were developed for acute and chronic mouse vagus nerve interfaces. The estimated cost of integrated electrodes was broken down into each primary step. The PI insulation lifetime and the stability of IrO_x_ were evaluated, using acclerated aging and stimulation-stability (Stim-Stab) testing, respectively. More importantly, the efficacy of the *Flex* electrode was demonstrated by evoking bradycardia and apnea, which are common biomarkers for VNS. The decreasing trend in respiratory rate with increasing stimulation current reach statistical significance for our cohort at 500 µA.

## 2. Materials and methods

### 2.1 Micro-fabrication process

*Flex* electrodes consist of 1) pad/trace and 2) electrode metallization layers sandwiched between 3) bottom and top PI dielectric encapsulations layers, and were fabricated on 100-mm silicon (Si) wafers, using MEMS processes at University of Utah Nanofabrication Lab. Briefly, a PI bottom layer (PI2611, HD Microsystems, Parlin, New Jersey) was spin-coated on the wafer, followed by a curing process in nitrogen atmosphere at 300°C, yeilding a 7 µm thick film. A lift-off photolithographic process was then conducted to pattern the pad and interconnection metal traces, using a thick (10 µm) AZ 9260 photoresist (PR) layer. After sputtering a titanium (Ti)/platinum (Pt)/gold (Au) (100/250/425nm) stack layer, the lift-off process was performed. Similarly, the IrO_x_ electrode sites were fabricated using a sputter desposited Ti/Ir/IrO_x_ (100/250/420 nm) stack, patterned by standard liftoff photolithography. The 7 µm top PI dielectric encapsulation was then spin-coated. Subsequently, a double layer of thick PR layer, AZ 9260 with a total thickness of ∼20 µm, served as the soft mask layer to define the *Flex* electrode outline, and open bonding pads and electrode sites. The PI was etched using oxygen plasma in an Oxford ICP reactive ion etcher (Oxford PlasmaLab 100, Oxford Instuments, Abingdon, United Kingdom), with the pad and electrode metalization serving as etch stops using a process that etches through the 14 µm of PI for the electrode outline. After the oxygen plasma reactive ion etching (RIE) process to etch the exposed PI, the PR was stripped with acetone. The exposed IrO_x_ electrode sites are 1418×170 µm^2^, resulting in a geometrical surface area of 0.00241 cm^2^ (2.41 × 10^5^ µm^2^) when the rounded corners are accounted for. The electrodes have a 0.9 mm distance from center-to-center.

The *Flex* electrodes are manually released from the Si wafer, using a scapel or tweezers. The current 100 mm wafer process can accommondate up to 160 of the *Flex* electrodes. Note that this is a relatively large design for the lab, and therefore represents a relatively low range for the number of die that can be accommondate on a wafer.

### 2.2 Surface characterization

A digital microscope (VHX-7100, Keyence Corporation of America, Itasca, Illinois) was utilized to capture optical images of the micro-fabricated *Flex* electrodes, and integrated *Flex* electrodes with leads and connectors for acute and chronic VNS. Surface morphology of IrO_x_ electrode was visualized using a scanning electron microscope (SEM, Apreo™, Thermo Fisher Scientific, Hillsboro, Oregon), under a backscattered mode, as backscattered SEM has more Z-contrast to enhance contrast between noble-metal from the electrodes and traces compared to the polymer dielectric materials.

### 2.3 Electrode integration

#### 2.3.1 Integration for acute vagus nerve stimulation

Two different integration strategies were developed to enable reliable and cost-effective leads for electrical connections to acute and chronic vagus nerve electrodes. For acute VNS, the silicone insulation was stripped to expose the thin multi-stranded copper wires (AS 155-36, Cooner Wire Co., Chatsworth, California), and the wires were soldered onto the bonding pads of the *Flex* electrodes. Soldering used a SAC 305 lead-free solder paste with water-soluble flux (Indium Corp., Clinton, New York), and a fine-tipped soldering iron. Subsequently, the unmelted solder paste and water-soluble flux were removed by thoroughly rinsing and irrigating the bonding pad area in warm deionized water and then carefully dried. The solder joints were strengthened by applying a small amount of medical grade epoxy (Loctite M-31CL, Rockhill, Connecticut). After a curing step in an oven (Heratherm™, Thermo Fisher Scientific, Waltham, Massachusetts) at 60 °C for 45 min, the medical grade silicone (MED-4211, NuSil Technology LLC, Carpinteria, California) was applied at the junction between the leads and the *Flex* electrode to further encapsulate and electrically insulate the solder joints and accommodate mechanical stress (Fig. 2a). To validate that the integration process provides enough strength for acute VNS, tensile testing was performed using an Instron mechanical tester for *Flex* electrodes with and without the epoxy reinforcement of the soldered wires.

#### 2.3.2 Integration for chronic vagus nerve stimulation

Chronic VNS requires more reliable lead and connector system for *Flex* electrodes. Instead of the Cooner wire, a flexible and stretchable lead using helically formed insulated Pt wires overmolded in silicone with a head-cap using an Omnetics connector was developed for chronic VNS. Briefly, 4 pieces of 16-cm-long multi-stranded and fluorinated ethylene propylene (FEP) insulated Pt wires (diameter = 0.1 mm, Medwire Corp., Mount Vernon, New York) were cut as the electrical connection for channel 1, channel 2, ground 1 and ground 2 (individual grounds), respectively. The Pt wires were tied on a steel rod mandrel with a separation of 5 mm, and then were fixed in place using a small drop of super glue (EA 9017™, Henkel Corp., Rocky Hill, Connecticut). To make the helical wires, the Pt wires for electrode connection (channel pair) crossed over those for ground (ground pair) as shown in Fig. S1a (Supplementary materials), and then the mandrel was turned to wrap the Pt wires. After 5 cm-long helical Pt wires were prepared with an approximate density of 25 turns/cm, the helical Pt wires were cut off using a scissor. Subsequently, the helically formed Pt wires were slid off the rod (Fig. 2b). The helical lead was then overmolded in a 1 mm diameter and 5 cm long cylindrical mold machined of aluminum (Fig. S1b, Supplementary materials). The two halves of the mold were pre-loaded with NuSil MED-4211 silicone, and the helical pre-form was placed in the lower mold, followed by carefully placing the top mold and compressing them together using integrated screws. Subsequently, the curing process was performed in an oven at 60 °C for 1 hour, followed by carefully detaching the integrated electrode from the mold. Both ends of the helical Pt wires were carefully and mechanically de-insulated and cut to the designed length using a razor and surgical micro-scissors, and the channel pair of helical Pt wires were soldered on the bonding pads of a *Flex* electrode. The other end of the helical lead was soldered to the metal pads of a PCB that the 16-channel Omnetics connector was reflow soldered upon. The solder joints at each end were encapsulatd with epoxy and silione.

For the comparision in mechanical properties, straight Pt wires were overmolded with the previously described mold (Fig. S1b), and then soldered on the bonding pads of the *Flex electrodes*, followed by epoxy reinforcement and silicone protection at the solder joints.

### 2.4 Tensile test

Mechanical properties of integrated electrodes, such as maximum load and extension at breakage were determined via a uniaxially tensile test, using an Instron mechanical testing system (Instron 5566, Norwood, Massachusetts) with a 100 N load cell.

The integrated electrodes for acute VNS were tested with and without the medical grade epoxy on the soldered joints. The Cooner wires were fixed by the upper grip of the Instron tester, while the integrated electrode was secured in the lower grip with a length of 20 mm between two grips. The tensile test was performed at an extension rate of 0.25 mm/sec until tensile failure of the specimens.

Similarly, mechanical properteis of integrated electrodes with and without the helical Pt lead structure were examined by the same approach as described above, for chronic VNS. The exposure length between two grips was 50 mm.

### 2.5 Electrochemical properties

### 2.5.1 Electrochemical impedance spectroscopy (EIS)

EIS was used to characterize the impedance of *Flex* electrodes, and to evaluate degradation in the PI encapsulation layers using interdigitated electrode (IDE) structures. To monitor the PI insulation behavior as a function of soaking time, EIS measurements were performed longitudinally for IDE devices using both 3-electrode and 2-electrode configurations, via a potentiostat (Reference 600 +™, Gamry Instruments, Warminster, Pennsylvania) in 1x phosphate-buffered saline (PBS) at room temperature. Typical 3-electrode configuration consists of one finger of the IDE device as the working electrode, a deinsulated Pt wire as the counter electrode and a saturated Ag|AgCl reference electrode, while in the 2-electrode configuration, the other finger of the IDE device serves as the combined counter electrode and reference electrode. This approach has been utilized by our lab in the past, as well as in other reports [31][32], and facilitates distinguishing between internal (between figures within the IDE) and external (through the PI insluation layer) electrical leakage pathways.

Impedance spectra were collected using a standard protocol with 10 points per frequency decade from 1 to 10^5^ Hz at 25 mV delivered to the working electrode [33]. To minimize the electromagnetic interference, all EIS measurements were performed in a Faraday cage at room temperature. Before and after the stimulation stability test (section 2.6), the impedance magnitude (|Z| (Ω)) and phase (°) of the *Flex* electrodes as a function of frequency were determined using the 3-electrode configuration.

#### 2.5.2 Cyclic voltammetry (CV)

To evaluate the electrochemical characteristics and charge storage capacity (CSC) of *Flex* electrodes (n=24), CV measurements were conducted, using a sweep rate of 50 mV/sec between the electrolysis potential (water window) limits of -0.6 and 0.8 V. The same potentiostat and 3-electrode configuration were employed to collect CV data for electrodes in 1x PBS (0.15 mM) at room temperature. The CV sweep was repeated at least three times for each electrode, and only the last sweep was used to calculate the CSC value. The CSC value is calculated by integrating the cathodic current density across the potential range [33].

#### 2.5.3 Voltage transient (VT)

Voltage transient (VT) waveforms in response to the stimulation current pulses without interpulse bias were acquired to determine the charge injection capacity (CIC) for *Flex* electrodes. Electrodes were characterized with symmetric, charge-balanced, cathodic leading, biphasic pulses with pulse-widths of 200 µs, 300 µs and 400 µs per phase, at a pulse rate of 10^3^ Hz (Fig. S2, supplementary materials). The cathodic and anodic phases are separated by a 20 µs interphase delay. A stimulator (STG4008, Multi Channel Systems MCS GmbH, German) was utilized to deliver the stimulation pulses in a monopolar configuration with a large-area Pt ground/counter electrode.

The maximum cathodic potential excursion (*E*_*mc*_) was determined in a manner of subtracting the access voltage (*V*_*ac*_), associated with Ohmic resistance from the solution, from the maximum negative voltage in the transient. To avoid water electrolysis, the *E*_*mc*_ should not exceed the cathodic limit of the water window (−0.6 V for IrO_x_ as is consistent with the CV measurements). CIC was calculated by the current amplitude and pulsing time, corresponding to the *E*_*mc*_ at the water window.

### 2.6 Stimulation stability test (Stim-Stab test)

The stability of the IrO_x_ electrodes for stimulation was assessed by the stimulation stability (Stim-Stab) test performed at room temperature. Electrode sites for *Flex* electrodes were immersed in 1x PBS in small glass vials (Fig. S3, supplementary materials). One channel of the *Flex* electrode was stimulated using a deinsulated Pt wire as the counter/ground electrode, while the second channel was retained as a control without stimulation current (unstimulated channel). A charge-balanced, cathodic-leading, symmetrical stimulation paradigm was used (Fig. S4, supplementary materials). The pulse-width per phase, interphase period, and interpulse period are 240 µs and 40 µs, and 480 µs, respectively. Two current amplitudes (12 mA and 16 mA), corresponding to a charge denstiy of 1.20 and 1.59 mC/cm^2^/phase, were used in the Stim-Stab test (n=3 for each current amplitude). As a control and to monitor the current pulses generated by the stimulator, a 1 kΩ resistor was connected to an output channel of the stimulator, and the voltage waveform of the resistor was displayed using an oscilloscope (DSOX2004A, Keysight Technologies, Santa Rosa, California). The voltage waveform of control and stimulated channels were recorded using a data acquisition system (Multifunction DAQ, National Instruments Corporation, Austin, Texas) and displayed in real-time VT waveforms using customized MATLB (2016a, Mathworks, Natick, Massachusetts) code. A sinppet of voltage waveforms that included 2 or 3 pulses was collected periodically once per 250,000 cycles (250 s).

After 1 billion pulses (11.6 days at 1000 pulses per second), all electrodes were gently rinsed using DI water, and electrical properties of unstimulated and stimulated electrode channels were then characterized for comparison, using EIS, CV and VT. The electrode sites were characterized for damage using reflected and transmitted light optical microscopy. Electrode damage was quantified, by analyzing the area damaged or degraded by the stimulation paradigm, using ImageJ (National Institutes of Health, Bethesda, Maryland).

### 2.7 Mouse vagus nerve stimulation

#### 2.7.1 Electrode Implantation Surgery

The efficacy of the electrodes were evaluated *in vivo* by stimulating the vagus nerve of mouse models, and monitoring heart and respiratory rates as the primary biomarker, as bradycardia and apnea are some of the most consistent physiological changes with VNS, which also respond and recover with short time constants. This study and all experimental protocols (No. 2016-029) were approved by the Institutional Animal Care and Use Committee (IACUC) at the Feinstein Institutes for Medical Research, Northwell Health System (Manhasset, NY, USA), which follows the National Institute of Health (NIH) guidelines for the ethical treatment of animals.

Male BALB/c mice (Charles River Laboratories 20-25 g, n=3) were used to evaluate the ability to interface the cervical vagus. Animals were anesthetized in an induction chamber at 3% isoflurane and then transferred to a nose cone at 1.5% isoflurane. A heat mat was placed under the animal for the duration of the procedure. Puralube (Dechra, UK) was placed on the eyes, and hair in the anterior cervical region was removed with Nair (Church & Dwight Co, NJ) and the surgical site was sterilized with alternating povidone-iodine and alcohol pads prior to incision.

While in the supine position, a midline cervical incision was made on the neck and the left vagus nerve was isolated from the carotid artery as previously described [7]. The vagus nerve was de-sheathed by removing the thin connective tissue surrounding the nerve using fine surgical forceps for blunt dissection under magnification.

#### 2.7.2 Vagus Nerve Stimulation and EKG Recording

A 2-channel *Flex* electrode was placed directly under the nerve and sutured closed to achieve intimate contact between the electrode and nerve surface. A ground electrode was placed under the contralateral salivary gland. Stimulation was performed using an Intan Technologies RHS2000 Stim/Recording System (Intan Technologies, Los Angeles, California). Subcutaneous needle electrodes were inserted to acquire electrocardiogram (EKG) using the Intan System.

Biphasic pulses with a pulse width of 333.4 µs were delivered to a single channel on the *Flex* electrode at 30 Hz for 6.667 s (or 200 pulses). Stimulation currents were tested from 100-500 µA, with 100 µA step sizes. Each current value was tested three times with > 30 sec rest between stimulation trains. Typical impedance value at 10^3^ Hz of the electrodes were 443 ± 32 Ω and this leads to a maximum possible current of 21.8 mA being delivered with the ± 9V compliance voltage of the stimulator. Therefore, we could successfully deliver all the stimulation current parameters used in this study, even with the increases in impedance often observed during acute *in vivo* electrode placement.

Both the rostral and caudal channels were independently stimulated, and the order of channel stimulation was randomized to control for potential order effects.

#### 2.7.3 Offline Data Analysis

Electrophysiological recordings for each elctrode channel (n=6) in the form of .RHS files were imported into MATLAB, and the heart beat and respiratory electrophysiological signals were extracted. Heart rate and respiratory rate were then calculated, using customized MATLB code. Briefly, the QRS complex peaks of the EKG were identified and the time between each peak was calculated for beats per second (BPS). Similarly, EMG oscillations associated with repiratory muscles were used to monitor respiratory rate. The stimulation timings were also recorded and used to calculate the average heart rate (HR) and respiratory rate (RR) before and during stimulation. The percent change of the HR and RR were calculated for each stimulation current by comparing the HR and RR during stimulation to the values immediately prior to stimulation. A one-way ANOVA with multiple comparisons was performed on the percent change data using GraphPad (2012, GraphPad, San Diego, California).

### 2.8 Statistical analysis

All experiments were conducted independently at least three times, and results are presented as mean ± standard deviation. Multiple comparisons were analyzed using one-way ANOVA followed by Bonferroni correction, and *p* values <0.05 were considered statistically significant. Each channel of the IDE devices and *Flex* electrodes used in EIS, CV, and VT measurement is considered as an independent sample in the statistical analysis. Similarly, each channel of the *Flex* electrode in Stim-Stab and in *vivo* testing was regarded as independent in the statistical analysis.

## 3. Results

The optical image of *Flex* electrodes with suture holes is shown in Fig. 1a. The interconnection and bonding pads are the highly reflective gold colored structures, labelled at the top aspect of the electrodes, while the IrO_x_ electrode sites are black. The suture holes are designed specifically to help secure the electrodes on the nerve for chronic implantation, and are mechanically strengthened by a ring of the Ti/Pt/Au trace metallization. Fig. 1b is a backscattered SEM image, showing the surface morphology of the IrO_x_ electrode sites with its nano-scale dendritic structure. The nanoporous structure can increase the effective electrochemical surface area, which can enhance the electrochemical properties of the *Flex* electrode. This can include decreasing the impedance, and increasing the CSC and CIC relative to a smooth dense film.

**Figure 1.**
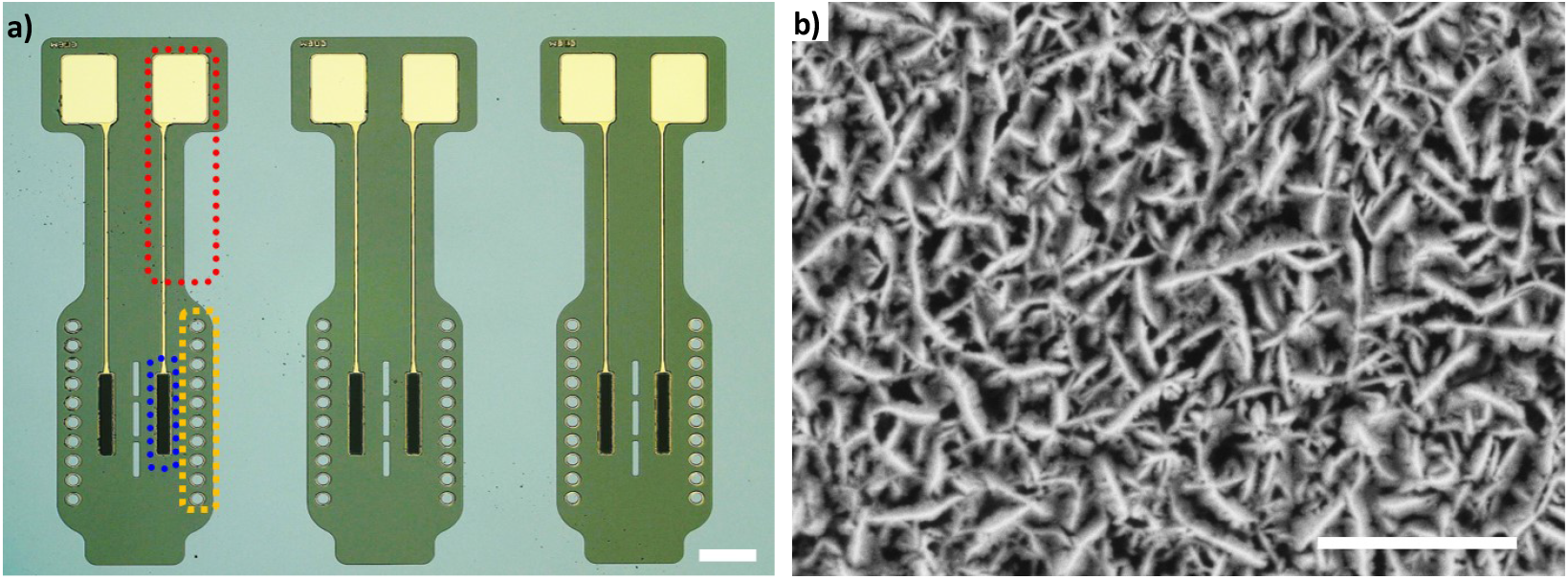
(a) Optical image of the micro-fabricated *Flex* IrO_x_ electrodes on the Si wafer, scale bar=1.0 mm (the red dot box indicates the interconnection and bonding pad, the IrO_x_ electrode site is labeled by the blue dot box, and the orange dot box shows suture holes) (b) SEM image of the IrO_x_ electrode site, the scale bar=500 nm

Fig. 2a presents an optical image of an integrated *Flex* electrode in the configuration for acute implantation. The solder attached Cooner wires, which are strengthened by epoxy and potted with silicone, are used as an inexpensive and robust lead to connect electrodes to electrophysiology (stimulation and recording) and characterization instruments. The optical image of the integrated electrode for chronic VNS and a higher magnification image of the electrode are presented in Fig. 2c and 2d, respectively. Two *Flex* electrodes were integrated with the PCB and Omnetics connector to enable measuring conduction velocities of evoked active in the vagus nerve, and can also be used for bilateral interface to the left and right vaus nerves. Each *Flex* electrode has two Pt wires with deinsulated tips (Fig. 2d), which serve as the individual ground and/or reference electrodes for each *Flex* electrode. Fig. 2e presents the load-extension curves of the integrated acute configuration electrodes with and without the epoxy reinforcement. All integrated electrodes with the epoxy reinforcement exhibit a higher load (7.29 ± 2.06 N) and longer extension (7.67 ± 3.78%) at tensile failure compared to electrodes (1.96 ± 0.82 N and 1.87 ± 0.54%, respectively) without epoxy reinforcement. Fig. S5a and b (Supplementary materials) reveal that the epoxy reinforcement at the solder joints offers 3.7 and 4.1 times greater the maximum load and extension, respectively (*p* < 0.05).

**Figure 2.**
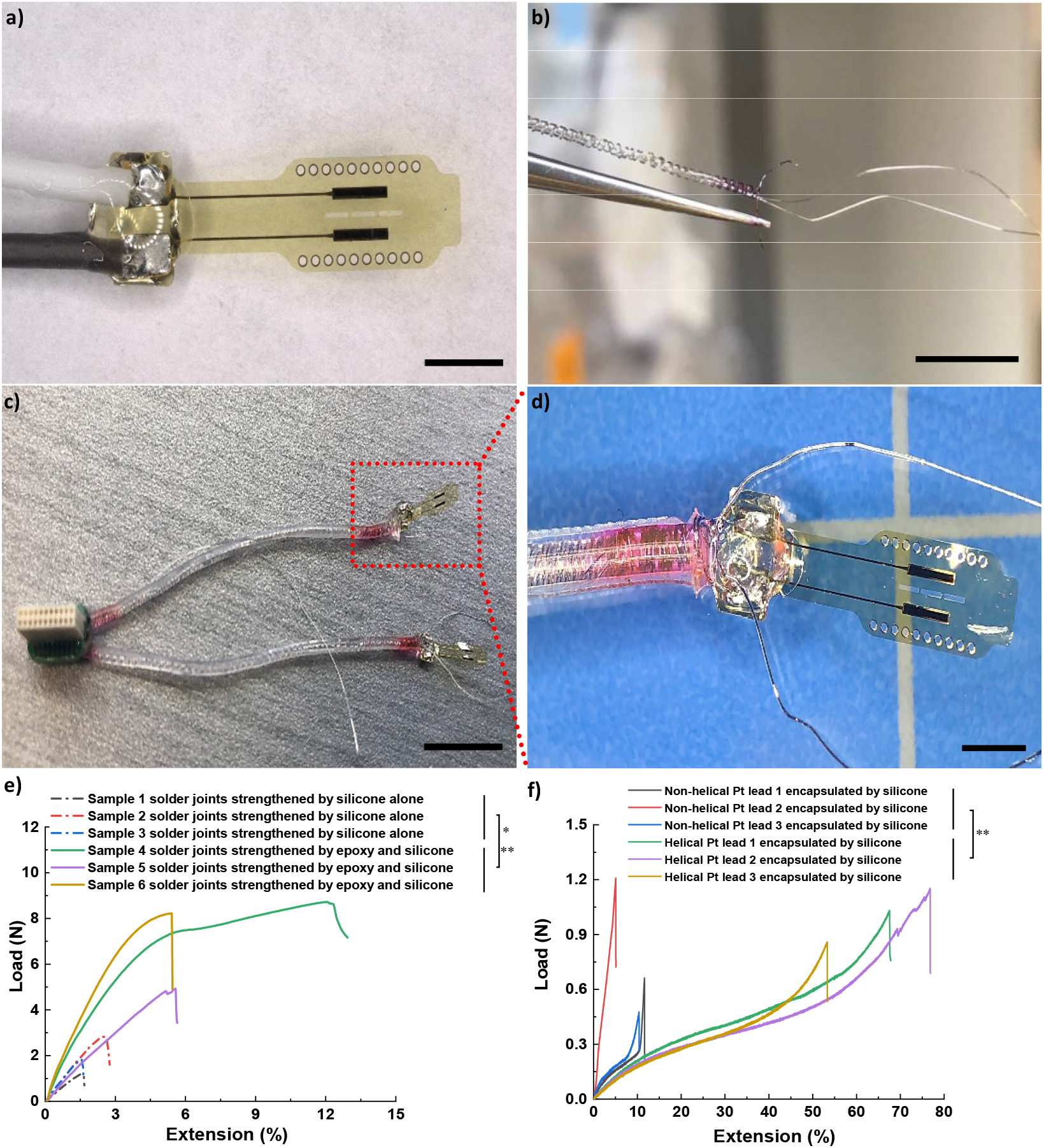
Integration of *Flex* electrodes. a) optical image of the integrated electrode for acute vagus nerve stimulation, the scale bar=2.0 mm, b) optical image of the helical Pt wires after the removal of the rod, the scale bar=5.0 mm, c) optical image of the integrated eletrode for chronic vagus nerve stimulation, the scale bar=1.0 cm, d) high-magnification optical image of the integrated electrode in Fig. 2c, the scale bar=2.0 mm, e) load-extension curves of the integrated acute configuration electrodes with and without the epoxy reinforcement (* *p* < 0.05 for maximum load, ** *p* < 0.05 for maximum extension), f) load-extension curves for the integrated chronic configuration electrodes with non-helical Pt leads and helical Pt leads (** *p* < 0.05 for maximum extension)..

Fig. 2f shows the load-extension curves, which compares the integrated chronic configuration electrodes with helical and straight Pt wire configurations. Although there is no statistical difference in terms of the maximum load at tensile failure (Fig. S5c, *p* > 0.05), the maximum extension was improved 7.3 times from 9.02 ± 3.48% to 65.89 ± 11.82%, due to the electrodes integration with helical Pt leads (Fig. S5d, *p* < 0.05). The dramatic increase in extention capability not only offers more reliability for the integrated electrodes, but also helps minimize tethering forces on the nerve to improve safety of the electrode.

The cost of the integrated electrodes is a crucial factor for both the fabrication of research electrodes, and for the business model to commercialize electrodes, but seldomly reported in literature. For acute electrodes, low cost is needed to facilitate large cohorts to accurately determine the statistical significance of findings, and due to the availability of inexpensive alternative such as reusable hook electrodes. Chronic electrodes demand better connectors and integration, which increases costs, but are often used in smaller cohorts due to the demands of chronic animal studies. In this study, the cost of the micro-fabrication and integration process are broken down by processes and components, and listed in Table 1 and 2, respectively. For the micro-fabrication process, the external academic pricing was used in the University of Utah Nanofabrication Lab, and each micro-fabricated electrode costs 10.5 US dollar (USD). The metalization of pad, interconnection and IrOx accounts for almost 50% of entire fabrication cost. The cost of micro-fabricaiton can be significantly decreased by 1) optimizing the number of electrodes per wafer, and 2) scaling the process up to larger wafers (e.g 200 mm).

**Table 1.** Cost of the micro-fabrication processes based on the external academic pricing.

| Items | Steps | Cost/wafer (USD) | Cost % |
| --- | --- | --- | --- |
| Micro-fabrication | Wafer preparation | 75 | 4.5 |
|  | 1<sup>st</sup> PI layer | 167 | 9.9 |
|  | Metalization for pad and interconnection | 410 | 24.4 |
|  | IrO<sub>x</sub> layer | 378 | 22.5 |
|  | 2<sup>nd</sup> PI layer | 195 | 11.6 |
|  | Reactive ion etching | 390 | 23.2 |
|  | Inspection | 65 | 3.9 |
| In total |  | 1680 | 100 |
| **Cost/electrode (USD)** |  | **10.5** |  |

**Table 2.** Cost of integration processes for acute and chronic configuration *Flex* electrodes.

| Items | Steps | Cost/ (USD) | Cost % |
| --- | --- | --- | --- |
| Integration for acute implantation (10 electrodes/batch) | Soldering | 52 | 44.8 |
|  | Epoxy reinforcement for joints | 22 | 19.0 |
|  | Silicone protection for joints | 32 | 10.6 |
|  | Inspection | 10 | 8.6 |
| In total |  | 116 | 100 |
| **Cost/electrode (USD)** |  | **11.6** |  |
| Items | Steps | Cost (USD) | Cost % |
| Integration for chronic implantation (4 electrodes/batch) | Helical leads preparation | 240 | 42.5 |
|  | Silicone encapsulation for leads | 52 | 9.2 |
|  | PCBs and Omnetics connectors | 160 | 28.3 |
|  | Soldering | 65 | 11.5 |
|  | Epoxy reinforcement for joints | 22 | 3.9 |
|  | Silicone protection for joints | 22 | 3.9 |
|  | Inspection | 4 | 0.7 |
| In total |  | 565 | 100 |
| **Cost/electrode (USD)** |  | **141.25** |  |

The cost of the integration process for chronic VNS is 141.25 USD, and more than 12 times higher than that (11.6 USD) for acute VNS, due to the usage of helical Pt leads, PCBs and Omnetics connectors, and the increased labor to form the helical lead and perform the integration. In addition, PCBs and Omnetics connectors account for more than 25% of the cost for the integrated chronic VNS devices. The total cost of the integration process for chronic VNS is more than 13 times higher than that of the micro-fabricated electrode. Developing surgical approaches and integration approaches compatible with lower cost connectors (e.g. PlasticsOne) and lead wires could help decrease the cost of chronic electrodes, which can in turn facilitate larger cohorts for improved statistics, and more lattitude for discovery driven research.

Fig. S6a (supplementary materials) shows the optical image of an integrated IDE, and Fig. 3 presents the representative Bode and Nyquist plots for the impedance of a IDE device (IDE 01-channel 01) as a function of time. IDEs are used to evaluate the stability of PI and its ability to encapsulate trace metalization [34]. Bode and Nyquist plots of other IDEs are presented in Fig. S7-9 (supplementary materials). 3-electrode configuration for IDE devices can identify the PI insulation degradation surrounding one electrode of the IDE structure, as the current path is from one IDE electrode to the counter electrode in the solution (Fig. S6b). The 2-electrode configuration measures current flow between the IDE electrodes, which can detect the encapsulation degradation between the electrodes through internal pathways (e.g. the interface between top and bottom dielectric layers). The structures can also be sensitive to external pathways, which can make discrimination of these failure mechanism a challenge, which is consistent with findings from other labs (Fig. S6c) [31]. Note that the use of *decoration* techniques to identify external pathway failures can be effective in disambiguating the failure pathway and localizing failure points, as has been shown for Utah arrays [25]

**Figure 3.**
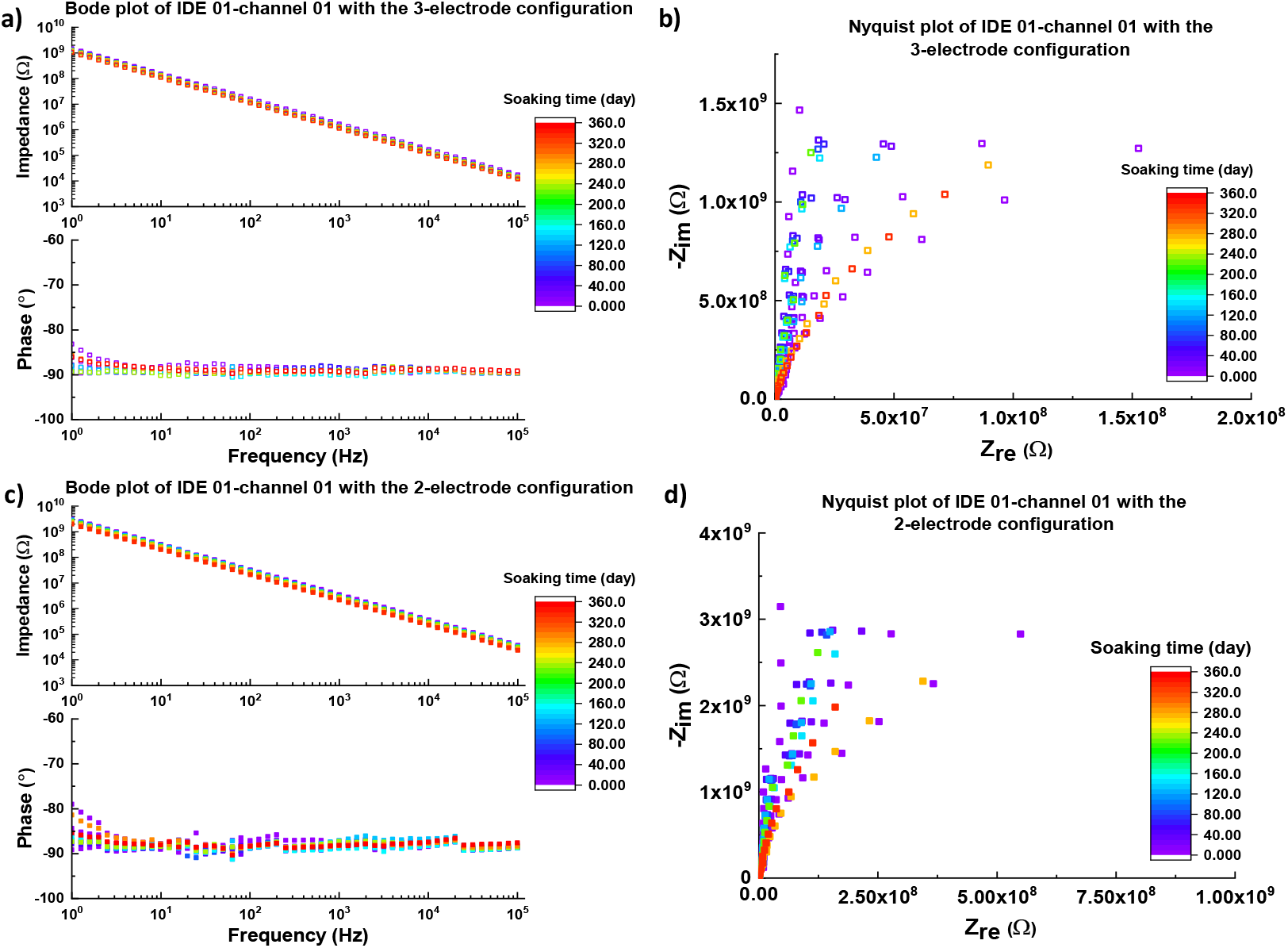
Impedance spectra of the representative IDE device (IDE 01-channel 01) as the function of the soaking time. a) Bode plot of IDE 01-channel 01 device as the function of the soaking time under 3-electrode configuration, b) Nyquist plot of IDE 01-channel 01 device as the function of the soaking time under 3-electrode configuration, c) Bode plot of IDE 01-channel 01 device as the function of the soaking time under 2-electrode configuration, d) Nyquist plot of IDE 01-channel 01 device as the function of the soaking time under 2-electrode configuration. Impedance spetra of other IDEs are shown in the supplimentary materials (Fig. S7-9).

For both 3-electrode and 2-electrode configurations (Fig. 3a and c), the slopes of the impedance magnitudes are nearly constant at -1, from 1 to 10^5^ Hz. Consistent with the slope of the impedance magnitude, the phase angle of impedance for IDEs is near -90°, indicating the expected capacitive behavior. Although the impedance of the IDE slightly decreased across the spectrum with soaking time due to the polymer hydration, water ingress, and degradation of the PI insulating layer, both the slope of impedance and the phase of the IDE did not significantly alter with time. Moreover, no semi-circle profile or Warburg impedance (a diagonal line with an slope of 45°) appears in Nyquist plots (Fig. 3b and d), revealing that the IDE trace metallization was not exposed to PBS. The IDE maintained capacitive behaviour, and the impedance did not fall below the failure criteria (1-Hz impedance < 10^8^ Ω), so survived in the accelerated aging test. Likewise, other IDEs did not failed and have an equivalent lifetime of ≥ 336 days at 37 °C.

The electrochemical properties of all IDE devices during the accelerated aging test at 57°C are summerized by plots that include the mean impedance and phase of all IDE devices with 3-electrode and 2-electrode configurations (Fig. 4a). As in 2-electrode configuration, the current passed through the PI insulation over the metal fingers twice. This is consistent with the approximately 2.2-fold higher impedance across the spectra for measurements made using the 2-electrode configuration, compared to the 3-electrode configuration. The slopes of spectra from both configurations did not change throughout the accelerated aging test and were close to -1. The filled error bars in Fig. 4a quantifies the standard deviation and is very narrow for both impedance and phase, revealing that the impedance characteristics of the micro-fabricated IDEs were highly consistent.

**Figure 4.**
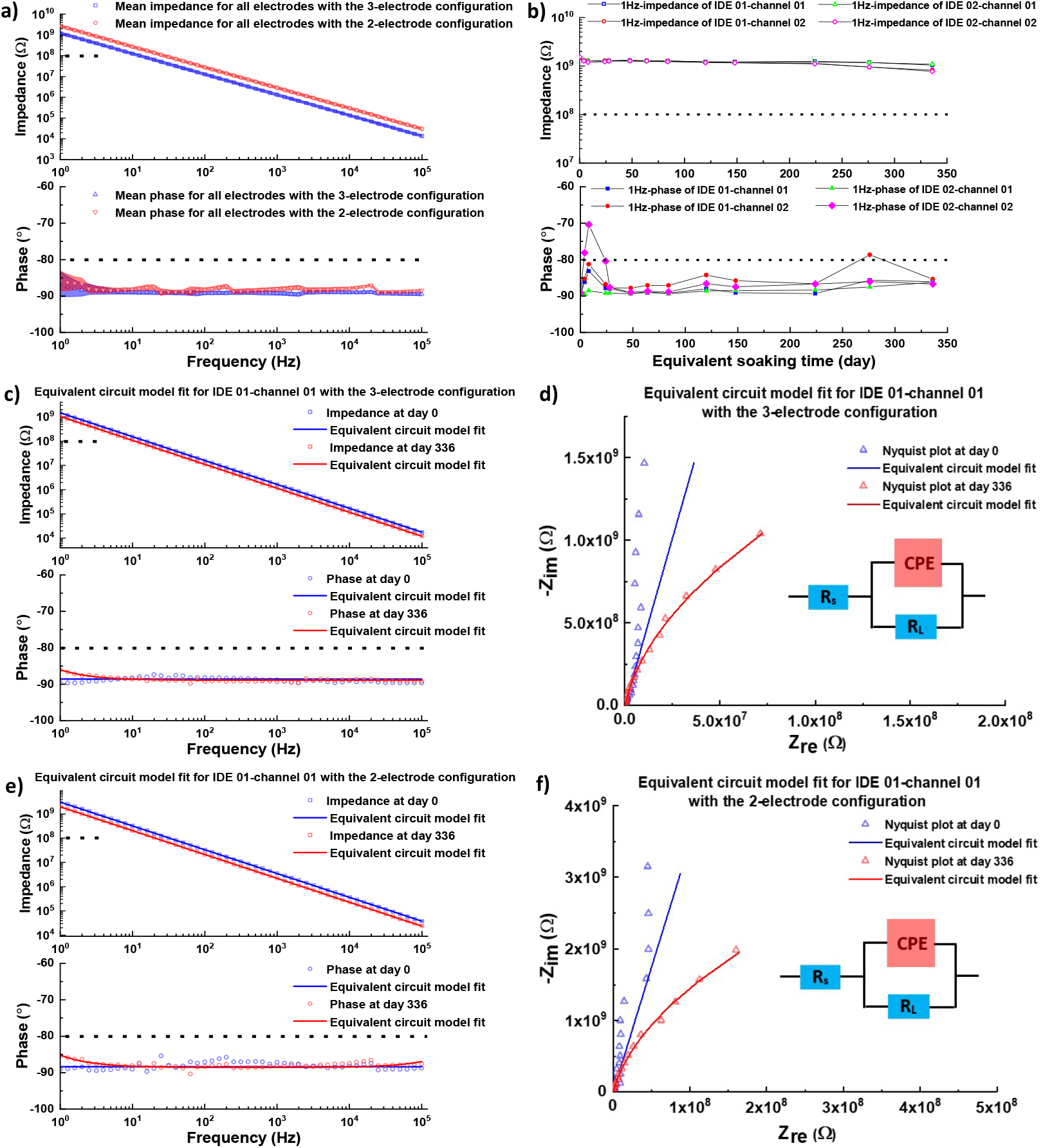
Electrochemcial properties of IDE devices as the function of the soaking time and the equivalent circuit model fit for the impedance spectra of IDE devices. a) mean Bode plot of all IDE devices throughout the entire soaking time (shaded areas indicate the standard deviation, and black dot lines show failure creteria), b) 1-Hz impedance and phase of each IDE devices as the function of the soaking time (black dot lines indicate the failure creteria), c) an equivalent circuit model fit for the Bode plot of an representative IDE device (IDE 01-channel 01) with the 3-electrode configuration, (black dot lines indicate the failure creteria) d) an equivalent circuit model fit for the Nyquist plot of an representative IDE device (IDE 01-channel 01) with the 3-electrode configuration, e) an equivalent circuit model fit for the Bode plot of an representative IDE device (IDE 01-channel 01) with the 2-electrode configuration (black dot lines indicate the failure creteria), f) an equivalent circuit model fit for the Nyquist plot of an representative IDE device (IDE 01-channel 01) with the 2-electrode configuration.

According to previous studies [23], [24], two metrics are employed to determine the failure in insulation materials which included Parylene-C and Alumina+Parylene-C in the cited studies, which include 1) impedance at 1 Hz no less than 10^8^ Ω, and 2) phase angle at 1 Hz no larger -80° at each time point. These criteria are choosen to be responsive to leakage current limits for medical device eletronics such as those identified in ISO 14708-1:2014 §16. The reported parylene IDEs have similar dimensions and design as the PI IDEs in the current study, further justifying these failure criteria. Fig. 4b displays the impedance and phase of each IDE at 1-Hz as a function of time. By the end of the accelerated aging test, the impedances of all IDEs are still an order of magnitude larger than the metric (10^8^ Ω), and the phase of each IDE device remains close to -90°, indicating capacitive behavior. Therefore, all IDE devices survived in the accelerated aging test at 57 °C and exhibit an equivalent soaking lifetime of ≥ 336 days at 37°C.

To investigate the variation of the PI insulation with soaking time, equivalent circuit models (ECMs) were developed for the IDEs and applied to data from the beginning and end of the accelerated aging test, respectively. As no device failure was observed in the test, ECMs are very similar for all devices at both beginning and end of the test, and consist of solution resistance (*R*_*s*_) in series with the parallel combination of the PI layer resistance (*R*_*L*_) and the constant phase element (CPE), commonly known as a Randles circuit. CPE is expressed by the following equation.

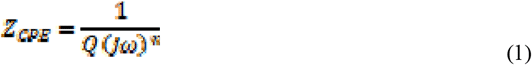

Where *Q* is a measure of the magnitude of CPE and associated with the capacitance, ω represents the angular frequency, and *n* is an exponential term that represents non-ideal behavior at electrode surface. When *n*=1, the CPE has ideal capacitor character, while *n*=0, the CPE behaves purely as a resistor.

The ECM fits the Bode and Nyquist plots reasonably well for all IDE devices (Fig. 4c-f, Fig. S10-12) with the 3-electrode and 2-electrode configurations, and fits the experimental data at the end of the test where the impedances are lower, much better than that at the beginning. The higher impedances from the 2-electrode configuration at the beginning of the study are closer to the measurement limits of the EIS instrument, which might impact modeling fits. The parameters of the ECM for all IDE devices at different time points, are listed in Table 3 and 4, corresponding to 3-electrode and 2-electrode configurataions, respectively. In both Table 3 and 4, there is no statical difference in *R*_*s*_ between day 0 and day 336, indicating the stability of the PBS in the accelerated aging test. In 3-electrode configuration, *R*_*L*_ remarkably reduced by 85%, from 129.1 ± 18.6 GΩ to 19.3 ± 2.2 GΩ (*p* < 0.05). Similarly, in 2-electrode configuration, *R*_*L*_ significantly decreased by 84% as well. Care must be used to interpret R_L_, as the data used to fit these values is limited and from values that are relatively high for the instrument. Consistent with previous reports, decreases in impedance and increases in phase angle at low frequencies are the most sensitive to encapsulation degradation. On the contrary, by the end of the test, |*Q*| rose by 66% for both 3-electrode and 2-electrode configurations. As water has higher relative permittivity (□_r-water_ = 78) than that of PI (□_r-PI_ = 3.4), water adsorption, absorption and penetration are thought to significantly increase the relative permittivity of the PI encapsulation in PBS solution [23], and thus result in the increased capacitance for IDEs. In addition to the relative permittivity, another potential mechanism is to generate more pathways from PBS solution to metal trace due to the degradation of PI insulating layers. More characterization information would be required to distinguish these mechanisms.

**Table 3.** Parameters of the ECM for IDE devices with the 3-electrode configuration (day 0 and day 336). * *p*<0.05.

| 3-electrode configuration | Day 0 (n=4, mean ±SD) | Day 336 (n=4, mean ±SD) |
| --- | --- | --- |
| $R_s$ ($\Omega$) | 256.6 ± 22.5 | 231.9 ± 28 |
| $R_L$ (GΩ) | 129.1 ± 18.6 | 19.3 ± 2.2 * |
| $n$ | 0.9902 ± 0.0063 | 0.9885 ± 0.0031 |
| $Q$ (pF) | 107.0 ± 4.7 | 177.5 ± 30.3 * |

**Table 4.** Parameters of the ECM for IDE devices with the 2-electrode configuration (day 0 and day 336). * *p*<0.05.

| 2-electrode configuration | Day 0 (n=4, mean ±SD) | Day 336 (n=4, mean ±SD) |
| --- | --- | --- |
| $R_s$ ($\Omega$) | 566.0 ± 28.7 | 598.0 ± 43.6 |
| $R_L$ (GΩ) | 233.0 ± 12.1 | 36.3 ± 3.4 * |
| $n$ | 0.9848 ± 0.0059 | 0.9858 ± 0.0024 |
| $Q$ (pF) | 50.1 ± 3.1 | 83.2 ± 0.7 * |

The electrical properties of *Flex* electrodes (n=24) are presented in Fig. 5, including the Bode plots for impedance magnitude and phase, CV plots, and CIC values at 3 common pulse widths. The impedance magnitude and phase of electrodes have a narrow distibution between electrodes at each frequency point in the Bode plot, due to the precise and repeatable micro-fabrication process. The 1-Hz and 10^3^-Hz impedance are 10557 ± 774 Ω and 443 ± 32 Ω, respectively, while the phase at 1 Hz and 10^3^ Hz are and -75.8 ± 1.2° and - 4. 4 ± 0.7°, respectively (Fig. S13a-d, Supplementary materials). The low impedance at 10^3^ Hz benefits both recording and stimulation of neural signals.

**Figure 5.**
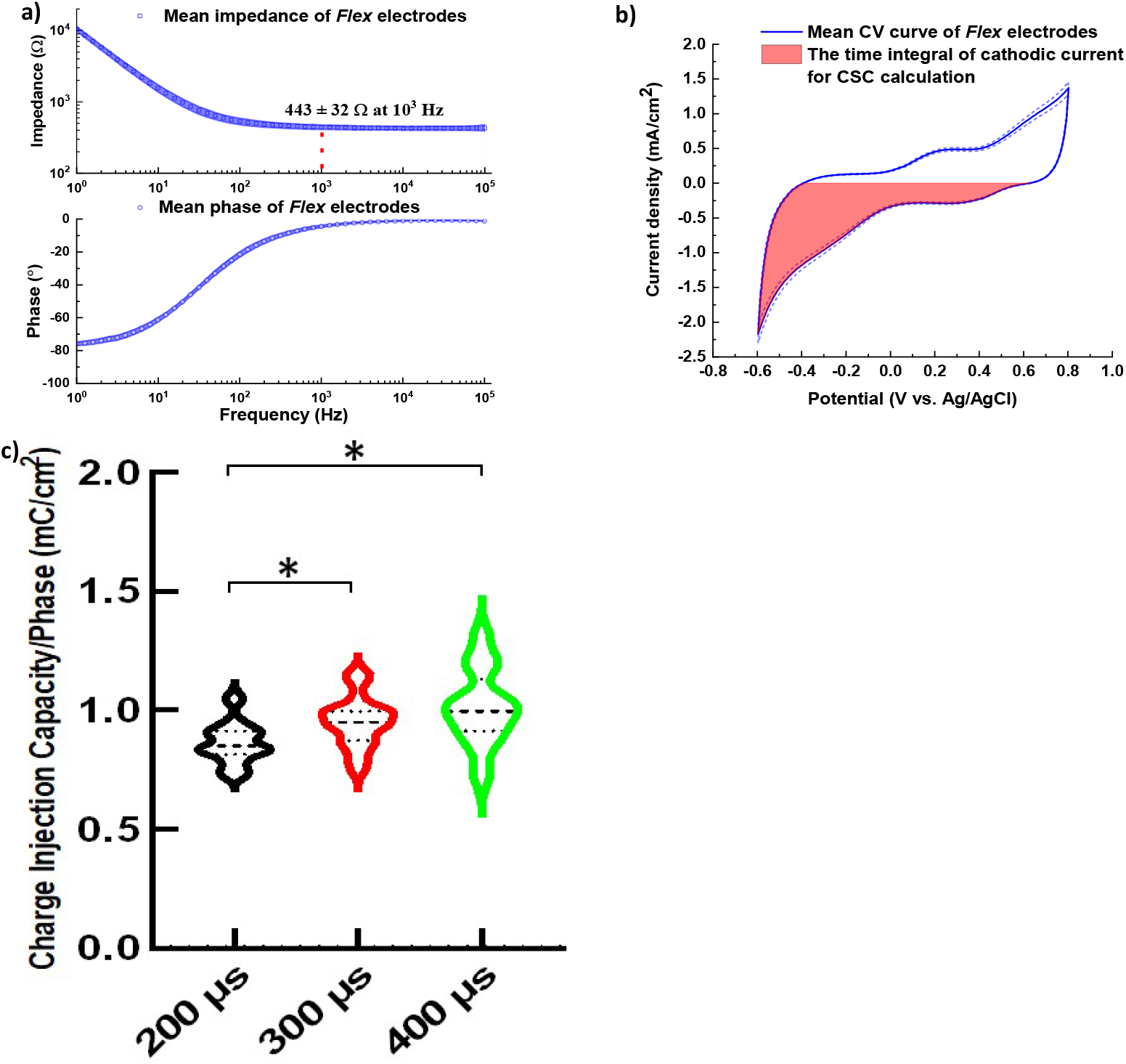
Electrochemical properties of *Flex* electrodes (n=24). a) Bode plot of *Flex* electrodes, the shaded area stands for the standard deviation (SD), (b) the mean CV curve of *Flex* electrodes, the dot lines stand for the SD of current density, while the shaded area refers to the calculation of CSC, (c) CIC of *Flex* electrodes at different pulse width, * *p* < 0.05.

The CV curves of *Flex* electrodes have tight distribution as well, and do not show strong redox reaction peaks within the water window (Fig. 5b). The CSC of the electrodes was calculated to be 12.76 ± 0.58 (mC/cm^2^). The violin plots of CIC at different pulse width is shown in Fig. 5c, and CIC increases with the pulse width from 0.86 ± 0.09 mC/cm^2^ at 200 µs to 1.01 ± 0.16 mC/cm^2^ at 400 µs, but there is no statistical differece in CIC between 300 µs and 400 µs.

The stability of IrO_x_ electrode sites on *Flex* electrodes was evaluated by the most aggressive stimulation test (Stim-Stab test) so far. In the Stim-Stab test, n=6 *Flex* electrodes were randomly and evenly divided into two groups stimulated 12 mA and 16 mA, respectively (n=3 for each group). Channel 1 of each electrode was unstimulated, while the charge-balanced, cathodic leading, biphasic pulses were applied to channel 2. After the Stim-Stab test, all unstimulated (control) and stimulated electrodes were evaulated using optical microscopy and electrically with potentiostatic and VT measurements. The representative results are summerized in Table 5 and Fig. 6, while Fig. S14-33 (supplementary materials) shows all results for each unstimulated (control) and stimulated electrodes. No unstimulated electrodes demonstrated visible damage to the electrode sites (Fig. S16, 19, 22, 25, 28, 31). For unstimulated channels tested at 12 mA, except the 1-Hz impedance, there is statistical difference in impedance and phase before and after the Stim-Stab test (Fig. S14, supplementary materials). For example, 10^3^-Hz impedance and phase significantly reduced after the test (17.3% and 32.7%, respectively), and 1-Hz phase increases due to the test (3.9%, *p* < 0.05). On the contrary, there is no statistical difference in impedance and phase for unstimulated channels tested at 16 mA before and after the Stim-Stab test. The impedance decrease occurs for the resistive component of the impedance spectra. Although the CSC remarkably rose in both groups of the unstimulated channels after the Stim-Stab test (32.5% and 22.3% for unstimulated channels at 12 mA and 16 mA, respectively, *p* < 0.05, Fig. S14g), CIC did not significantly change for all unstimulated channels (Fig S14h and i).

**Table 5.** Representative results from the Stim-Stab test for unstimulated and stimulated electrode channels.

| Sample | Stimulation current (mA) | Electrochemical properties |  |  |  |  | Optical damage after stimulation | Category |
| --- | --- | --- | --- | --- | --- | --- | --- | --- |
| | | Max variation in the real-time VT ($E_{ac}$) | Max variation in $E_{mc}$ | $\Delta$ Impedance at 1 kHz | $\Delta$ CSC | $\Delta$ CIC at 400 μs | | |
| Electrode 1 channel 1 | Unstimulated | N.A. | N.A. | -19.4 % | 16.0 % | 11.3 % | No | Unstimulated |
| Electrode 1 channel 2 | 12 | 6.7 % | 46.8 % | -13.6 % | 17.1 % | 39.1 % | Minor: grade 2 (optical damage: 3.75 %) | Mixed |
| Electrode 2 channel 1 | Unstimulated | N.A. | N.A. | -17.6 % | 18.1 % | -34.4 % | No | Unstimulated |
| Electrode 2 channel 2 | 12 | 7.4 % | 49.5 % | -17.8 % | 24.6 % | -35.3 % | Medium: grade 3 (optical damage: 5.32 %) | Mixed |
| Electrode 3 channel 1 | Unstimulated | N.A. | N.A. | -14.6 % | 63.7 % | -38.3 % | No | Unstimulated |
| Electrode 3 channel 2 | 12 | 8.4 % | 45.9 % | 305.1 % | -83.0 % | -33.0 % | Serious: grade 4 (optical damage: 18.29 %) | Fully Failed |
| Electrode 4 channel 1 | Unstimulated | N.A. | N.A. | -11.7 % | 23.0 % | 17.1 % | No | Unstimulated |
| Electrode 4 channel 2 | 16 | 7.1 % | 30.2 % | -13.9 % | 26.2 % | -9.1 % | Minor: grade 2 (optical damage: 3.58 %) | Mixed |
| Electrode 5 channel 1 | Unstimulated | N.A. | N.A. | 10.6% | 27.9 % | -16.7 % | No | Unstimulated |
| Electrode 5 channel 2 | 16 | 7.2 % | 27.1 % | -0.9% | 35.4 % | -12.0 % | No obvious: grade 1, (optical damage, « 1%) | Survived |
| Electrode 6 channel 1 | Unstimulated | N.A. | N.A. | -0.9 % | 16.4 % | -32.8 % | No | Unstimulated |
| Electrode 6 channel 2 | 16 | 22.8 % | 37.2 % | 2.0 % | 13.1 % | -28.0 % | Serious: grade 4 (optical damage: 18.74 %) | Mixed |

**Figure 6.**
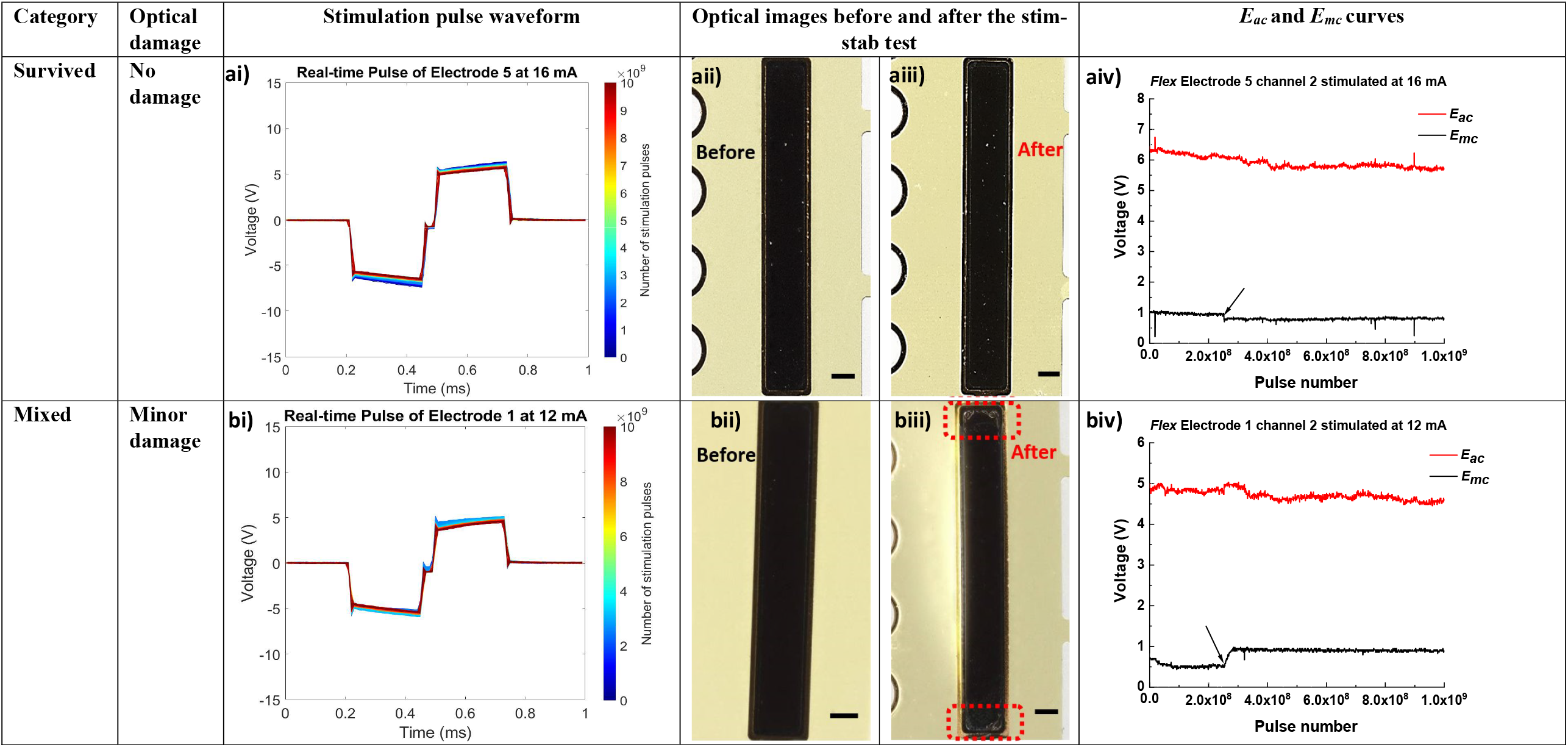
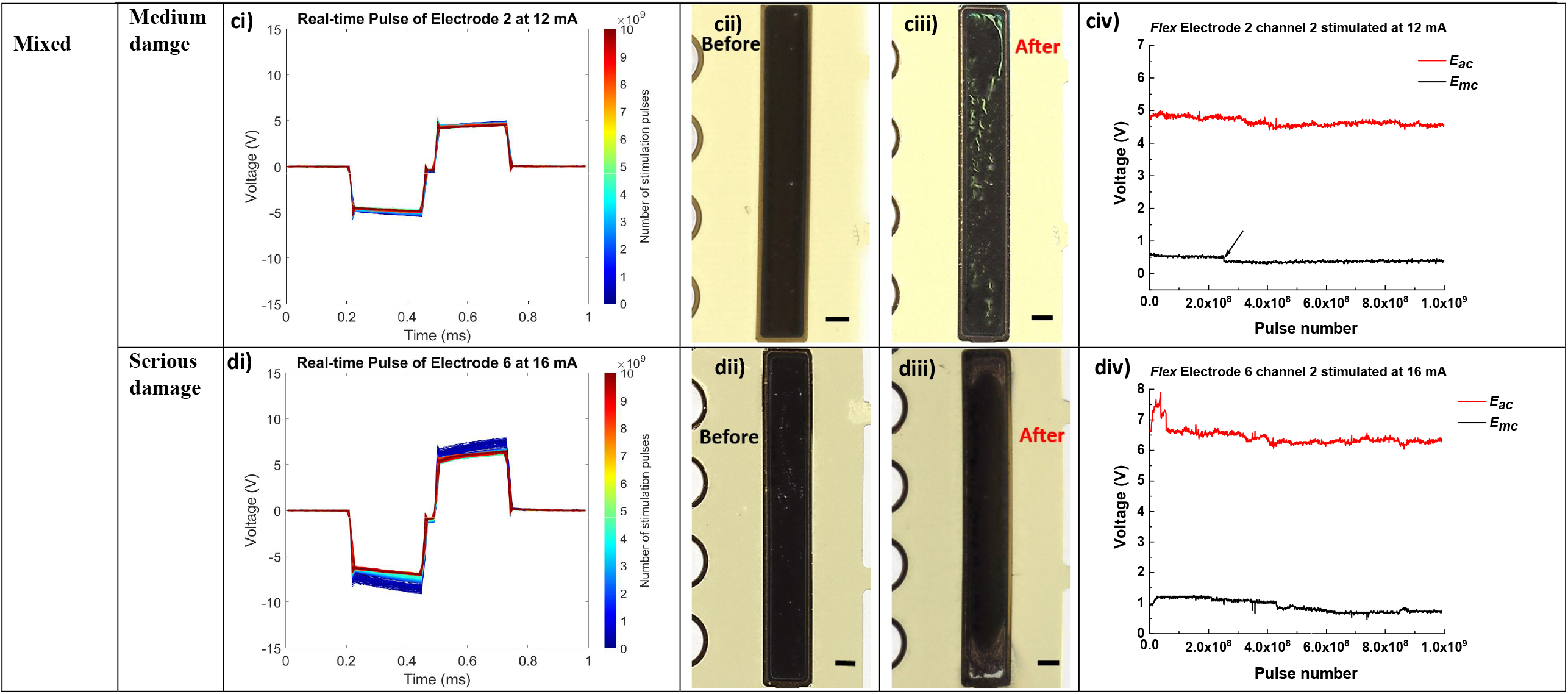
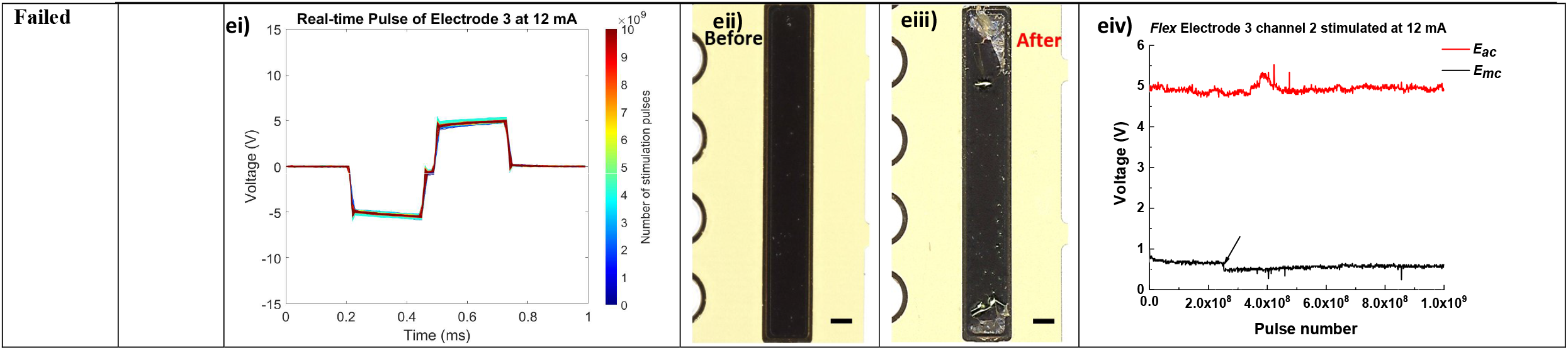
Stimulation and stability (Stim-Stab) test of the *Flex* electrodes. ai) real-time VT waveform of the survided electrode channel (electrode 5 channel 2 stimulated at 16 mA) as a function of the pulse number, aii) optical image of the survived electrode channel before the Stim-Stab test, aiii) optical image of the survived electrode channel after the Stim-Stab test, aiv) *E*_*ac*_ and *E*_*mc*_ curves of the survived electrode channel as a function of the pulse number, bi) real-time VT waveform of the mixed electrode channel (electrode 1 channel 2 stimulated at 12 mA) with minor damage as a function of the pulse number, bii) optical image of the mixed electrode channel with minor optical damage before the Stim-Stab test, biii) optical image of the mixed electrode channel with minor damage after the Stim-Stab test, biv) *E*_*ac*_ and *E*_*mc*_ curves of the mixed electrode channel with minor damage as a function of the pulse number, ci) real-time VT waveform of the mixed electrode channel (electrode 2 channel 2 stimulated at 12 mA) with midium damage as a function of the pulse number, cii) optical image of the mixed electrode channel with midium damage before the Stim-Stab test, ciii) optical image of the mixed electrode channel with midium damage after the Stim-Stab test, civ) *E*_*ac*_ and *E*_*mc*_ curves of the mixed electrode channel with midium damage as a function of the pulse number, di) real-time VT waveform of the mixed electrode channel (electrode 6 channel 2 stimulated at 16 mA) with serious damage as a function of the pulse number, dii) optical image of the mixed electrode channel with serious damage before the Stim-Stab test, diii) optical image of the mixed electrode channel with serious damage after the Stim-Stab test, div) *E*_*ac*_ and *E*_*mc*_ curves of the mixed electrode channel with serious damage as a function of the pulse number, ei) real-time VT waveform of the failed electrode channel (electrode 3 channel 2 stimulated at 12 mA) as a function of the pulse number, eii) optical image of the failed electrode channel before the Stim-Stab test, eiii) optical image of the failed electrode channel (electrode 3 channel 2 stimulated at 12 mA) after the Stim-Stab test, eiv) *E*_*ac*_ and *E*_*mc*_ curves of the failed electrode channel as a function of the pulse number (the arrows in Fig. 6aiv, biv, civ and eiv indicate a power failure, resulting in the instrumentation being operated under backup power). Scale bar=100 µm.

For stimulated channels in Table 5, the variation in the real-time VT waveform during the Stim-Stab test can be determined by the difference between maximum *E*_*ac*_ and mean *E*_*ac*_ during the Stim-Stab, and the variation in *E*_*mc*_ was calculated in the same manner. According to the damage areas calculated by ImageJ in the optical images, the stimulated electrode channels were classifed into 4 categories from 1-4. An electrode with no obvious damage (damage area < 1 %) is 1, electrode with minor damage (1 % ≤ damage area < 5 %) is graded a 2, electrode with medium damage (5 % ≤ damage area ≤ 10 %) is 3, and electrode with serious damage (damage area > 10 %) is 4. Meanwhile, the electrical properties of stimulated channels (impedance, CSC, and CIC) were evaluated to determine if these properties changed, and how these correlated with the real-time VT waveforms collected during the Stim-Stab test. Hence, three metrics including optical images, electrical properties, and the variation in real-time VT waveform, are employed to thoroughly identify failures, lifetimes, and to help determine failure mechanisms for electrodes. For example, electrode 5 channel 2 that has no obviuos optical damage («1 %) and variation in real-time VT waveform (7.2 %), remains functional from the perspective of electrical properties, and therefore is determined to survive throughout the Stim-Stab. Four stimulated channels were found to have damage by optical microscopy, but maintained their electrical function after the Stim-Stab test. In additon, only small changes were present in their real-time VT waveforms during the test, making outcomes for these electrodes mixed. Electrode 3 channel 2 demonstrated failure from optical inspection, a modest but relatively large variation (8.4%) in the real-time VT waveform during the Stim-Stab test, and the degradation of electrical properties (increase in impedance, and reduction in CSC and CIC), and was categorized as failed.

In Table 5, compared to those of other stimulated channels, the real-time VT waveforms of the stimulated channels with serious optical damage (electrode 5 channel 2, and electrode 3 channel 2) have higher variation (22.8% and 8.4%, respectively) during the Stim-Stab test. Similarly, except these two stimulated channels, all other stimulated channels exhibited lower impedance at 1 kHz after the Stim-Stab. Interestingly, except the failed channel, the CSC of other stimulated channels increased after the Stim-Stab, while CIC decreased for these stimulated channels. The substaintial increase in CSC and the significant decreased CIC are consistent with previous observations for Ir-based electrodes [15], [35], and are often referred to as activation. The formation of IrO_x_ by electrochemical activation of Ir is one putative mechanism behind the phenomena. Surprisingly, one electrode (electrode 3 channel 2) stimulated at 12 mA entirely failed due to dramtic increase in impedance (305.1%) at 1 kHz, reduced CSC (−83%) and CIC (−33%), and demonstrated catastrophic failure in optical imaging. Comparatively, another electrode (electrode 5 channel 2) stimulated at the higher current of 16 mA had minimal degradation from optical microscopy and electrochemical characterization. Although narrow distrbution in electrical properties and stable electrode quality arising from MEMS technology are shown in Fig. 5, these results suggest the non-equilibrium and harsh conditions (high-frequncy stimulation, large stimulation current density, and large number of stimulation pulses) does provide an important differentiation between electrodes. Additional Stim-Stab studies are being performed to generate sufficient population for statistical analysis and failure modes and effects analysis (FMEA) studies to gain more in-depth insight into the lifetime and failure mechanisms for the IrO_x_ electrodes.

Fig. S15 (supplementary materials) shows the real-time VT waveform of the 1 kΩ resistor as the function of the stimulation cycle. Due to its pure resistive behavior, the expected lack of polarization was observed between the cathodic and anodic phases, and each pulse waveform overlaped well from the beginning to end of the Stim-Stab test. As expected, the entire voltage drop was *E*_*ac*_ for the 1 kΩ resistor, and E_mc_ was 0 since no polarization occurs as shown in (Fig. S15a) and (Fig. S15b). The real-time VT waveform of the survived electrode channel (electrode 5 channel 2) as the function of the stimulation cycle is shown in Fig. 6ai, and has a variation of 7.2% (in Table 5) with the stimulation cycles. Similar to those of unstimulated channels, there is no evident damage on the electrode site of the survived electrode channel after the Stim-Stab test (Fig. 6aii and aiii). As 10^3^ Hz impedance of the survived electrode is 399 Ω before the test, under the stimulation current of 16 mA, the *E*_*ac*_ curve started from around 6.4V and gradually declined to 5.7 V at the end of the Stim-Stab test, corresponding the varation in the real-time VT waveform (Fig. 6ai) and indicating the reduced impedance due to the Stim-Stab test. The drop in the *E*_*mc*_ curve indicated by the star marker was resulted from the instrumentation being operated under backup power during a power failure. Except the decrease resulting from the power failure, the *E*_*mc*_ curves are relatively flat and stable not only for the survived electrode channel, but also for the failed electrode channel and mixed electrode channels.

Fig. 6bi displays the real-time VT waveforms for mixed electrodes with electrode degradation scores of 2. After the Stim-Stab test, for the mixed electrode with a damage score of 2, degradation of the IrO_x_ is oberved at the corners of the electrode site (Fig. 6biii). For the electrode that had a damage score of 3 but without evidence of electrode failure from electrical properties and VT waveforms (Fig. 6ci), the wrinkle-like features were present on the surface of the electrode site (Fig. 6ciii). More advanced analysis by electron microscopy and analytical techniques is needed to better understand the damage mechanisms. Similarly, there is no evident peaks in the *E*_*ac*_ curves of the mixed electrodes with the damage score of 2 or 3, and the *E*_*ac*_ slightly reduced at the end of the Stim-Stab test.

The real-time VT waveforms of the seriously damaged eletrode channel with mixed outcomes, and the failed electrode channel are shown in Fig. 6di and ei, respectively. Although the VT waveform of the electrode channel with a damage score of 4 has larger variaion (22.8%) under the stimulation current of 16 mA, the electrode channel still remained functional at the end of the Stim-Stab test, due to the silight incresase in impedance (2%) and CSC (13.1%). Discoloration at the upper corners appears to have been caused by the delimination of IrO_x_ along with damage to lower layers of the electrode stack (Fig. 6diii). For the failed electrode channel, it is evident that serious delimination of the IrO_x_ top layer occurred at the upper and bottom electrode site (Fig. 6eiii). Corresponding to the large variation in real-time VT waveforms, there are broad peaks in the *E*_*ac*_ curves of the mixed electrode channel with serious damage and failed electrode channel (Fig. 6div and eiv, respectively). But it is not clear whether the electrode damage occurred at the moment when the peaks were present in the *E*_*ac*_ curves. Despite the fact that 10^3^-Hz impedance of the failed electrode channel rose by 3 times after the Stim-Stab test, the *E*_*ac*_ of the failed electrode channel did not reach around 15V at the end of the Stim-Stab test, indicating that the delaminated IrO_x_ top layer still weakly and physically attached to the electrode site of the failed electrode channel. It is the cleaning process after the Stim-Stab by deionized water that removed the delaminated IrO_x_, resulting in the substantial increase in the impedance for the failed electrode channel.

To further explore the possible failure mechanism of the stimulated electrode channels, ECMs were created to fit the impedance spectra of unstimulated and stimulated electrode channels. Fig. 7 depicts the ECM fit for the impedance spectra of representative survivied, mixed and failed electrode channels before and after the Stim-Stab test. For unstimulated electrode channels, the survived electrode channel and other mixed electrode channels, the ECM based on classic Randles circuit fits well for their Bode and Nyquist plots before and after the Stim-Stab (Fig. 7a, b, c and d). The ECM includes a CPE, shunted by a charge transfer resistance R_ct_, together in series with the solution resistance R_s_. Similar to the CPE in the ECM for IDEs, the CPE for these electrode channels is an approximation to an ideal interfacial capacitance resulting from a double layer that forms on the surface of the IrO_x_ electrode site, when the *Flex* electrodes are immersed in an electrolyte. The deviation from the ideal pure capacitance was reported by the surface roughness, defects or even protein adsorption on the electrode (that is not a factor in this *in vitro* test) [36]. The CPE can be expressed by the same equation in eq (1). R_ct_ is a resistance of a faradaic reaction against the process of electron tranfer from one phase (electrode) to another (electrolyte). The fitting parameters are shown in Table S1 (supplementary materials). For unstimulated electrode channels, the survived electrode channel and the mixed electrode channel with serious damge, their impedance spectra change little between before and after the Stim-Stab test, resulting in modest changes to the fitting paremeters. For the failed electrode channel, however, not only the fitting parameters, but also the ECM is significantly different from others. After the Stim-Stab test, Warburg impedance (Z_warburg_), accounting for diffusion of ions across the interface, was presented in the ECM of the failed electrode channel, revealing that delamination of IrO_x_ or corroion of underlying electrode materials may have occured.

**Figure 7.**
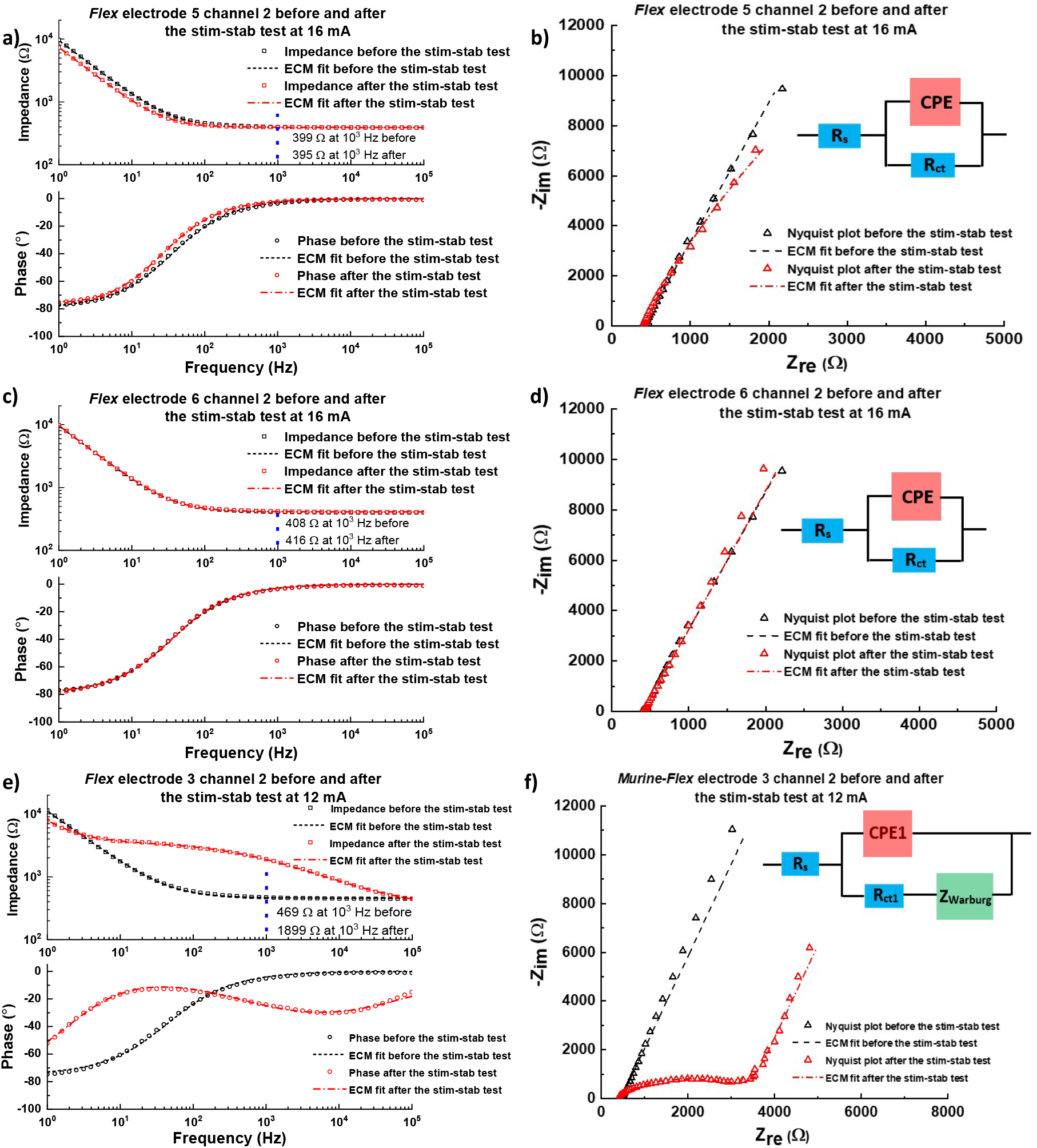
Equivalent circuit model fit for representative stimulated electrode channels before and after the Stim-Stab test. a) ECM fit for Bode plots of the survived electrode channel (electrode 5 channel 2), b) ECM fit for Nyquist plots of the survived electrode channel, c) ECM fit for Bode plots of the mixed electrode channel with serious damage (electrode 6 channel 2), d) ECM fit for Nyquist plots of the survived electrode channel, e) ECM fit for Bode plots of the failed electrode channel (electrode 3 channel 2), f) ECM fit for Nyquist plots of the failed electrode channel before and after the Stim-Stab test.

The efficacy of *Flex* electrodes were evaluated by stimulating the vagus nerve of mouse models (n=6), with nerve activation evaluated using heart rate and respiratory rate as biomarkers to judge the efficacy of stimulation. VNS has been widely observed to evoke a slowing heart rate (bradycardia) and respiratory rate (apnea) [37]–[39]; therefore, both the heart rate and respiratory rate were monitored as a function of stimulation current to validate efficacy of the electrodes with acute mouse preparations. The electrodes were place around the cervical vagus nerve, with sutures in the suture holes to retain the electrode on the nerve as shown in Fig. 8c. The vagus nerve was stimulated with currents from 100 to 500 µA in increments of 100 µA, while the EKG and respiratory signals were recorded (Fig. 8a). VNS 1 and VNS 2 for Fig. 8a are labled in red boxes.

**Figure 8.**
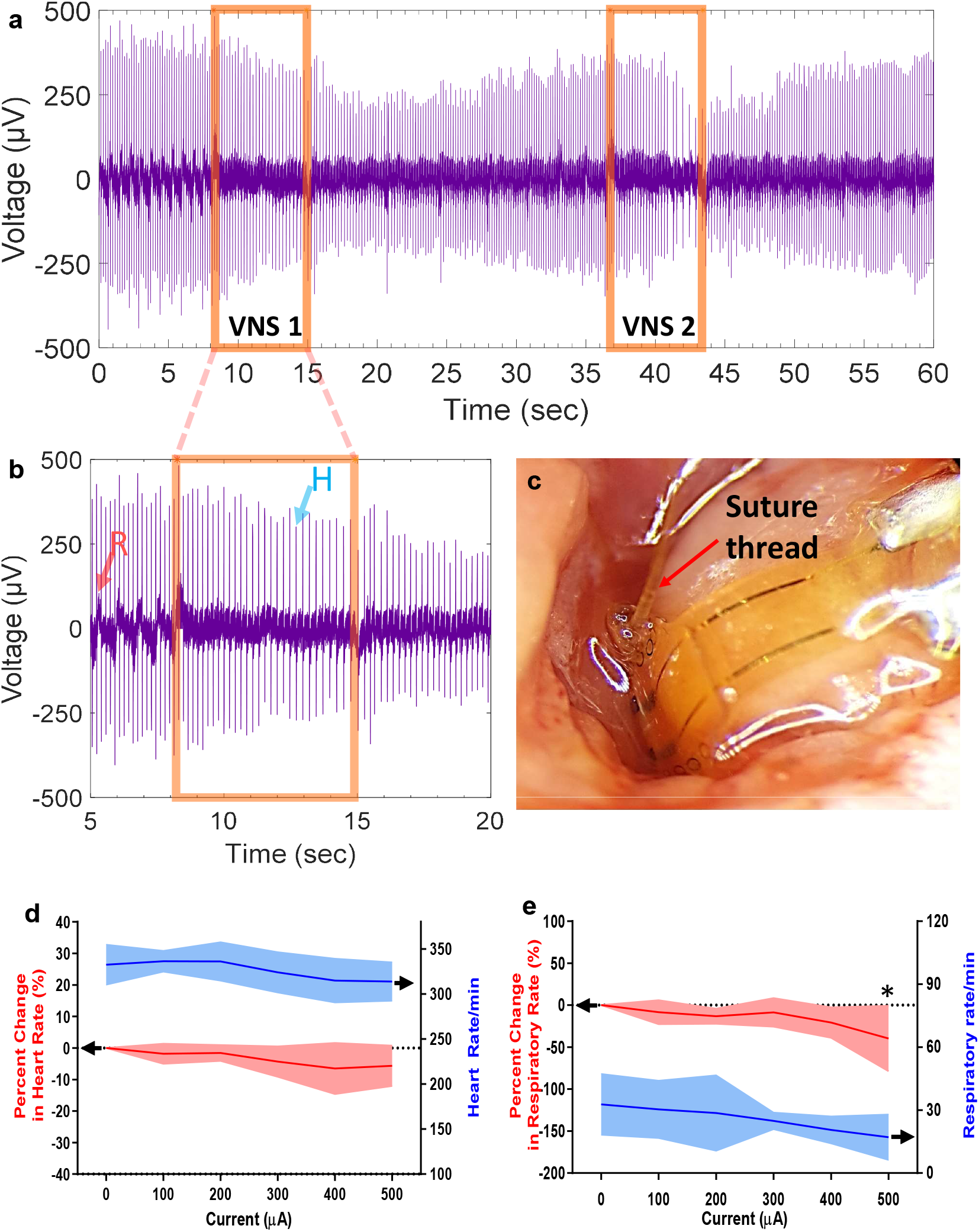
Acute mouse vagus nerve stimulation using *Flex* electrodes. a) typical ECG waveform in the acute mouse VNS (the red boxes indicate the duration of the VNS 1 and VNS 2 for Fig. 8a), b) zoom-in figure of VNS 1 for Fig. 8a (the peaks corresponding to respiratory and heart beat are labled as R and H, respectively), c) optical image of the *Flex* electrode used for acute mouse VNS (the suture thread to fix the *Flex* electrode is indicated by the red arrow), d) heart rate and percentage change in heart rate as a function of the stimulation current, e) respiratory rate and percentage change in respiratory rate as a function of the stimulation current.

In the zoom-in figure of VNS 1 (Fig. 8b), small and broad peaks from the dark purple curve correspond to the respiratory (which is labled by R), and the sharp peaks with the amplitude of ∼ 400 mV result from heart beats (which is marked by H). During the VNS 1, the number of the respiratory peaks decreased, indicating the induced apnea occurred. In addition, the time between two heart beat peaks slightly increased, revealing the heart rate slowed down. The phenomenon is more evident at the end of VNS 2 in Fig. 8a.

The heart rate and percentage change in heart rate during stimulation compared to baseline (ΔHR) were extracted, and both of them are plotted as a function of stimulation current in Fig. 8d. Likewise, the respiratory rate and percentage change in respiratory rate compared to baseline (ΔRR) are shown in Fig. 8e, as a function of stimulation current. Interestingly, the decreased heart rate was not found to be statistically different at each stimulation condition (*p* > 0.05), while the trend of decreasing respiratory rate reached statistical significance at the 500 µA condition (*p* < 0.05), demonstrating the efficacy of the *Flex* electrodes.

## 4. Discussion

In this study, a flexible IrO_x_ electrode technology was specifically developed as a mouse vagus nerve interface. It is essential that the flexible IrO_x_ electrode remain functional in the harsh *in vivo* environment after long-term neural stimuation and recording. Hence, the lifetime evaluation of the PI insulating layers and IrO_x_ electrode are of paramount importance. However, the electrochemical properties and lifetime for electrodes fabricated from these materials need further exploration, especially regarding the stability of IrO_x_ electrodes on polymeric substrates for these to be used in truly long-term bi-directional interfaces.

Consistent with methods used in previous studies on Parylene C and Al_2_O_3_-parylene C, IDEs were used to assess the lifetime for PI insulating layers in this investigation. For IDEs with Parylene C encapsulation, the mean time to failure was found to be ∼ 8 months, at a similar thickness (6 µm). In this study, IDEs using the same design, but fabricated with 7 µm-thick PI encapsulation entirely survived in the accelerated aging test and have an equivalent lifetime of ≥ 336 days (∼ 11 months) at 37 °C, which is comparable to that of Parylene C. The optimized and reproduciable micro-fabrication processes developed in this study are contributed to the enhanced lifetime of bottom and top PI insulating layers, compared to the state-of-the-art. Additional accelerated aging tests at a higher temperature (67 °C instead of 57 °C) are also underway, and will be used to improve the statistical accuracy, and also continue to explore failure mechanisms.

Electrical stimulation is a critical aspect of neural technologies to implement bioelectronic medicines, restore senses, and modulate neural states, making stimulation stability a critical need for bi-directional neural interfaces. The stimulation tests to evaluate the stability of the electrode materials that have been reported are listed in Table 6 [40]– [44]. Although electrochemical properties of PEDOT coatings on PtIr substrate were stable at a high charge densities up to 2.89 mC/cm^2^, the current pulsing in the stimulation test only lasted for 24 hours at a frequency of 100 Hz. PEDOT performance in long-term testing is still unknown [40]. Moreover, images of the PEDOT coating after the stimulation test were not provided. Mechanical delaminaiton from substrates continues to be concerns for chronic *in vivo* use of the material [45]–[47].

**Table 6.** State-of-the-art for the eletrical stimulation test

| Electrode materials | Stimulation current (mA) | Current density (A/cm<sup>2</sup>) | Pulse width/phase (μs) | Charge density (mC/cm<sup>2</sup>/phase) | Stimulation pulse (10<sup>6</sup>) | Stimulation time (hour) | Stimulation frequency (Hz) | Total delivered negative charge (C/cm<sup>2</sup>) | Literature |
| --- | --- | --- | --- | --- | --- | --- | --- | --- | --- |
| PEDOT-PSS | 0.13 | 2.89 | 1000 | 2.89 | 8.64 | 24 | 100 | $24.97 \times 10^3$ | [40] |
| Glassy carbon | 0.45 | 0.625 | 400 | 0.25 | 3600 | 1000 | 1000 | $900 \times 10^3$ | [41] |
| Diamond on Pt foil | ~1.8 | ~0.23 | 400 | 0.092 | 21.6 | 120 | 50 | $1.98 \times 10^3$ | [42] |
| Electrodeposited Pt-Ir on Pt | 2.0 | 1.43 | 140 | 0.2 | 544 | 504 | 300 | $108.8 \times 10^3$ | [43] |
| Sputtered IrO<sub>x</sub> in water vapor | $40 \times 10^{-3}$ | 2 * | 200 | 0.4 | 1000 | 1388.9 | 200 | $400 \times 10^3$ | [44] |
| Sputtered IrO<sub>x</sub> in this study | 16.0 | 6.64 | 240 | 1.59 | 1000 | 277.78 | 1000 | $1590 \times 10^3$ | This work |

Glassy carbon microelectrodes with Durimide® encapsulation were subjected to 3.5 billion cycles of current pulsing at the charge density of 0.25 mC/cm^2^/phase [41]. Although there was no obvious change in the surface morphology characterized by SEM and atomic force microscopy (AFM) after the stimulation test, the impedance spectra and CV curves of glassy carbon microelectrodes evidently changed even after 200 million cycles of current pulsing. Moreover, the delamination of graphene on a Pt substrate, and subsequent Pt substrate corrosion have been reported at the stimulation charge density of 0.2 mC/cm^2^/phase, indicating carbon-based electrode materials are recognized for their susceptibility to degradation by electrical stimulation, particularly at intensities and durations required for chronic interfaces [43].

Recently, the stability of SIROF deposited using water vapor as part of the reactive sputtering plasma environment were investigated by coating the IrO_x_ films on the tips of a Utah electrode array [44]. For the estimated tip area of 2.0 × 10^−5^ cm^2^ [48], the SIROF deposited using water vapor showed minial changes in terms of surface morphology and electrical properties, after a 10^9^ current pulsing stimulation at 0.4 mC/cm^2^/phase. But the stability of sputtered IrO_x_ films on PI substrates under electrical stimulation has rarely been investigated for neural interfaces.

Compared to these state-of-the-art, the Stim-Stab test developed in this study generates a harsher environment to evaluate the stability of the IrO_x_ electrode, because of the high charge density/phase (1.59 mC/cm^2^/phase), large number of stimulation pulse (10^9^), and high stimulation frequency (10^3^ Hz). Accordingly, the total delivered negative charge is as high as 1.59 ×10^6^ C/cm^2^, and at least 1.5-fold higher than the-state-of-the-art. These Stim-Stab studies support collecting data to enable optimization of electrode and insulation materials for the electrodes. Additioanlly, the data can help initiate the process of developing more quantitative models to predict lifetime. More importantly, rigorous metrics to determine the electrode failure were built on the thorough optical and electrical characterization, including optical imaging, real-time VT waveform, EIS, CSC and CIC. Because we found that any one from these three metrics alone cannot accurately reflect the lifetime of the electrode materials. For example, in this study, the electrode (electrode 6 channel 2) with serious damage (damage score of 4) still remained electrically functional, and its impedance spectra before and after the Stim-Stab almost overlaped to each other (Fig. S32).

Real-time VT waveform during the Stim-Stab test enables monitoring for changes of *E*_*ac*_ and *E*_*mc*_. A large voltage increase in the real-time VT waveform can be one of metrics to detect electrode failure. An important aspect of monitoring the VT during the Stim-Stab test is the ability to characterize the failure lifetime within the study endpoints, which can greatly improve statistical failure analysis and lifetime analysis for electrodes. Though we did not observe the large increases in voltage associated with failure for the electrodes in this study, voltage increases associated with electrode failure have been observed in heretofore unpublished Stim-Stab test data for Utah electrode arrays, which were corroborated by electron microscopy imaging. In the current study, *E*_*mc*_ curves were stable even for the electrode with serious damage and the electrode that was found to have failed electrically by EIS, CSC, and/or CIC characterization at the study endpoint. The exception was the passing increase in voltage transients associated with the time point when a power failure resulted in the instrumentation being operated under backup power. It is not clear whether the 3D structure, Si substrate, differences in the electrode metallization stack, electrode geometry, or other factors in the Utah electrode array that resulted in the expected large increase in voltage for the VT waveforms of failed electrodes. Such failed Utah electrode arrays were observed to have catastrophic damage with all metalization gone from the electrode tip, which was more extreme than electrode metalization failures we observed with *Flex* electrodes.

High-resolution optical microscopy using reflected and transmitted light is an effective and direct approach to identify electrode damage from a large-area field of view, while SEM and AFM can detect and provide more comprehensive materials characterization of damge in small scanning areas. In this studies, the category of the electrode damage is based on the ratio of damaged area in optical images. In future studies, deeper failure analysis to understand the failure mechanisms associated with the Stim-Stab paradigm will use techniques such as X-ray photoelectron spectroscopy (XPS), and inductively coupled plasma mass spectrometry (ICP-MS), to characterize the chemical states of IrO_x_, and metal ion concentration in the PBS solution, respectively. With these rigous metrics to determine the electrode failure reported here, half of stimulated electrode channels (3 out of 6) exhibit minor damage or less after the most aggressive Stim-Stab test so far, indicating the excellent stability of the IrO_x_.

More than a decade ago, IrO_x_ was proposed as the electrode material for flexible thin-film electrodes with PI insulation [49]–[51], and recently a PI based TIME electrode with 80-µm-diameter IrO_x_ contact sites was reported to deliver intraneural sensory feedback to an upper-limb amputee in a first-in-human study [27]. But there is a wide range of electrochemical properties variation for IrO_x_ thin films, depending on the difference in fabrication methods (electrochemical deposition, sputtering, electrochemical activation), fabrication parameters (the ratio of oxygen plasma, plasma species), electrode designs (thickness, electrode dimension, substrates), and testing parameters (pulse width, bias potential). As one of the critical parameters to evaluate the electrical properties of neural elecrodes, CIC refers to the charge quantity that polarizes the electrode interface to the potential for water reduction or oxidation, and therefore defines the maximum quantity of charge that can be safely injected or transferred to surrounding tissues by reversible processes. Lee *et al*. reported that the CIC of a SIROF on PI was 0.03 mC/cm^2^ at the VT pulse width of 1 ms for the flexible cuff electrode with an electrode area of 1.0 × 10^−2^ cm^2^ [52], while 3D Utah electrode array with tips coated by a SIROF showed a CIC of 2.0 mC/cm^2^ at 0.2 ms pulse [48]. Using 0.4 ms pulses, Cogan *et al*. determined the CIC of a SIROF on a planar PI test structure to be 1.9 mC/cm^2^ [49]. Therefore, the IrO_x_ used for the thin film electrode in this study not only is comparable to the state-of-the-art in terms of electrical properties (CIC = 1.01 ± 0.16 mC/cm^2^ at 400 µs), but also exhibits excellent stability demonstrated by the most aggressive Stim-Stab test so far.

## 5. Conclusions and future directions

Highly stable *Flex* IrO_x_ thin film electrodes with PI insulating layers were fabricated towards development of chronic vagus nerve interface, using cost-effective and scalable micro-fabrication processes. The cost of manufacturing and integration processes was broken down, and we determined that the technology could be a cost effective and scalable solution to interface small peripheral nerves. For acute and chronic vagus nerve interfaces, two reliable integration configurations were developed, respectively, and verified using electrical and mechanical testing paradigms. PI insulating layers evaluated by the accelerated aging test at 57 °C have an equivalent lifetime of ≥ 336 days at 37 °C. An aggressive electrical stimulation test (∼1.59 mC/cm^2^/phase, 10^3^ Hz, 10^9^ current pulses) was employed to assess the stability of IrO_x_ electrode sites, with rigorous metrics developed to identify and quantify the damage and failure for the electrodes. At the endpoints of our aggressive Stim-Stab paradigm, half of electrodes show minor damage or less, indicating their excellent stability. The efficacy of *Flex* electrodes was demonstrated by stimulation of the mouse vagus nerve, and evoking statistically significant apnea in mice at the stimulation current of 500 µA. In future studies, more advanced analysis will be conducted to better understand the underlying damage mechanisms during Stim-Stab testing. More comprehensive efficacy studies, and investigation of the electrochemical properties of the electrodes *in vivo* will be performed to validate the robust performance of the electrode as well. This knowledge will be used as the basis to drive engineering improvements to further improve the functional lifetime and performance of our electrodes.

## Acknowledgements

We thank Brian van Devener, Randy Polson, and Paulo Perez for providing instrumental help, guidance, and discussion, and Mahendar Ochani for providing help and guidance for animal surgical procedures performed as part of this work.

