## Supplementary figures and table for "High-Stability Polyimide-based Flexible Electrodes with IrO_x_ to Interface the Mouse Vagus Nerve"

### Supplemetary materials of the manuscript

**Caption list**

**Figure S1**. a) optical image of the setup to make the helical structure, b) optical image of the mould for the silicone encapsulation surrounding the helical Pt leads, the scale bar=2.0 cm.

**Figure S2**. Representative VT waveform of a *Flex* electrode in response to the current pulsing

**Figure S3**. Optical image of the integrated *Flex* electrode partially soaked in PBS for the Stim-Stab test (scale bar=10 mm).

**Figure S4.** Current waveforms generated by the stimulator for the Stim-Stab test.

**Figure S5.** Mechanical properties of the integrated electrodes for acute and chronic vagus nerve stimulation. a) the maximum load at the breakage for integrated acute configuration electrodes with and without epoxy reinforcement, b) the maximum extension at the breakage for integrated acute configuration electrodes with and without epoxy reinforcement (acute vagus nerve stimulation), c) the maximum load at the breakage for integrated chronic configuration electrodes with nonhelical and helical Pt leads, d) the maximum extension at the breakage for integrated chronic configuration electrodes with nonhelical and helical Pt leads.

**Figure S6**. a) Optical image of an integrated IDE for the accelerated aging test, scale bar=5.0mm, b) the current flow for 3-electrode configuration, c) the current flow for 2-electrode configuration.

**Figure S7.** Impedance spectra of the IDE 01-channel 02 device as a function of the soaking time. a) Bode plot of IDE 01-channel 02 device with 3-electrode configuration as a function of the soaking time, b) Nyquist plot of IDE 01-channel 02 device with 3-electrode configuration as a function of the soaking time, c) Bode plot of IDE 01-channel 02 device with 2-electrode configuration as a function of the soaking time, d) Nyquist plot of IDE 01-channel 02 device with 2-electrode configuration as a function of the soaking time.

**Figure S8.** Impedance spectra of the representative IDE device (IDE 02-channel 01) as a function of the soaking time. a) Bode plot of IDE 02-channel 01 device with the 3-electrode configuration as a function of the soaking time, b) Nyquist plot of IDE 02-channel 01 device with the 3-electrode configuration as a function of the soaking time, c) Bode plot of IDE 02-channel 01 device with the 2-electrode configuration as a function of the soaking time, d) Nyquist plot of IDE 02-channel 01 device with the 2-electrode configuration as a function of the soaking time.

**Figure S9.** Impedance spectra of the IDE 02-channel 02 device as a function of the soaking time. a) Bode plot of IDE 02-channel 02 device with 3-electrode configuration as a function of the soaking time, b) Nyquist plot of IDE 02-channel 02 device with 3-electrode configuration as a function of the soaking time, c) Bode plot of IDE 02-channel 02 device with 2-electrode configuration as a function of the soaking time, d) Nyquist plot of IDE 02-channel 02 device with 2-electrode configuration as a function of the soaking time.

**Figure S10.** Equivalent circuit model fit for the impedance spectra of IDE 01-channel 02. a) an equivalent circuit model fit for the Bode plot of IDE 01-channel 02 with the 3-electrode configuration, b) an equivalent circuit model fit for the Nyquist plot of IDE 01-channel 02 with the 3-electrode configuration, c) an equivalent circuit model fit for the Bode plot of IDE 01-channel 02 with the 2-electrode configuration, d) an equivalent circuit model fit for the Nyquist plot of IDE 01-channel 02 with the 2-electrode configuration.

**Figure S11.** Equivalent circuit model fit for the impedance spectra of IDE 02-channel 01. a) an equivalent circuit model fit for the Bode plot of IDE 02-channel 01 with the 3-electrode configuration, b) an equivalent circuit model fit for the Nyquist plot of IDE 02-channel 01 with the 3-electrode configuration, c) an equivalent circuit model fit for the Bode plot of IDE 02-channel 01 with the 2-electrode configuration, d) an equivalent circuit model fit for the Nyquist plot of IDE 02-channel 01 with the 2-electrode configuration.

**Figure S12.** Equivalent circuit model fit for the impedance spectra of IDE 02-channel 02. a) an equivalent circuit model fit for the Bode plot of IDE 02-channel 02 with the 3-electrode configuration, b) an equivalent circuit model fit for the Nyquist plot of IDE 02-channel 02 with the 3-electrode configuration, c) an equivalent circuit model fit for the Bode plot of IDE 02-channel 02 with the 2-electrode configuration, d) an equivalent circuit model fit for the Nyquist plot of IDE 02-channel 02 with the 2-electrode configuration.

**Figure S13.** Electrochemical properties of *Flex* electrodes (n=24). a) violin curve of the impedance at 1 Hz, b) violin curve of the impedance at $10^3$ Hz, c) violin curve of the phase at 1 Hz, d) violin curve of the phase at $10^3$ Hz, e) violin curve of the CSC.

**Figure S14**. Electrical properties of unstimulated electrode channels before and after the Stim-Stab test. a) Bode plots of unstimulated electrode channels before and after the Stim-Stab test at 12 mA (shaded areas indicate the standard deviation), b) Bode plots of unstimulated electrode channels before and after the Stim-Stab test at 16 mA (shaded areas indicate the standard deviation), c) 1-Hz impedance of unstimulated electrode channels before and after the Stim-Stab test, d) $10^3$-Hz impedance of unstimulated electrode channels before and after the Stim-Stab test, e) 1-Hz phase of unstimulated electrode channels before and after the Stim-Stab test, f) $10^3$-Hz phase of unstimulated electrode channels before and after the Stim-Stab test, g) CSC of unstimulated electrode channels before and after the Stim-Stab test, h) CIC of unstimulated electrode channels before and after the Stim-Stab test at 12 mA, i) CIC of unstimulated electrode channels before and after the Stim-Stab test at 16 mA.

**Figure S15.** a) Real-time stimulation pulse of the 1-kΩ resistor as a function of the pulse number, b) $E_{ac}$ and $E_{mc}$ curves of the 1-kΩ resistor as a function of the pulse number.

**Figure S16.** Optical image of *Flex* electrode 1 before and after the Stim-Stab test. a) the optical image of the unstimulated channel (electrode 1 channel 1) before the Stim-Stab test, b) the optical image of the unstimulated channel after the Stim-Stab test, c) the optical image of the stimulated channel (electrode 1 channel 2) before the Stim-Stab test, d) the optical image of the stimulated channel after the Stim-Stab test, scale bar=0.25 mm.

**Figure S17**. Real-time VT waveform, ElS and CV curves of *Flex* electrode 1. a) Real-time VT waveform of electrode 1 channel 2 stimulated at 12 mA as a function of the pulse number, b) impedance of electrode 1 before and after the Stim-Stab test, and the equivalent circuit model fit, c) phase of electrode 1 before and after the Stim-Stab test, and the equivalent circuit model fit, d) Nyquist plot of electrode 1 before and after the Stim-Stab test, and the equivalent circuit model fit, e) CV curves of electrode 1 before and after the Stim-Stab test.

**Figure S18**. Electrochemical properties of *Flex* electrode 1 before and after the Stim-Stab test. a) 1-Hz impedance before and after the Stim-Stab test, b) $10^3$-Hz impedance before and after the Stim-Stab test, c) 1-Hz phase before and after the Stim-Stab test, d) $10^3$-Hz phase before and after the Stim-Stab test, e) CSC of *Flex* electrode 1 before and after the Stim-Stab test, f) CIC of *Flex* electrode 1 before and after the Stim-Stab test.

**Figure S19.** Optical image of *Flex* electrode 2 before and after the Stim-Stab test. a) the optical image of the unstimulated channel (electrode 2 channel 1) before the Stim-Stab test, b) the optical image of the unstimulated channel after the Stim-Stab test, c) the optical image of the stimulated channel (electrode 2 channel 2) before the Stim-Stab test, d) the optical image of the stimulated channel after the Stim-Stab test, scale bar=0.25 mm.

**Figure S20**. Real-time VT waveform, ElS and CV curves of *Flex* electrode 2. a) real-time VT waveform of electrode 2 channel 2 stimulated at 12 mA as a function of the pulse number, b) impedance of electrode 2 before and after the Stim-Stab test, and the equivalent circuit model fit, c) phase of electrode 2 before and after the Stim-Stab test, and the equivalent circuit model fit, d) Nyquist plot of electrode 2 before and after the Stim-Stab test, and the equivalent circuit model fit, e) CV curves of electrode 2 before and after the Stim-Stab test.

**Figure S21**. Electrochemical properties of *Flex* electrode 2 before and after the Stim-Stab test. a) 1-Hz impedance before and after the Stim-Stab test, b) $10^3$-Hz impedance before and after the Stim-Stab test, c) 1-Hz phase before and after the Stim-Stab test, d) $10^3$-Hz phase before and after the Stim-Stab test, e) CSC of *Flex* electrode 2 before and after the Stim-Stab test, f) CIC of *Flex* electrode 2 before and after the Stim-Stab test.

**Figure S22.** Optical image of *Flex* electrode 3 before and after the Stim-Stab test. a) the optical image of the unstimulated channel (electrode 3 channel 1) before the Stim-Stab test, b) the optical image of the unstimulated channel after the Stim-Stab test, c) the optical image of the stimulated channel (electrode 3 channel 2) before the Stim-Stab test, d) the optical image of the stimulated channel after the Stim-Stab test, scale bar=0.25 mm.

**Figure S23**. Real-time VT waveform, ElS and CV curves of *Flex* electrode 3. a) real-time VT waveform of electrode 3 channel 2 stimulated at 12 mA as a function of the pulse number, b) impedance of electrode 3 before and after the Stim-Stab test, and the equivalent circuit model fit, c) phase of electrode 3 before and after the Stim-Stab test, and the equivalent circuit model fit, d) Nyquist plot of electrode 3 before and after the Stim-Stab test, and the equivalent circuit model fit, e) CV curves of electrode 3 before and after the Stim-Stab test.

**Figure S24**. Electrochemical properties of *Flex* electrode 3 before and after the Stim-Stab test. a) 1-Hz impedance before and after the Stim-Stab test, b) $10^3$-Hz impedance before and after the Stim-Stab test, c) 1-Hz phase before and after the Stim-Stab test, d) $10^3$-Hz phase before and after the Stim-Stab test, e) CSC of *Flex* electrode 3 before and after the Stim-Stab test, f) CIC of *Flex* electrode 3 before and after the Stim-Stab test.

**Figure S25.** Optical image of *Flex* electrode 4 before and after the Stim-Stab test. a) the optical image of the unstimulated channel (electrode 4 channel 1) before the Stim-Stab test, b) the optical image of the unstimulated channel after the Stim-Stab test, c) the optical image of the stimulated channel (electrode 4 channel 2) before the Stim-Stab test, d) the optical image of the stimulated channel after the Stim-Stab test, scale bar=0.25 mm.

**Figure S26**. Real-time VT waveform, ElS and CV curves of *Flex* electrode 4. a) real-time VT waveform of electrode 4 channel 2 stimulated at 16 mA as a function of the pulse number, b) impedance of electrode 4 before and after the Stim-Stab test, and the equivalent circuit model fit, c) phase of electrode 4 before and after the Stim-Stab test, and the equivalent circuit model fit, d) Nyquist plot of electrode 4 before and after the Stim-Stab test, and the equivalent circuit model fit, e) CV curves of electrode 4 before and after the Stim-Stab test.

**Figure S27**. Electrochemical properties of *Flex* electrode 4 before and after the Stim-Stab test. a) 1-Hz impedance before and after the Stim-Stab test, b) $10^3$-Hz impedance before and after the Stim-Stab test, c) 1-Hz phase before and after the Stim-Stab test, d) $10^3$-Hz phase before and after the Stim-Stab test, e) CSC of *Flex* electrode 4 before and after the Stim-Stab test, f) CIC of *Flex* electrode 4 before and after the Stim-Stab test.

**Figure S28.** Optical image of *Flex* electrode 5 before and after the Stim-Stab test. a) the optical image of the unstimulated channel (electrode 5 channel 1) before the Stim-Stab test, b) the optical image of the unstimulated channel after the Stim-Stab test, c) the optical image of the stimulated channel (electrode 5 channel 2) before the Stim-Stab test, d) the optical image of the stimulated channel after the Stim-Stab test, scale bar=0.25 mm.

**Figure S29**. Real-time VT waveform, ElS and CV curves of *Flex* electrode 5. a) Real-time VT waveform of electrode 5 channel 2 stimulated at 16 mA as a function of the pulse number, b) impedance of electrode 5 before and after the Stim-Stab test, and the equivalent circuit model fit, c) phase of electrode 5 before and after the Stim-Stab test, and the equivalent circuit model fit, d) Nyquist plot of electrode 5 before and after the Stim-Stab test, and the equivalent circuit model fit, e) CV curves of electrode 5 before and after the Stim-Stab test.

**Figure S30**. Electrochemical properties of *Flex* electrode 5 before and after the stim-stab test. a) 1-Hz impedance before and after the Stim-Stab test, b) $10^3$-Hz impedance before and after the Stim-Stab test, c) 1-Hz phase before and after the Stim-Stab test, d) $10^3$-Hz phase before and after the Stim-Stab test, e) CSC of *Flex* electrode 5 before and after the Stim-Stab test, f) CIC of *Flex* electrode 5 before and after the Stim-Stab test.

**Figure S31.** Optical image of *Flex* electrode 6 before and after the Stim-Stab test. a) the optical image of the unstimulated channel (electrode 6 channel 1) before the Stim-Stab test, b) the optical image of the unstimulated channel after the Stim-Stab test, c) the optical image of the stimulated channel (electrode 6 channel 2) before the Stim-Stab test, d) the optical image of the stimulated channel after the Stim-Stab test, scale bar=0.25 mm.

**Figure S32**. Real-time VT waveform, ElS and CV curves of *Flex* electrode 6. a) real-time VT waveform of electrode 6 channel 2 stimulated at 16 mA as a function of the pulse number, b) impedance of electrode 6 before and after the Stim-Stab test, and the equivalent circuit model fit, c) phase of electrode 6 before and after the Stim-Stab test, and the equivalent circuit model fit, d) Nyquist plot of electrode 6 before and after the Stim-Stab test, and the equivalent circuit model fit, e) CV curves of electrode 6 before and after the Stim-Stab test.

**Figure S33**. Electrochemical properties of *Flex* electrode 6 before and after the Stim-Stab test. a) 1-Hz impedance before and after the Stim-Stab test, b) $10^3$-Hz impedance before and after the Stim-Stab test, c) 1-Hz phase before and after the Stim-Stab test, d) $10^3$-Hz phase before and after the Stim-Stab test, e) CSC of *Flex* electrode 6 before and after the Stim-Stab test, f) CIC of *Flex* electrode 6 before and after the Stim-Stab test.

**Table S1.** Parameters of the ECM for unstimulated and stimulated electrode channels before and after the Stim-Stab test.

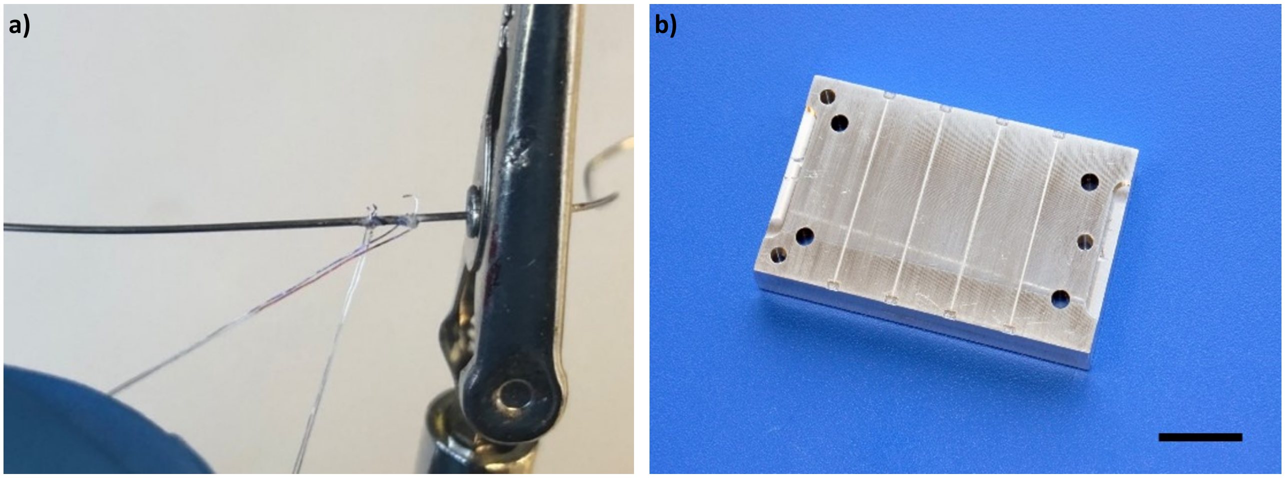


**Figure S1**. a) optical image of the setup to make the helical structure, b) optical image of the mould for the silicone encapsulation surrounding the helical Pt leads, the scale bar=2.0 cm.

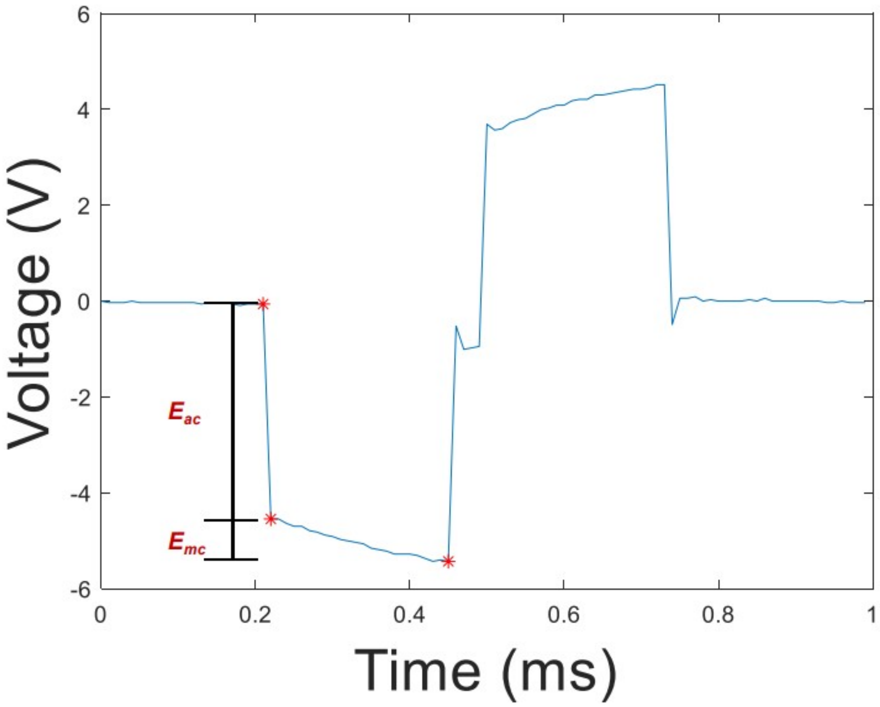


**Figure S2**. Representative VT waveform of a *Flex* electrode in response to the current pulsing.

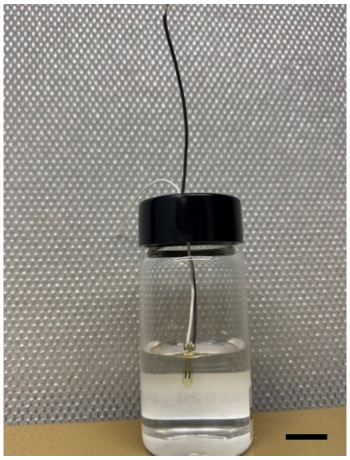

**Figure S3**. Optical image of the integrated *Flex* electrode partially soaked in PBS for the Stim-Stab test (scale bar=10 mm).

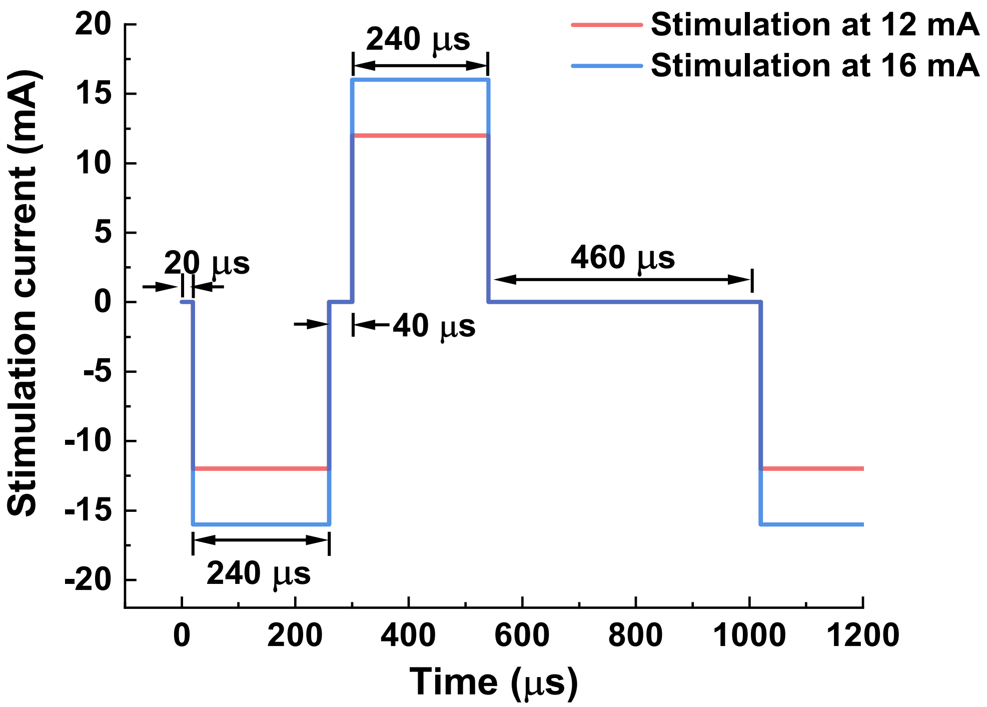


**Figure S4.** Current waveforms generated by the stimulator for the Stim-Stab test.

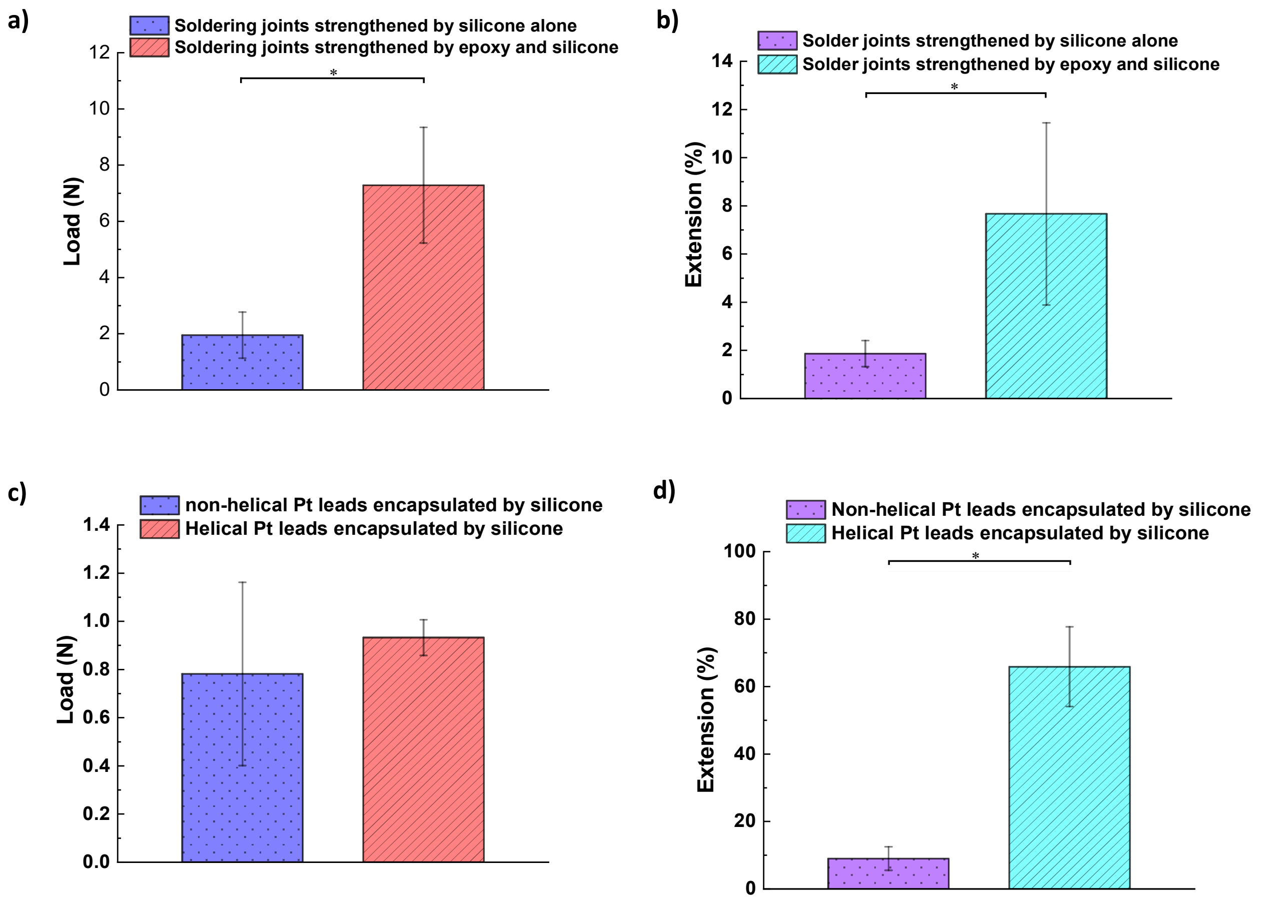


**Figure S5.** Mechanical properties of the integrated electrodes for acute and chronic vagus nerve stimulation. a) the maximum load at the breakage for integrated acute configuration electrodes with and without epoxy reinforcement, b) the maximum extension at the breakage for integrated acute configuration electrodes with and without epoxy reinforcement (acute vagus nerve stimulation), c) the maximum load at the breakage for integrated chronic configuration electrodes with nonhelical and helical Pt leads, d) the maximum extension at the breakage for integrated chronic configuration electrodes with nonhelical and helical Pt leads.

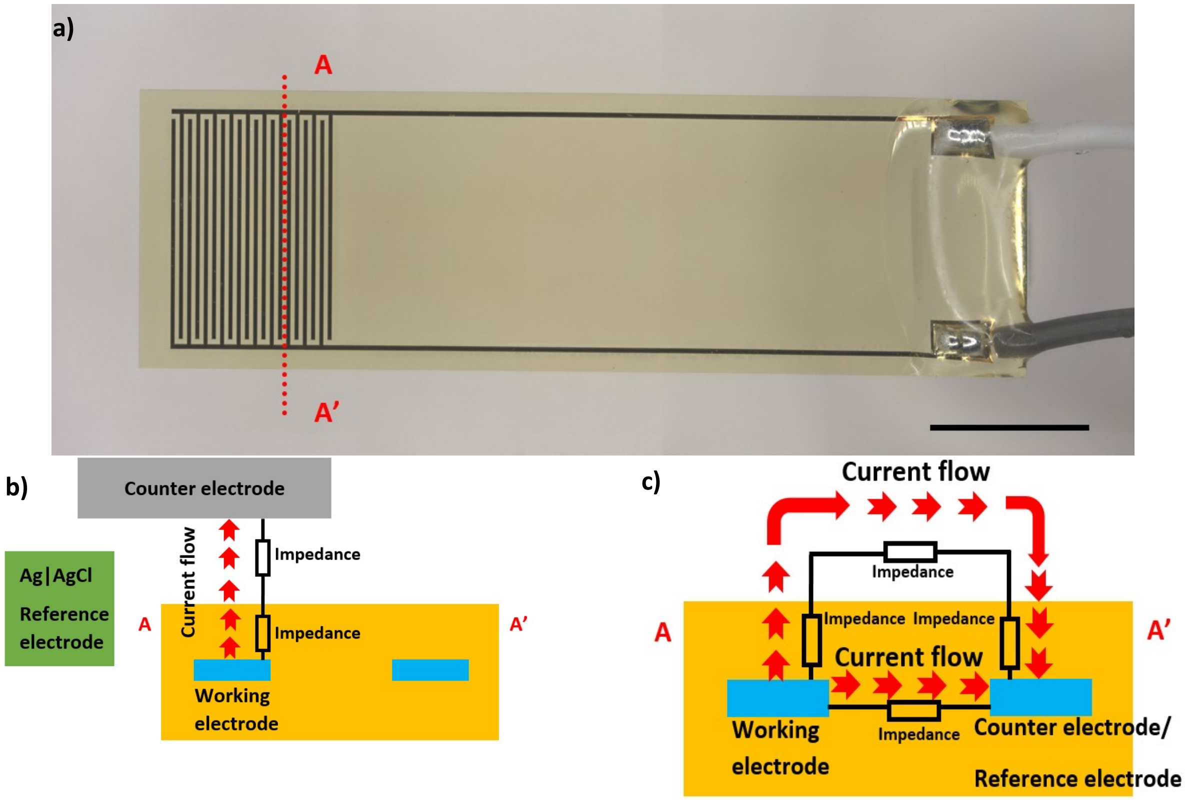


**Figure S6**. a) Optical image of an integrated IDE for the accelerated aging test, scale bar=5.0mm, b) the current flow for 3-electrode configuration, c) the current flow for 2-electrode configuration.

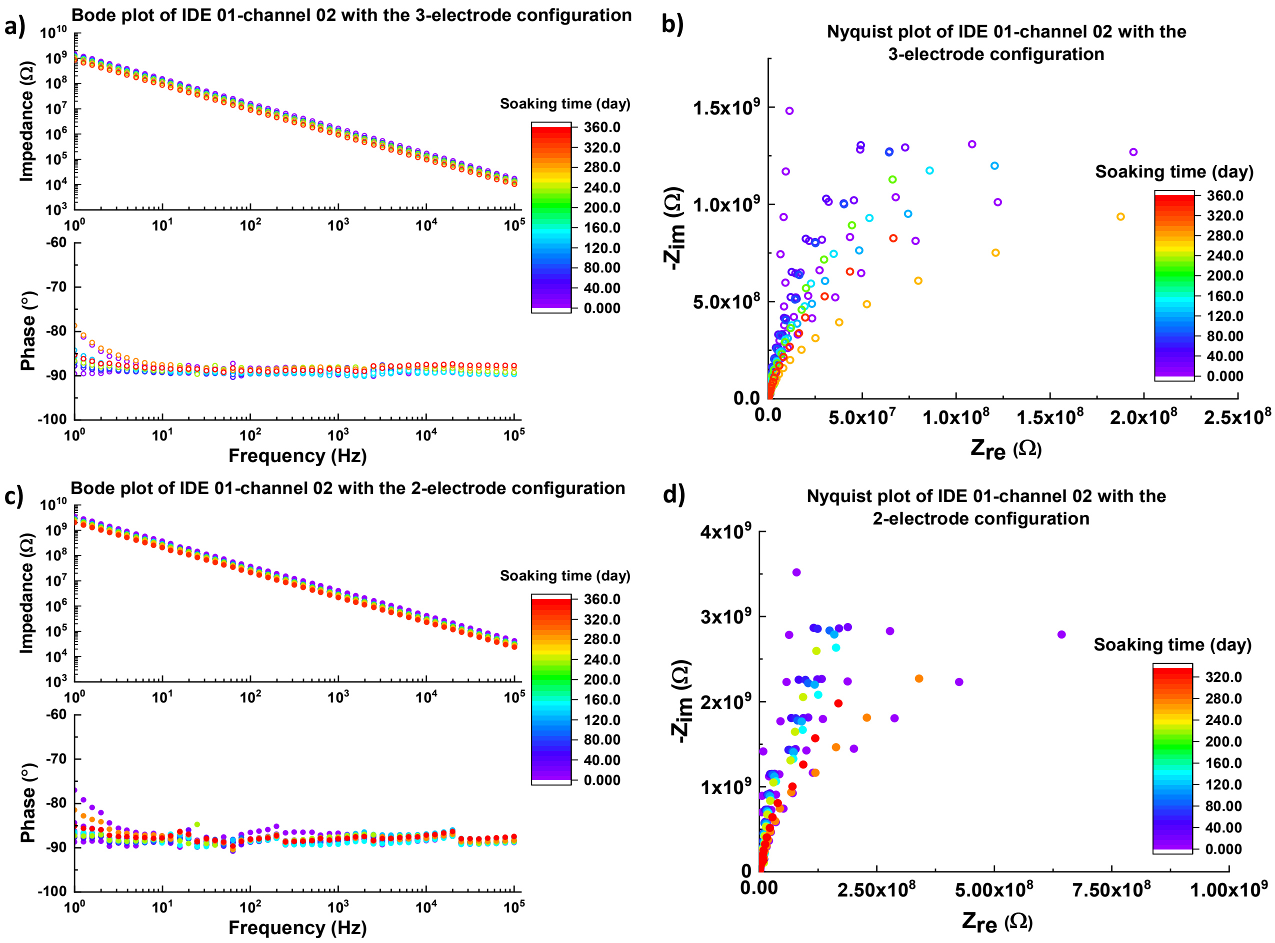


**Figure S7.** Impedance spectra of the IDE 01-channel 02 device as a function of the soaking time. a) Bode plot of IDE 01-channel 02 device with 3-electrode configuration as a function of the soaking time, b) Nyquist plot of IDE 01-channel 02 device with 3-electrode configuration as a function of the soaking time, c) Bode plot of IDE 01-channel 02 device with 2-electrode configuration as a function of the soaking time, d) Nyquist plot of IDE 01-channel 02 device with 2-electrode configuration as a function of the soaking time.

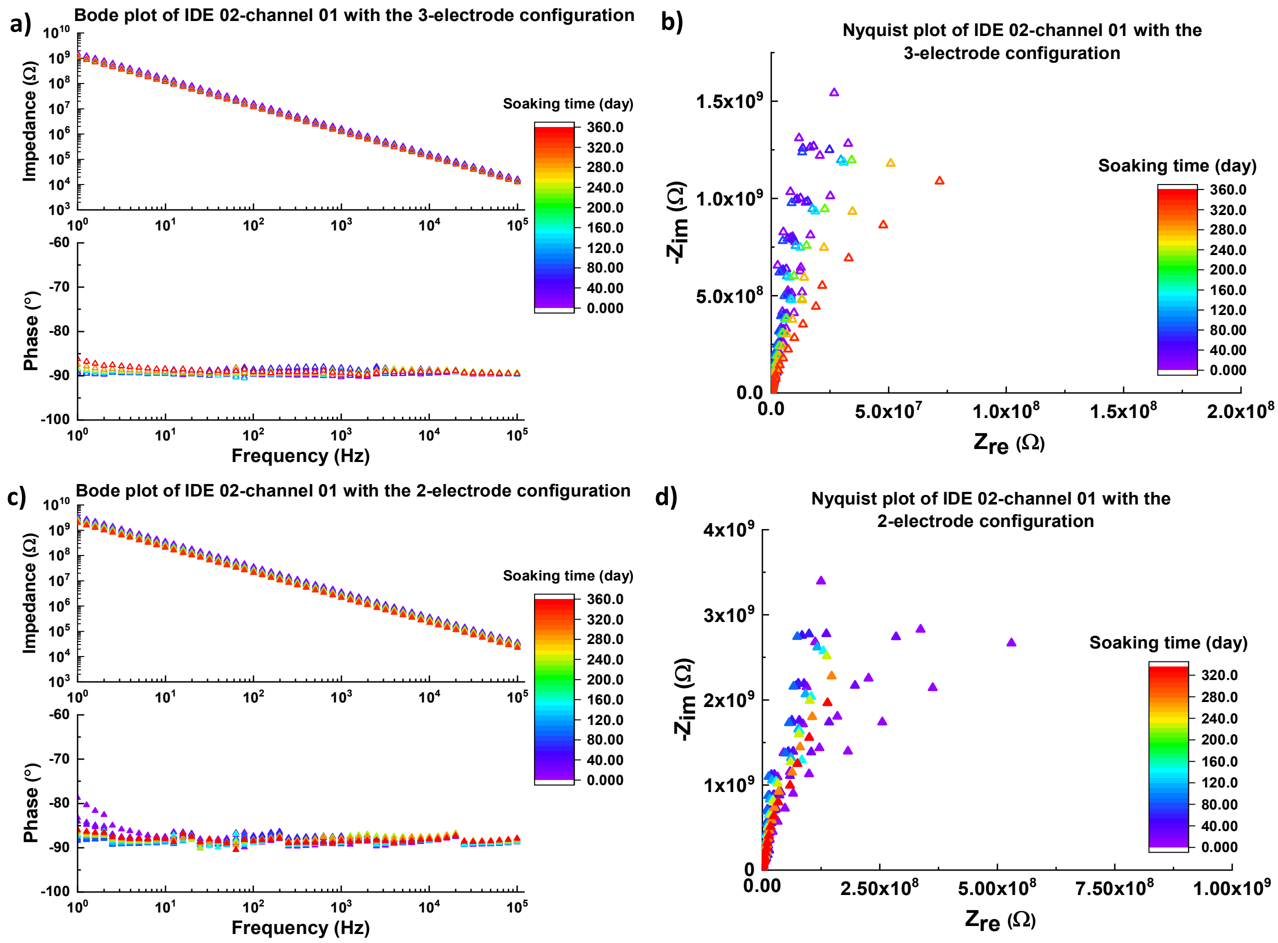


**Figure S8.** Impedance spectra of the representative IDE device (IDE 02-channel 01) as a function of the soaking time. a) Bode plot of IDE 02-channel 01 device with the 3-electrode configuration as a function of the soaking time, b) Nyquist plot of IDE 02-channel 01 device with the 3-electrode configuration as a function of the soaking time, c) Bode plot of IDE 02-channel 01 device with the 2-electrode configuration as a function of the soaking time, d) Nyquist plot of IDE 02-channel 01 device with the 2-electrode configuration as a function of the soaking time.

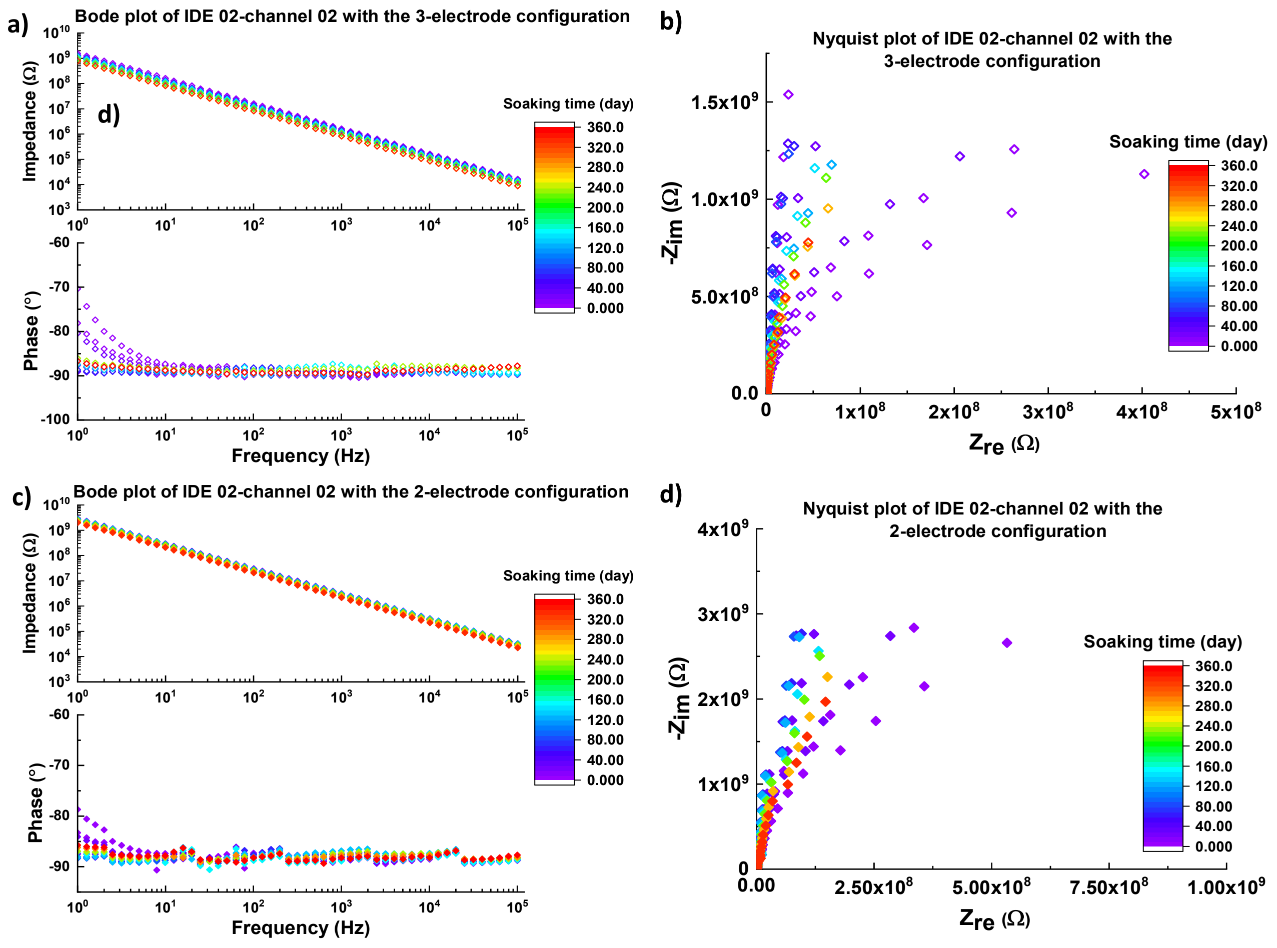


**Figure S9**. Impedance spectra of the IDE 02-channel 02 device as a function of the soaking time. a) Bode plot of IDE 02-channel 02 device with 3-electrode configuration as a function of the soaking time, b) Nyquist plot of IDE 02-channel 02 device with 3-electrode configuration as a function of the soaking time, c) Bode plot of IDE 02-channel 02 device with 2-electrode configuration as a function of the soaking time, d) Nyquist plot of IDE 02-channel 02 device with 2-electrode configuration as a function of the soaking time.

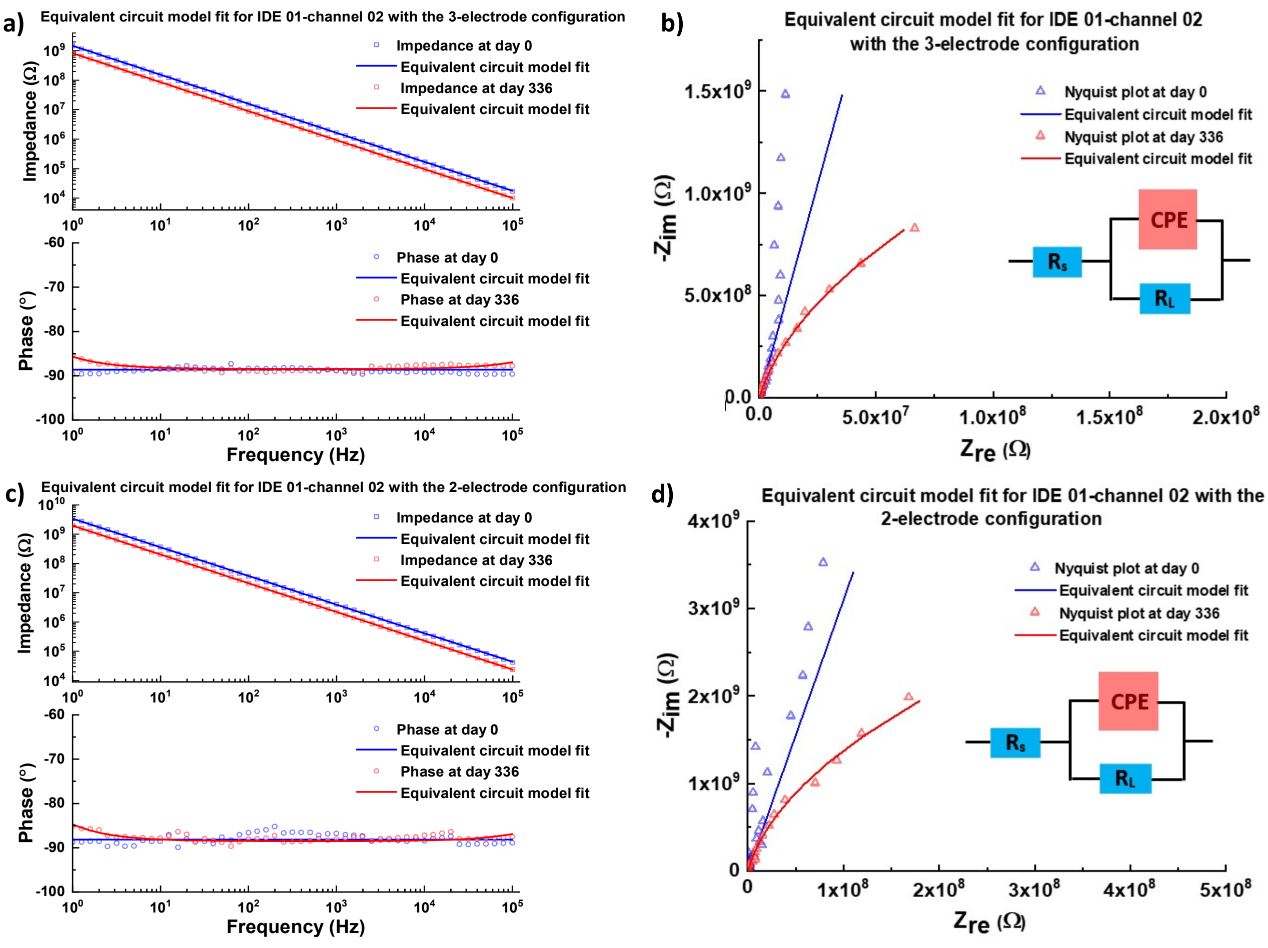


**Figure S10.** Equivalent circuit model fit for the impedance spectra of IDE 01-channel 02. a) an equivalent circuit model fit for the Bode plot of IDE 01-channel 02 with the 3-electrode configuration, b) an equivalent circuit model fit for the Nyquist plot of IDE 01-channel 02 with the 3-electrode configuration, c) an equivalent circuit model fit for the Bode plot of IDE 01-channel 02 with the 2-electrode configuration, d) an equivalent circuit model fit for the Nyquist plot of IDE 01-channel 02 with the 2-electrode configuration.

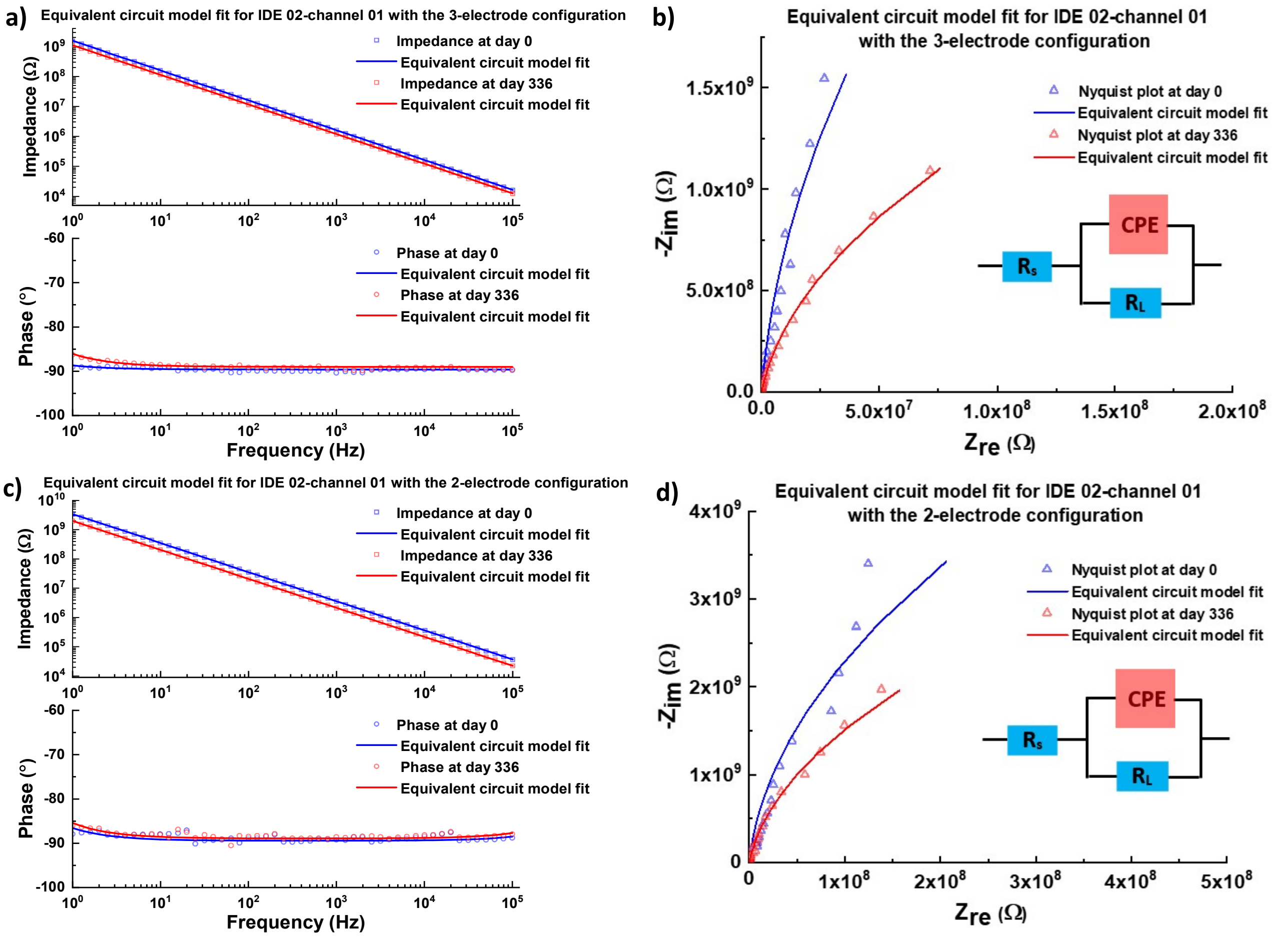


**Figure S11.** Equivalent circuit model fit for the impedance spectra of IDE 02-channel 01. a) an equivalent circuit model fit for the Bode plot of IDE 02-channel 01 with the 3-electrode configuration, b) an equivalent circuit model fit for the Nyquist plot of IDE 02-channel 01 with the 3-electrode configuration, c) an equivalent circuit model fit for the Bode plot of IDE 02-channel 01 with the 2-electrode configuration, d) an equivalent circuit model fit for the Nyquist plot of IDE 02-channel 01 with the 2-electrode configuration.

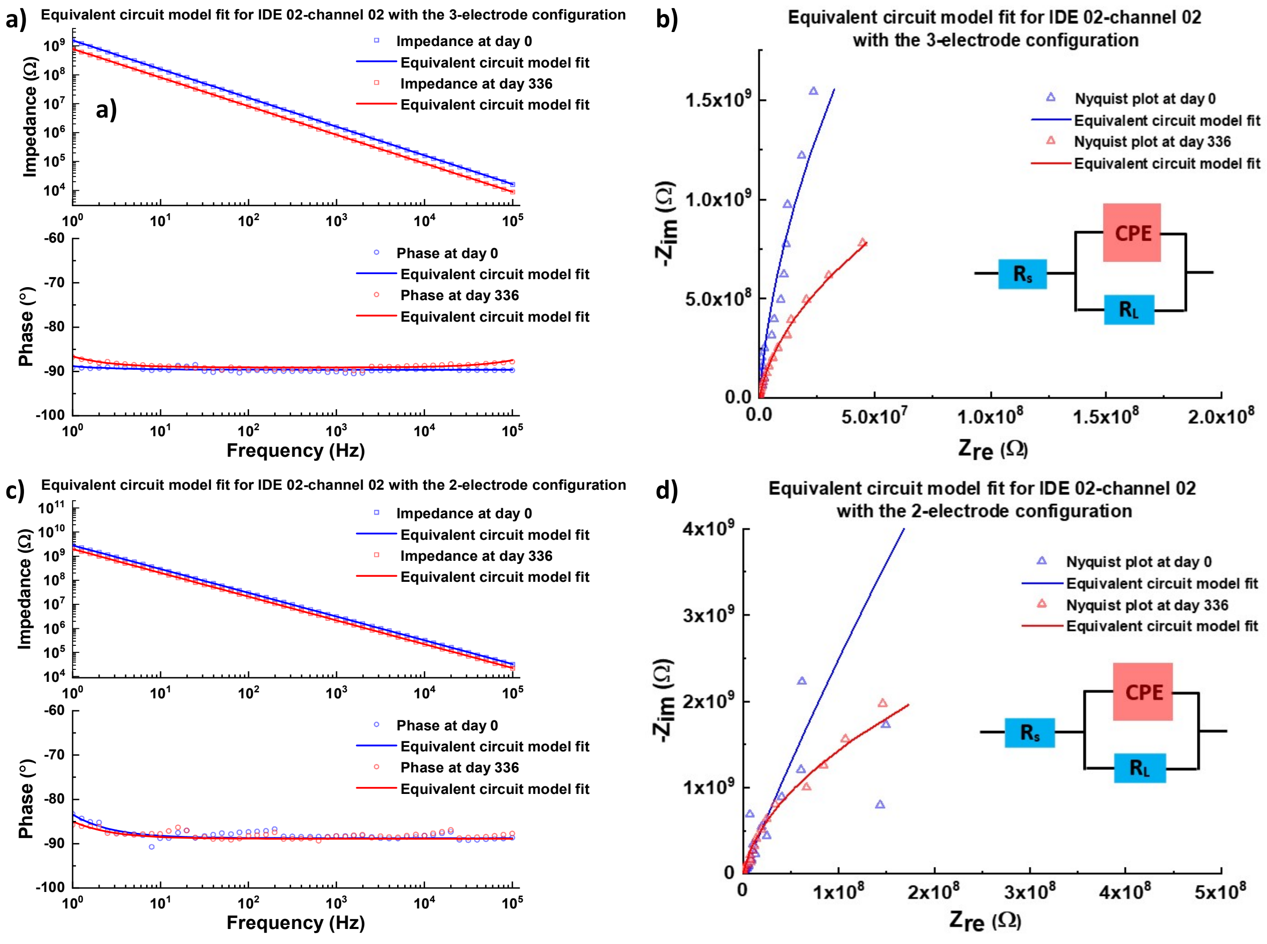


**Figure S12.** Equivalent circuit model fit for the impedance spectra of IDE 02-channel 02. a) an equivalent circuit model fit for the Bode plot of IDE 02-channel 02 with the 3-electrode configuration, b) an equivalent circuit model fit for the Nyquist plot of IDE 02-channel 02 with the 3-electrode configuration, c) an equivalent circuit model fit for the Bode plot of IDE 02-channel 02 with the 2-electrode configuration, d) an equivalent circuit model fit for the Nyquist plot of IDE 02-channel 02 with the 2-electrode configuration.

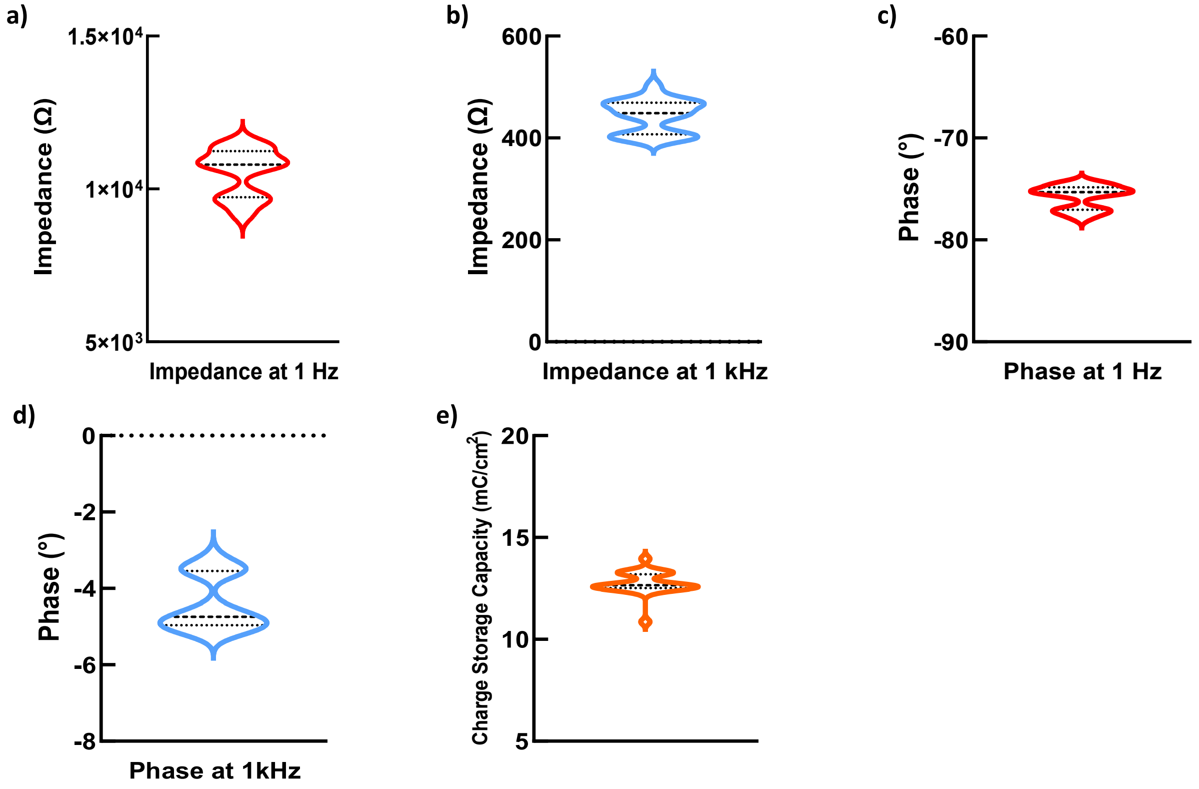


**Figure S13.** Electrochemical properties of *Flex* electrodes (n=24). a) violin curve of the impedance at 1 Hz, b) violin curve of the impedance at $10^3$ Hz, c) violin curve of the phase at 1 Hz, d) violin curve of the phase at $10^3$ Hz, e) violin curve of the CSC.

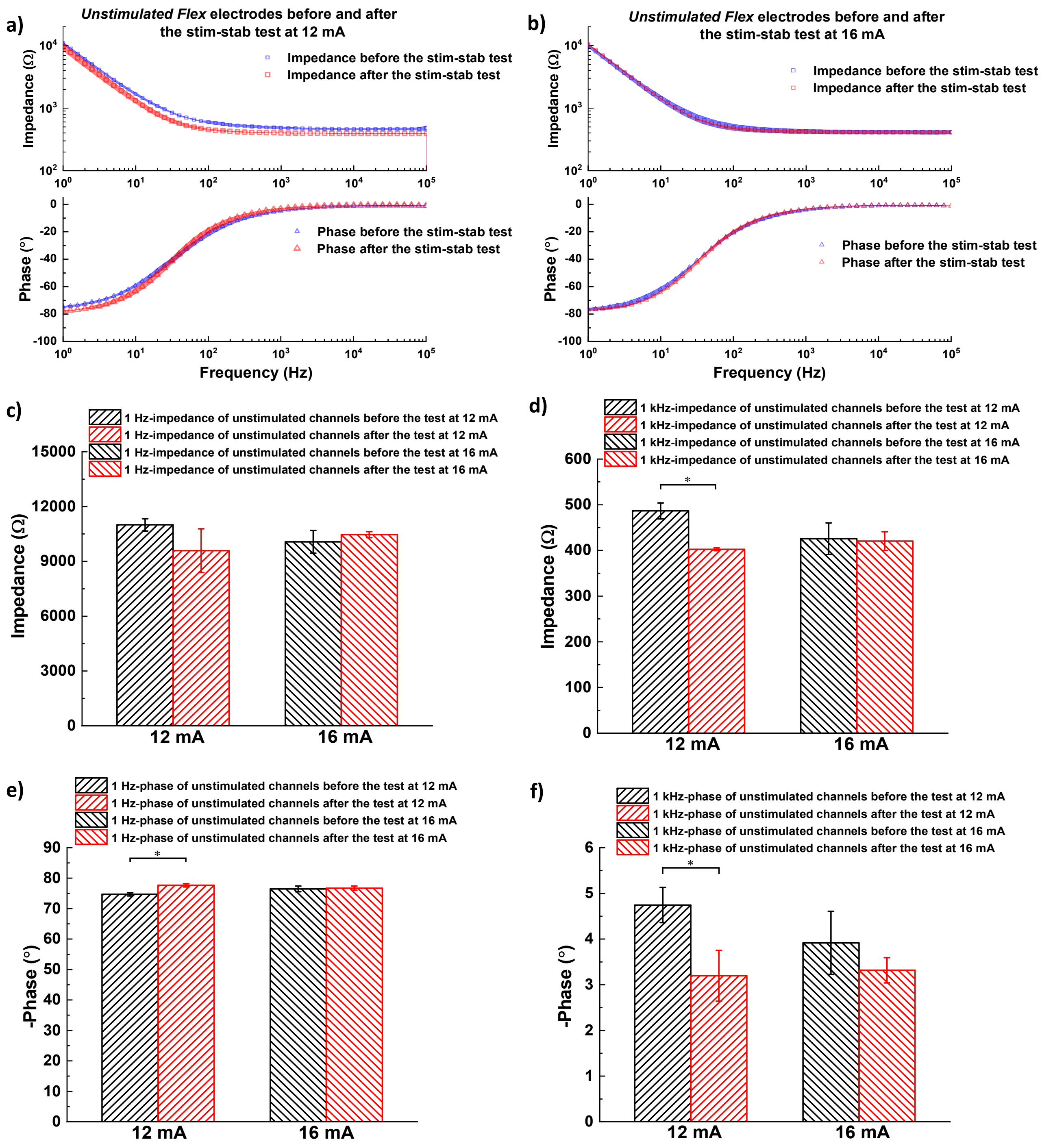
a)
*Unstimulated Flex* electrodes before and after the stim-stab test at 12 mA
Impedance before the stim-stab test
Impedance after the stim-stab test
Impedance (Ω)
Phase before the stim-stab test
Phase after the stim-stab test
Phase (°)
Frequency (Hz)
b)
*Unstimulated Flex* electrodes before and after the stim-stab test at 16 mA
Impedance before the stim-stab test
Impedance after the stim-stab test
Impedance (Ω)
Phase before the stim-stab test
Phase after the stim-stab test
Phase (°)
Frequency (Hz)
c)
1 Hz-impedance of unstimulated channels before the test at 12 mA
1 Hz-impedance of unstimulated channels after the test at 12 mA
1 Hz-impedance of unstimulated channels before the test at 16 mA
1 Hz-impedance of unstimulated channels after the test at 16 mA
Impedance (Ω)
12 mA
16 mA
d)
1 kHz-impedance of unstimulated channels before the test at 12 mA
1 kHz-impedance of unstimulated channels after the test at 12 mA
1 kHz-impedance of unstimulated channels before the test at 16 mA
1 kHz-impedance of unstimulated channels after the test at 16 mA
Impedance (Ω)
12 mA
16 mA
e)
1 Hz-phase of unstimulated channels before the test at 12 mA
1 Hz-phase of unstimulated channels after the test at 12 mA
1 Hz-phase of unstimulated channels before the test at 16 mA
1 Hz-phase of unstimulated channels after the test at 16 mA
-Phase (°)
12 mA
16 mA
f)
1 kHz-phase of unstimulated channels before the test at 12 mA
1 kHz-phase of unstimulated channels after the test at 12 mA
1 kHz-phase of unstimulated channels before the test at 16 mA
1 kHz-phase of unstimulated channels after the test at 16 mA
-Phase (°)
12 mA
16 mA

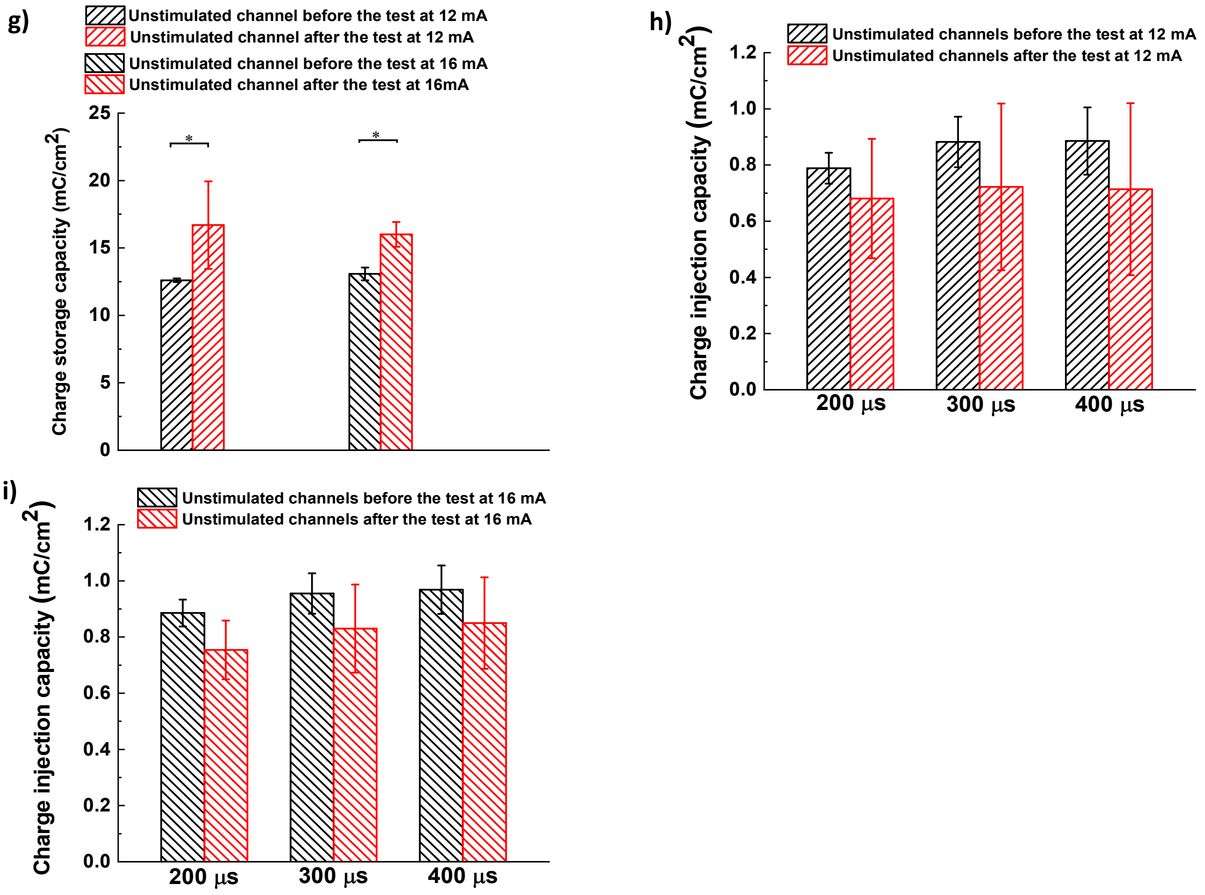


**Figure S14.** Electrical properties of unstimulated electrode channels before and after the Stim-Stab test. a) Bode plots of unstimulated electrode channels before and after the Stim-Stab test at 12 mA (shaded areas indicate the standard deviation), b) Bode plots of unstimulated electrode channels before and after the Stim-Stab test at 16 mA (shaded areas indicate the standard deviation), c) 1-Hz impedance of unstimulated electrode channels before and after the Stim-Stab test, d) $10^3$-Hz impedance of unstimulated electrode channels before and after the Stim-Stab test, e) 1-Hz phase of unstimulated electrode channels before and after the Stim-Stab test, f) $10^3$-Hz phase of unstimulated electrode channels before and after the Stim-Stab test, g) CSC of unstimulated electrode channels before and after the Stim-Stab test, h) CIC of unstimulated electrode channels before and after the Stim-Stab test at 12 mA, i) CIC of unstimulated electrode channels before and after the Stim-Stab test at 16 mA.

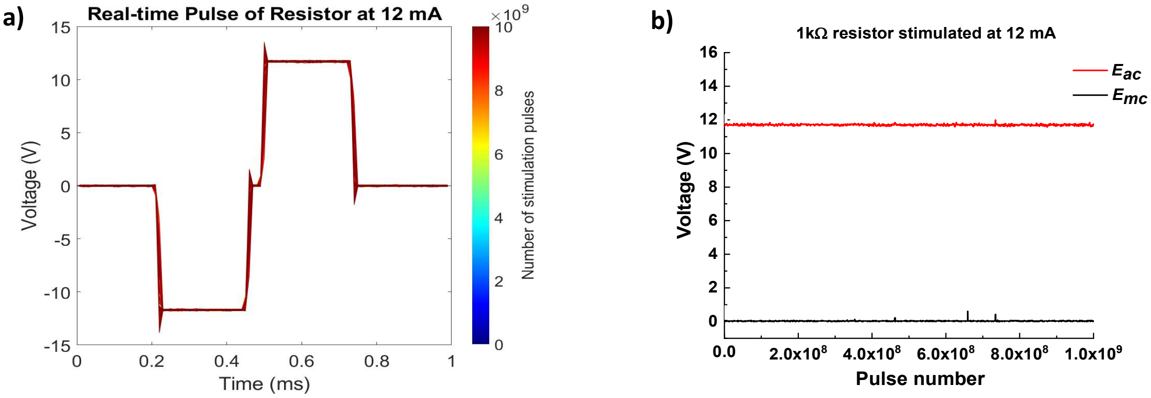


**Figure S15.** a) Real-time stimulation pulse of the 1-kΩ resistor as a function of the pulse number, b) Eac and Emc curves of the 1-kΩ resistor as a function of the pulse number.

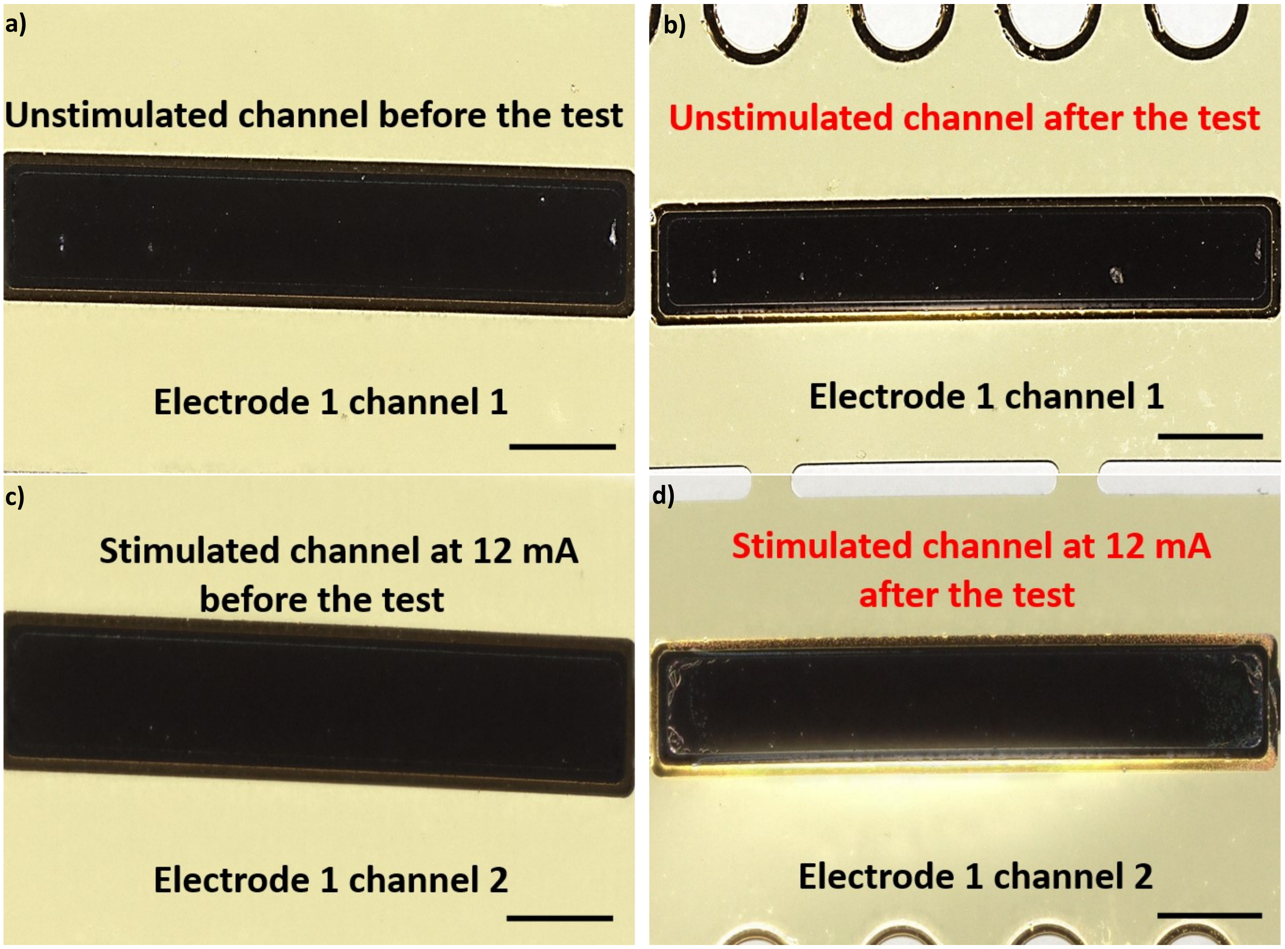


**Figure S16.** Optical image of *Flex* electrode 1 before and after the Stim-Stab test. a) the optical image of the unstimulated channel (electrode 1 channel 1) before the Stim-Stab test, b) the optical image of the unstimulated channel after the Stim-Stab test, c) the optical image of the stimulated channel (electrode 1 channel 2) before the Stim-Stab test, d) the optical image of the stimulated channel after the Stim-Stab test, scale bar=0.25 mm.

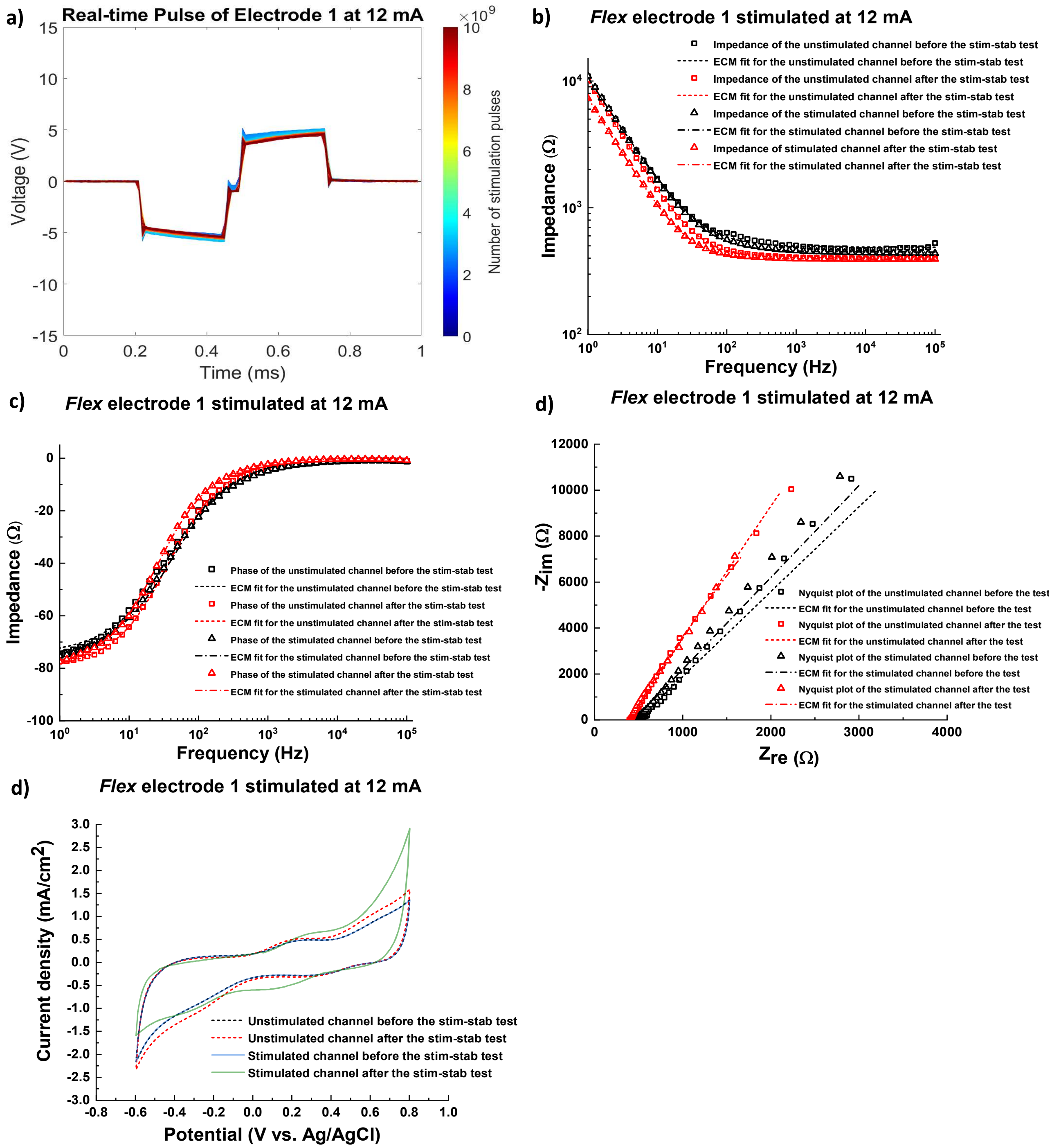


**Figure S17.** Real-time VT waveform, EIS and CV curves of *Flex* electrode 1. a) Real-time VT waveform of electrode 1 channel 2 stimulated at 12 mA as a function of the pulse number, b) impedance of electrode 1 before and after the Stim-Stab test, and the equivalent circuit model fit, c) phase of electrode 1 before and after the Stim-Stab test, and the equivalent circuit model fit, d) Nyquist plot of electrode 1 before and after the Stim-Stab test, and the equivalent circuit model fit, e) CV curves of electrode 1 before and after the Stim-Stab test.

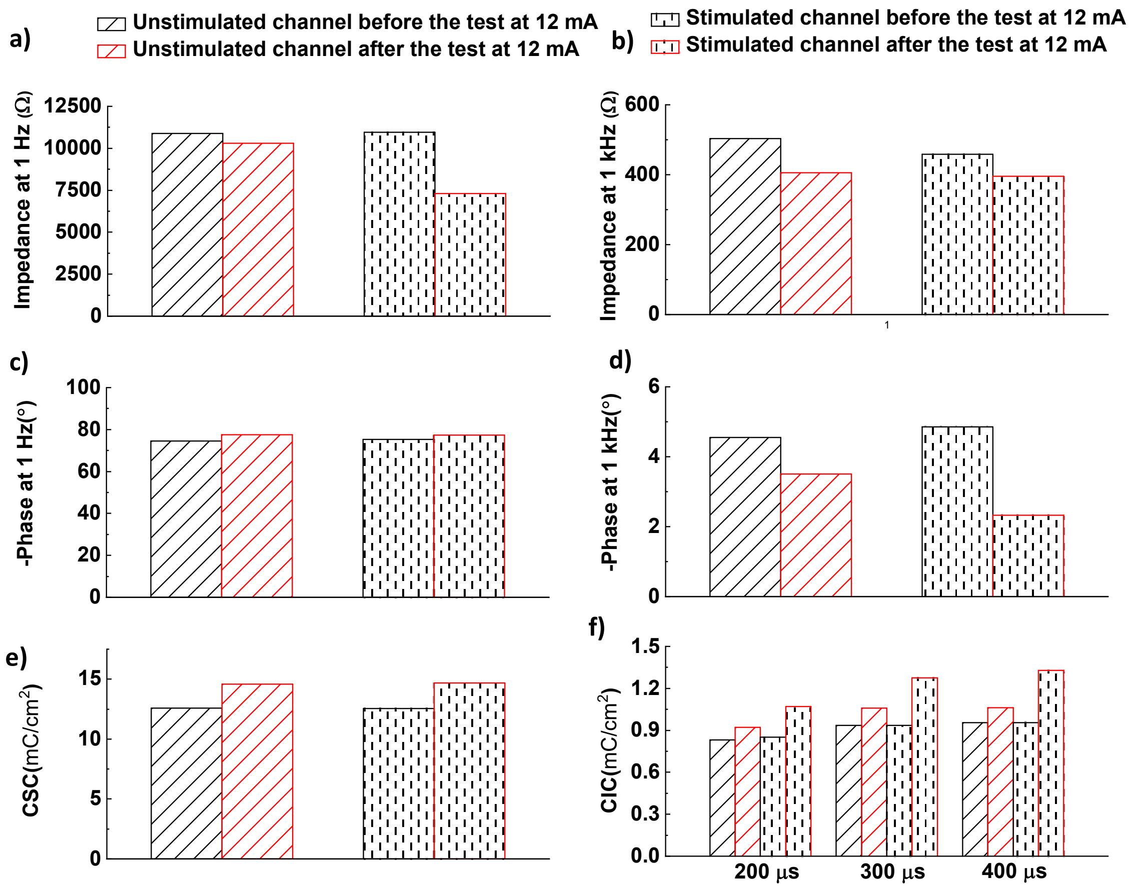


**Figure S18.** Electrochemical properties of *Flex* electrode 1 before and after the Stim-Stab test. a) 1-Hz impedance before and after the Stim-Stab test, b) $10^3$-Hz impedance before and after the Stim-Stab test, c) 1-Hz phase before and after the Stim-Stab test, d) $10^3$-Hz phase before and after the Stim-Stab test, e) CSC of *Flex* electrode 1 before and after the Stim-Stab test, f) CIC of *Flex* electrode 1 before and after the Stim-Stab test.

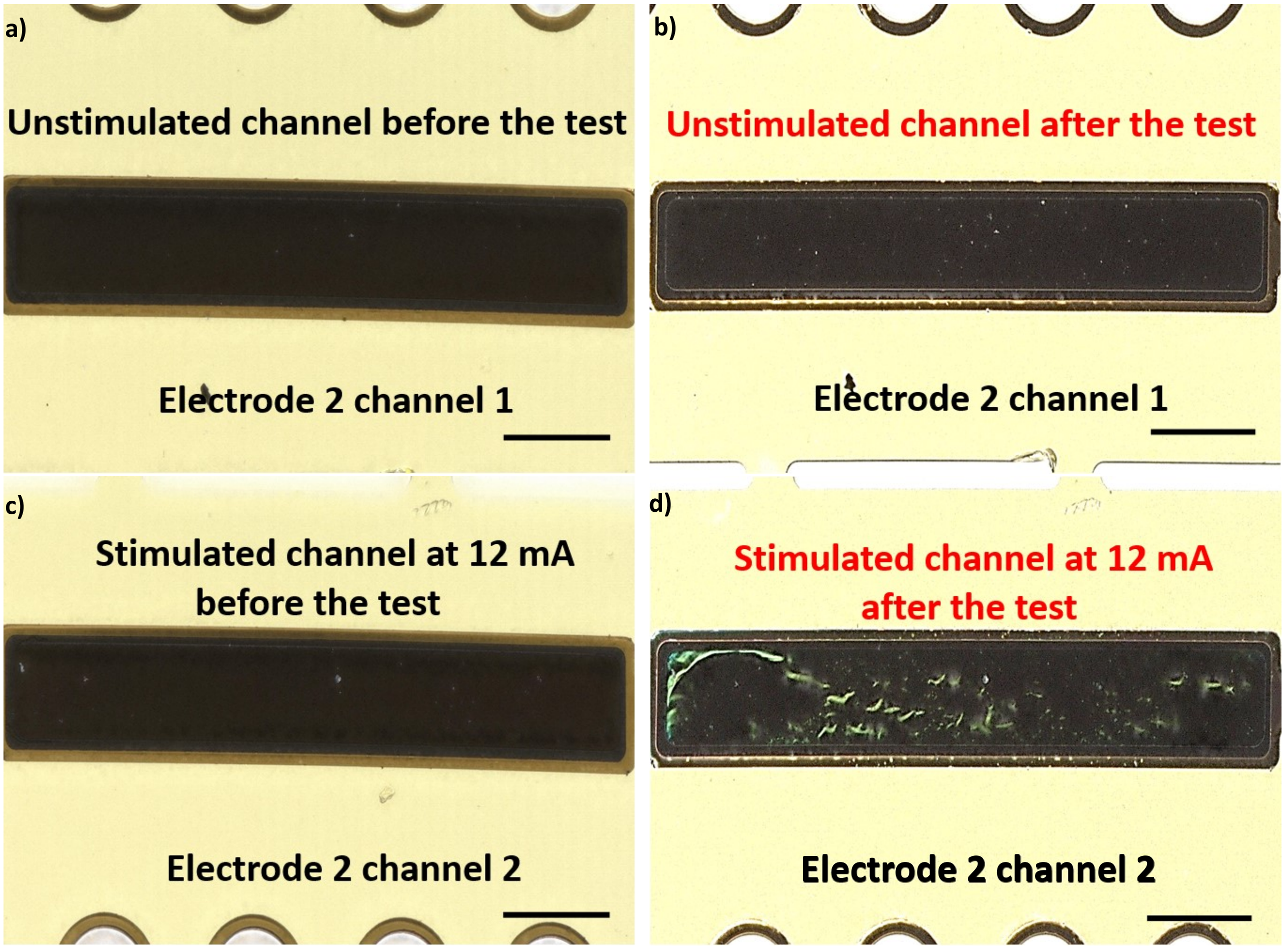


**Figure S19.** Optical image of *Flex* electrode 2 before and after the Stim-Stab test. a) the optical image of the unstimulated channel (electrode 2 channel 1) before the Stim-Stab test, b) the optical image of the unstimulated channel after the Stim-Stab test, c) the optical image of the stimulated channel (electrode 2 channel 2) before the Stim-Stab test, d) the optical image of the stimulated channel after the Stim-Stab test, scale bar=0.25 mm.

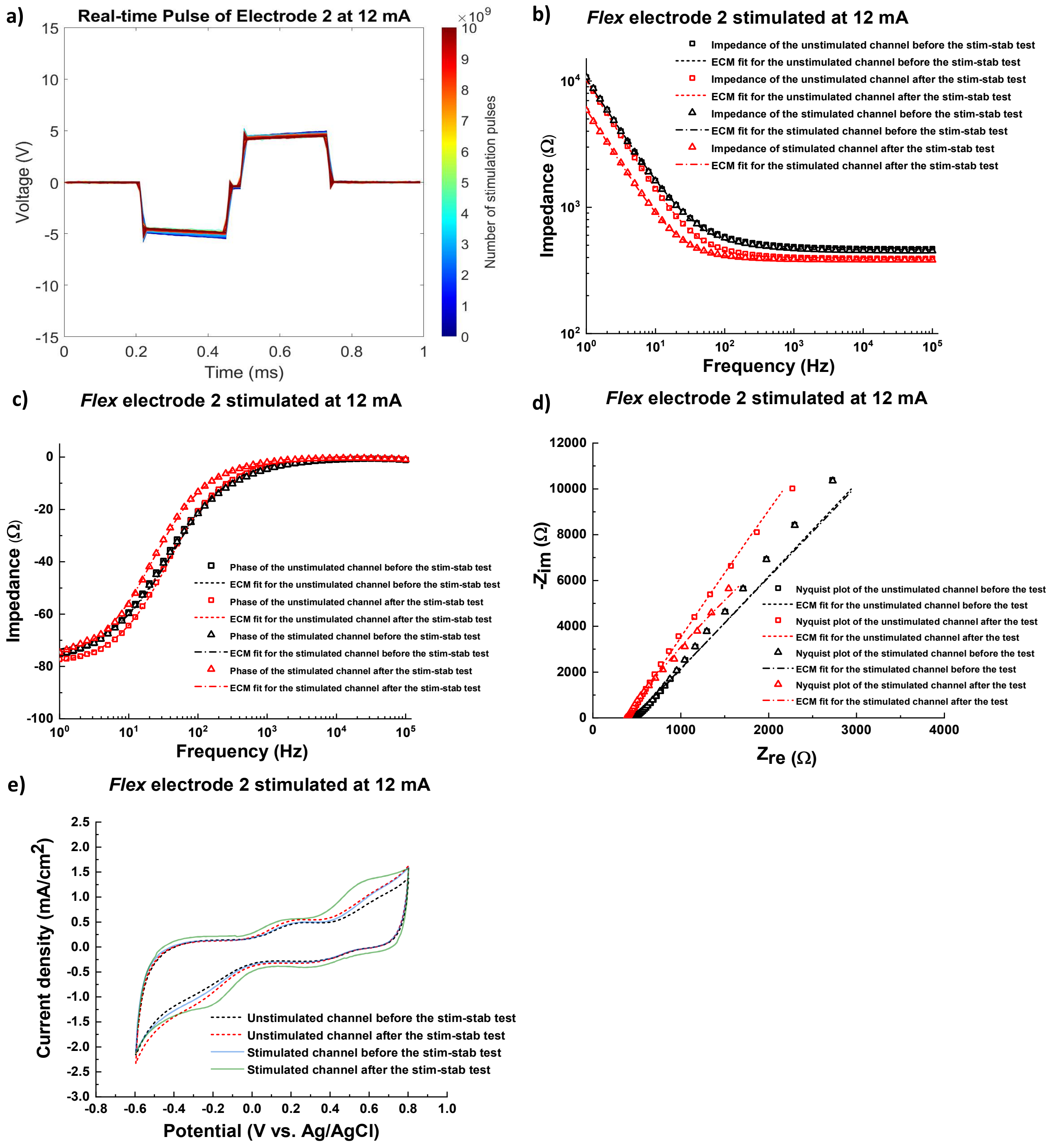


**Figure S20**. Real-time VT waveform, EIS and CV curves of *Flex* electrode 2. a) real-time VT waveform of electrode 2 channel 2 stimulated at 12 mA as a function of the pulse number, b) impedance of electrode 2 before and after the Stim-Stab test, and the equivalent circuit model fit, c) phase of electrode 2 before and after the Stim-Stab test, and the equivalent circuit model fit, d) Nyquist plot of electrode 2 before and after the Stim-Stab test, and the equivalent circuit model fit, e) CV curves of electrode 2 before and after the Stim-Stab test.

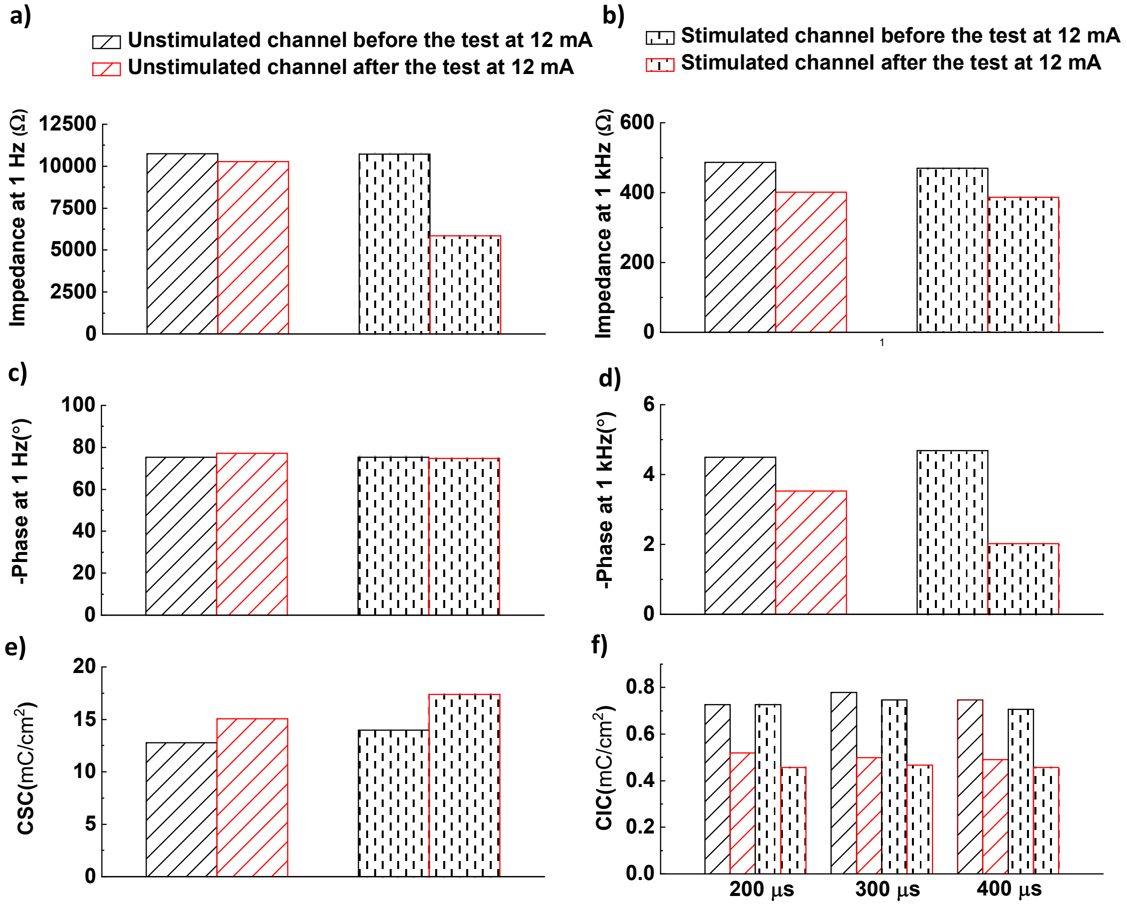


**Figure S21**. Electrochemical properties of *Flex* electrode 2 before and after the Stim-Stab test. a) 1-Hz impedance before and after the Stim-Stab test, b) $10^3$-Hz impedance before and after the Stim-Stab test, c) 1-Hz phase before and after the Stim-Stab test, d) $10^3$-Hz phase before and after the Stim-Stab test, e) CSC of *Flex* electrode 2 before and after the Stim-Stab test, f) CIC of *Flex* electrode 2 before and after the Stim-Stab test.

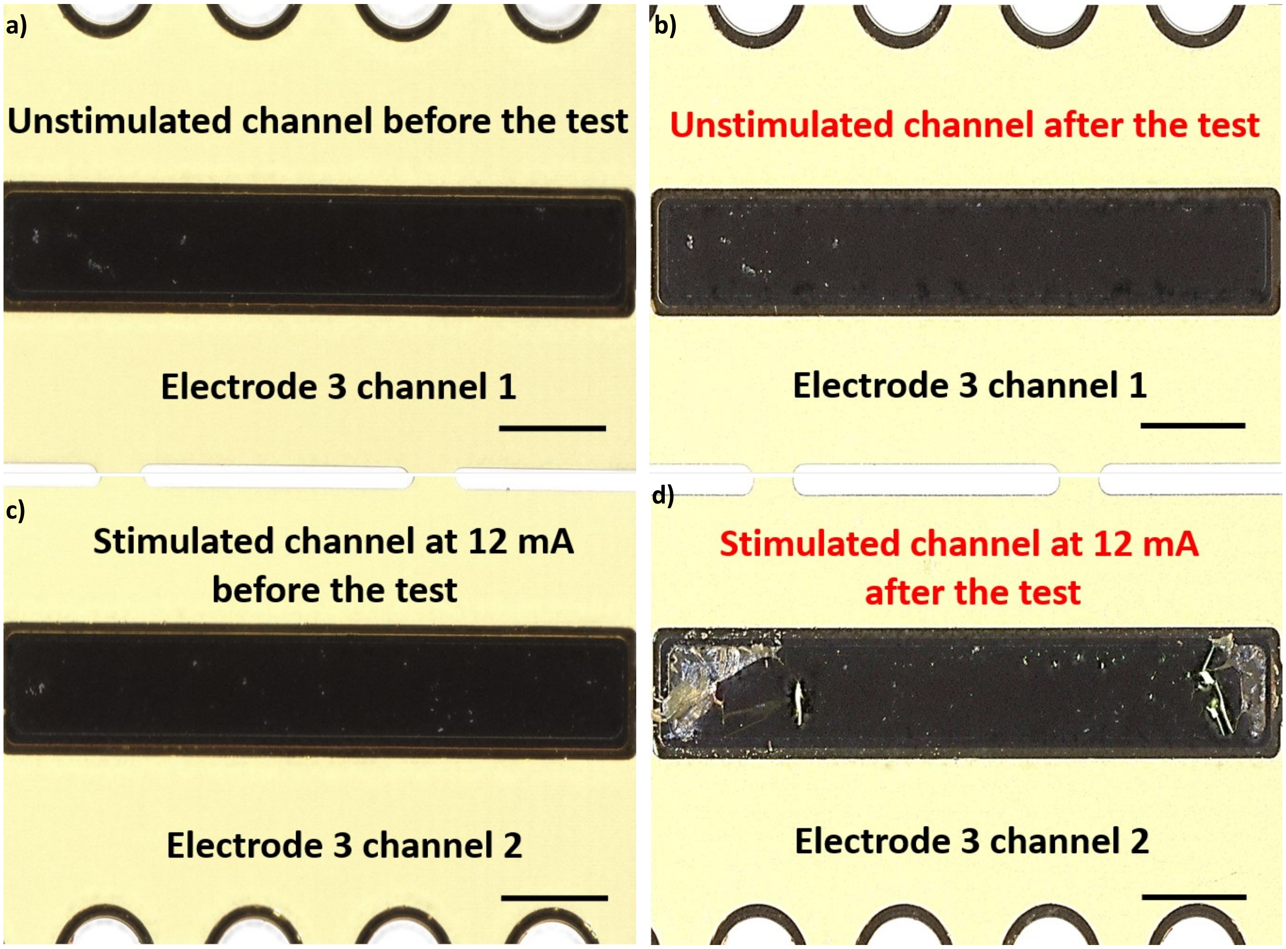


**Figure S22.** Optical image of *Flex* electrode 3 before and after the Stim-Stab test. a) the optical image of the unstimulated channel (electrode 3 channel 1) before the Stim-Stab test, b) the optical image of the unstimulated channel after the Stim-Stab test, c) the optical image of the stimulated channel (electrode 3 channel 2) before the Stim-Stab test, d) the optical image of the stimulated channel after the Stim-Stab test, scale bar=0.25 mm.

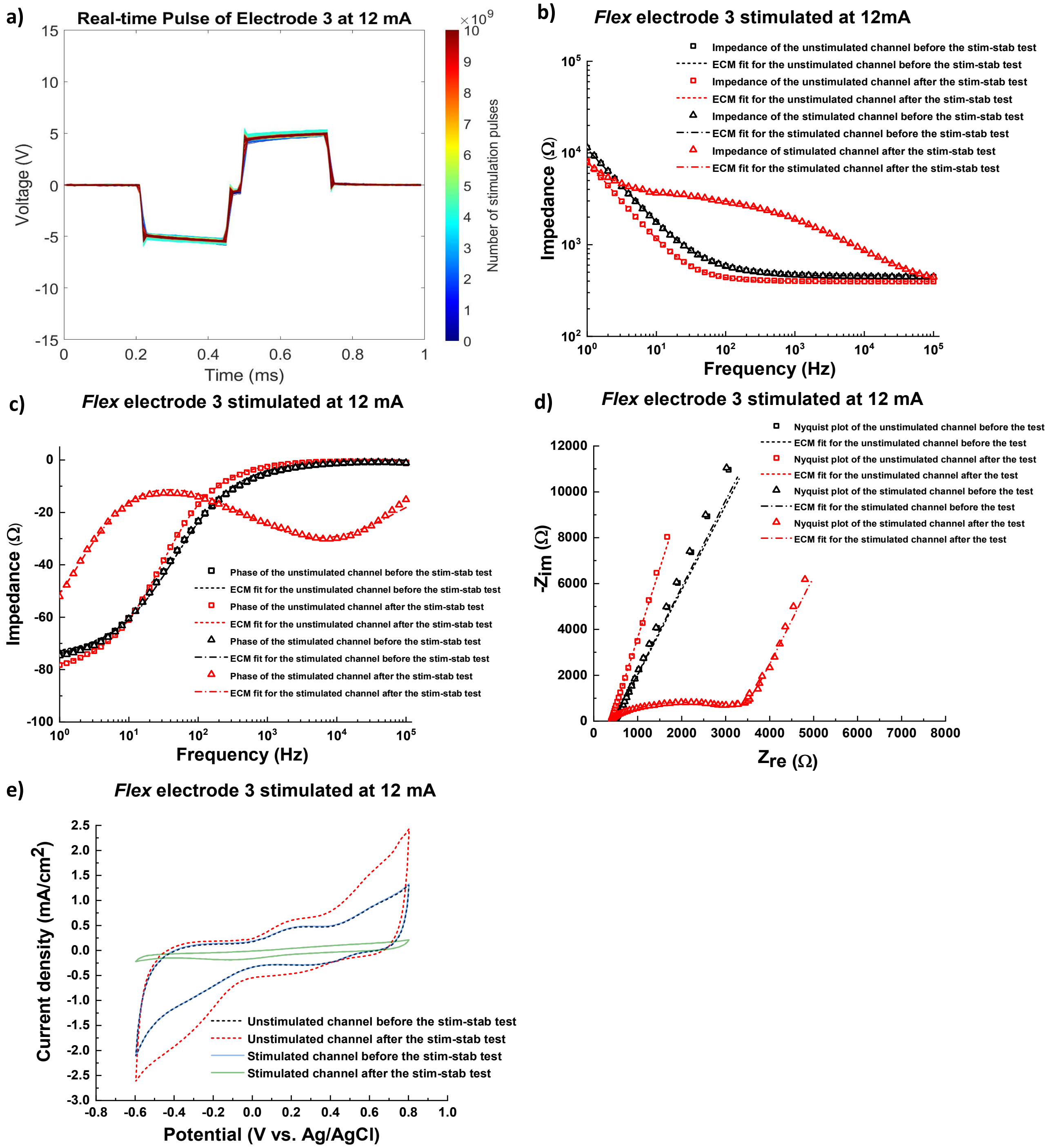


**Figure S23**. Real-time VT waveform, EIS and CV curves of *Flex* electrode 3. a) real-time VT waveform of electrode 3 channel 2 stimulated at 12 mA as a function of the pulse number, b) impedance of electrode 3 before and after the Stim-Stab test, and the equivalent circuit model fit, c) phase of electrode 3 before and after the Stim-Stab test, and the equivalent circuit model fit, d) Nyquist plot of electrode 3 before and after the Stim-Stab test, and the equivalent circuit model fit, e) CV curves of electrode 3 before and after the Stim-Stab test.

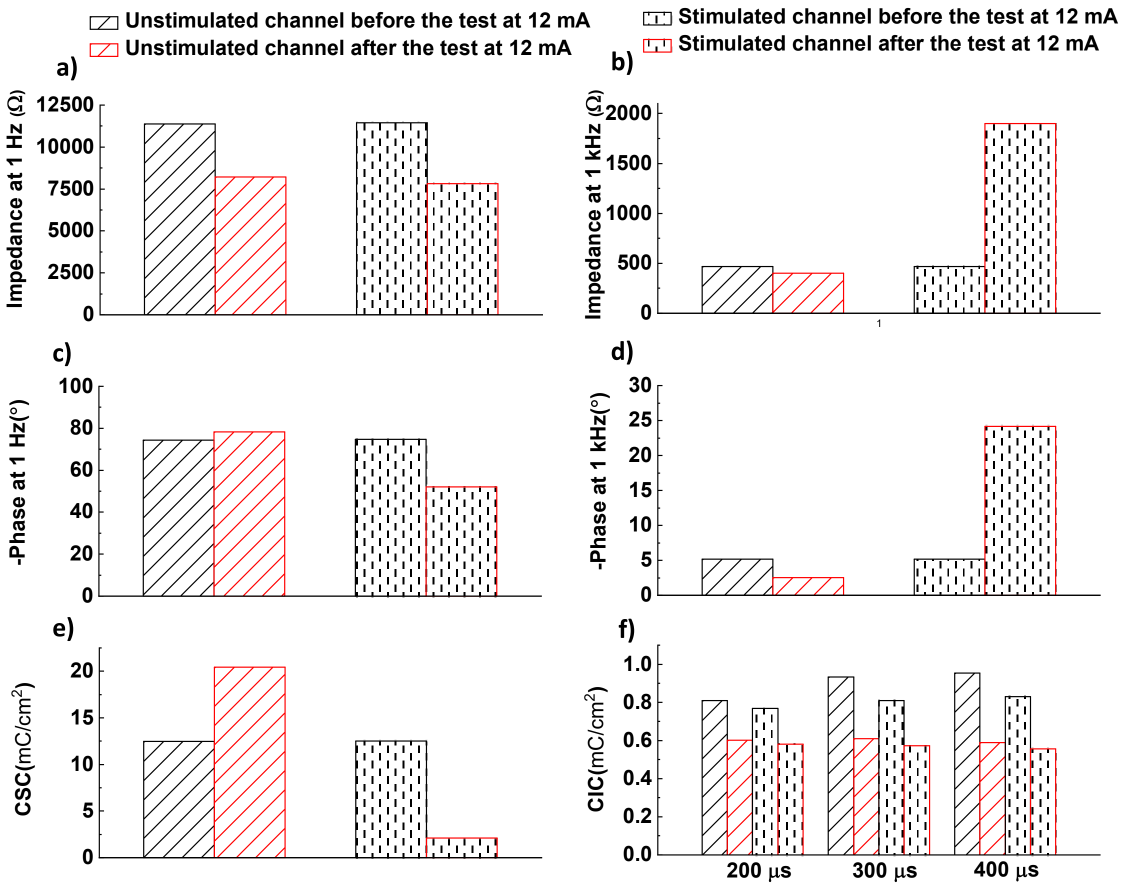


**Figure S24**. Electrochemical properties of *Flex* electrode 3 before and after the Stim-Stab test. a) 1-Hz impedance before and after the Stim-Stab test, b) $10^3$-Hz impedance before and after the Stim-Stab test, c) 1-Hz phase before and after the Stim-Stab test, d) $10^3$-Hz phase before and after the Stim-Stab test, e) CSC of *Flex* electrode 3 before and after the Stim-Stab test, f) CIC of *Flex* electrode 3 before and after the Stim-Stab test.

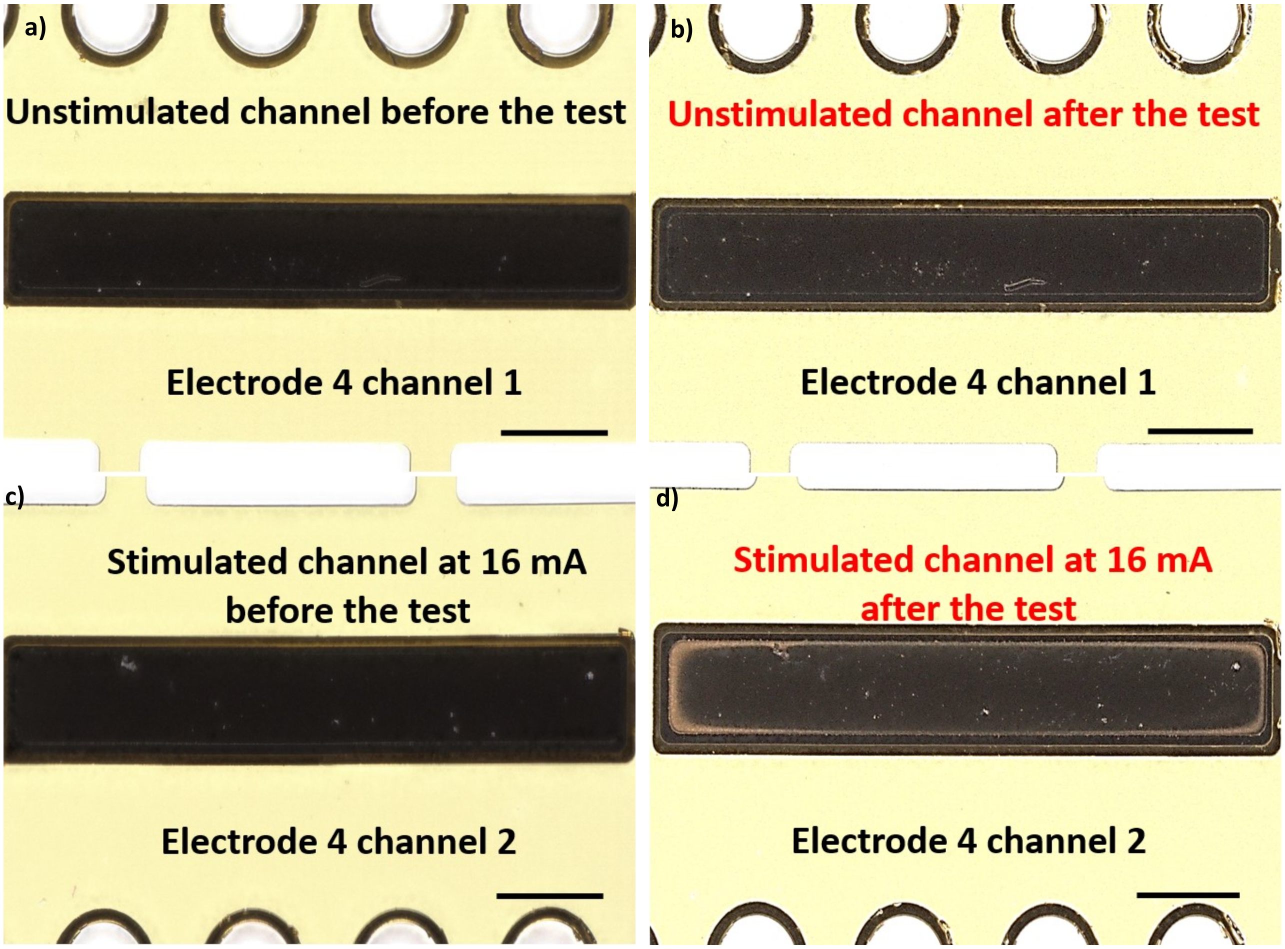


**Figure S25.** Optical image of *Flex* electrode 4 before and after the Stim-Stab test. a) the optical image of the unstimulated channel (electrode 4 channel 1) before the Stim-Stab test, b) the optical image of the unstimulated channel after the Stim-Stab test, c) the optical image of the stimulated channel (electrode 4 channel 2) before the Stim-Stab test, d) the optical image of the stimulated channel after the Stim-Stab test, scale bar=0.25 mm.

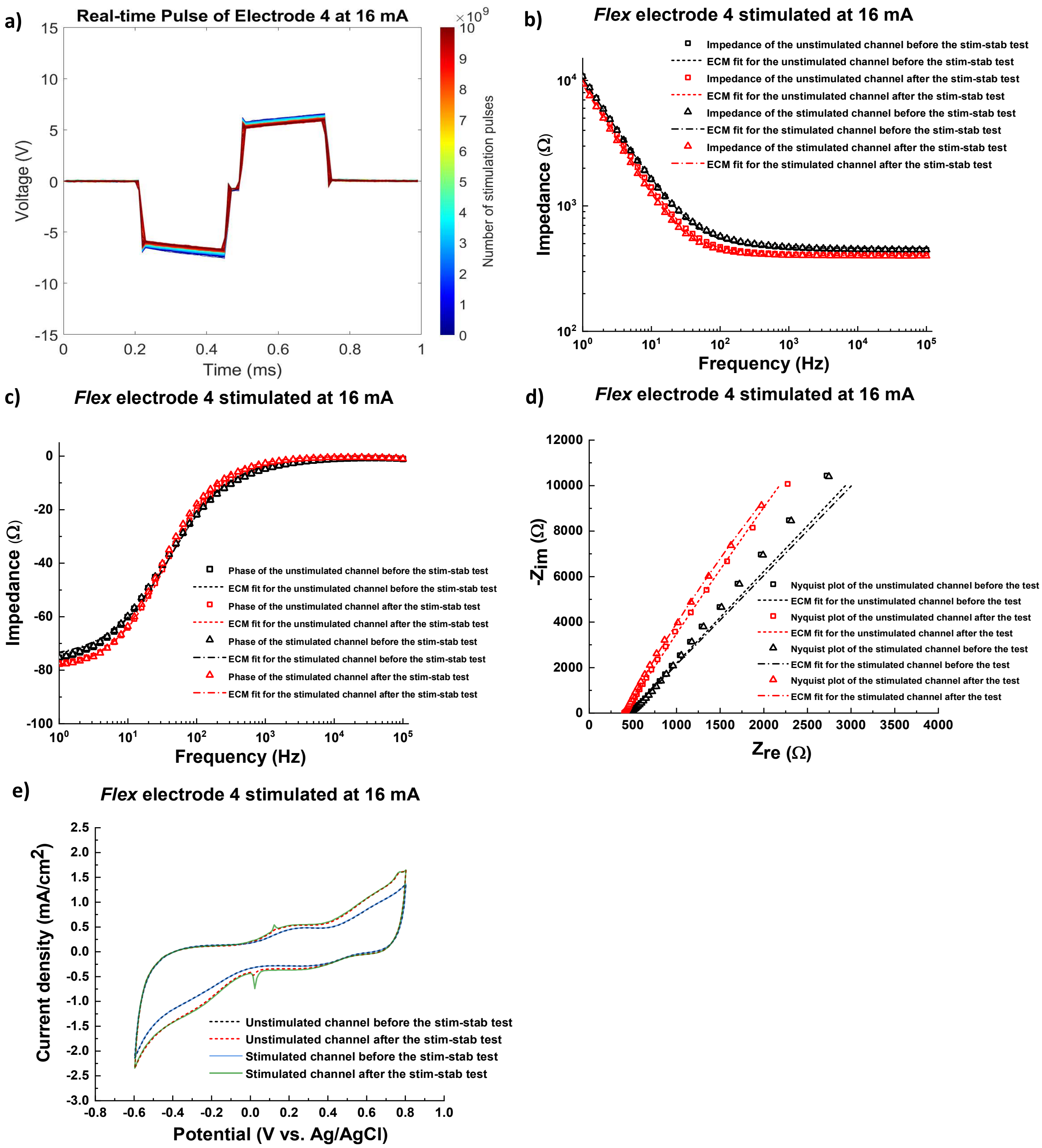


**Figure S26**. Real-time VT waveform, EIS and CV curves of *Flex* electrode 4. a) real-time VT waveform of electrode 4 channel 2 stimulated at 16 mA as a function of the pulse number, b) impedance of electrode 4 before and after the Stim-Stab test, and the equivalent circuit model fit, c) phase of electrode 4 before and after the Stim-Stab test, and the equivalent circuit model fit, d) Nyquist plot of electrode 4 before and after the Stim-Stab test, and the equivalent circuit model fit, e) CV curves of electrode 4 before and after the Stim-Stab test.

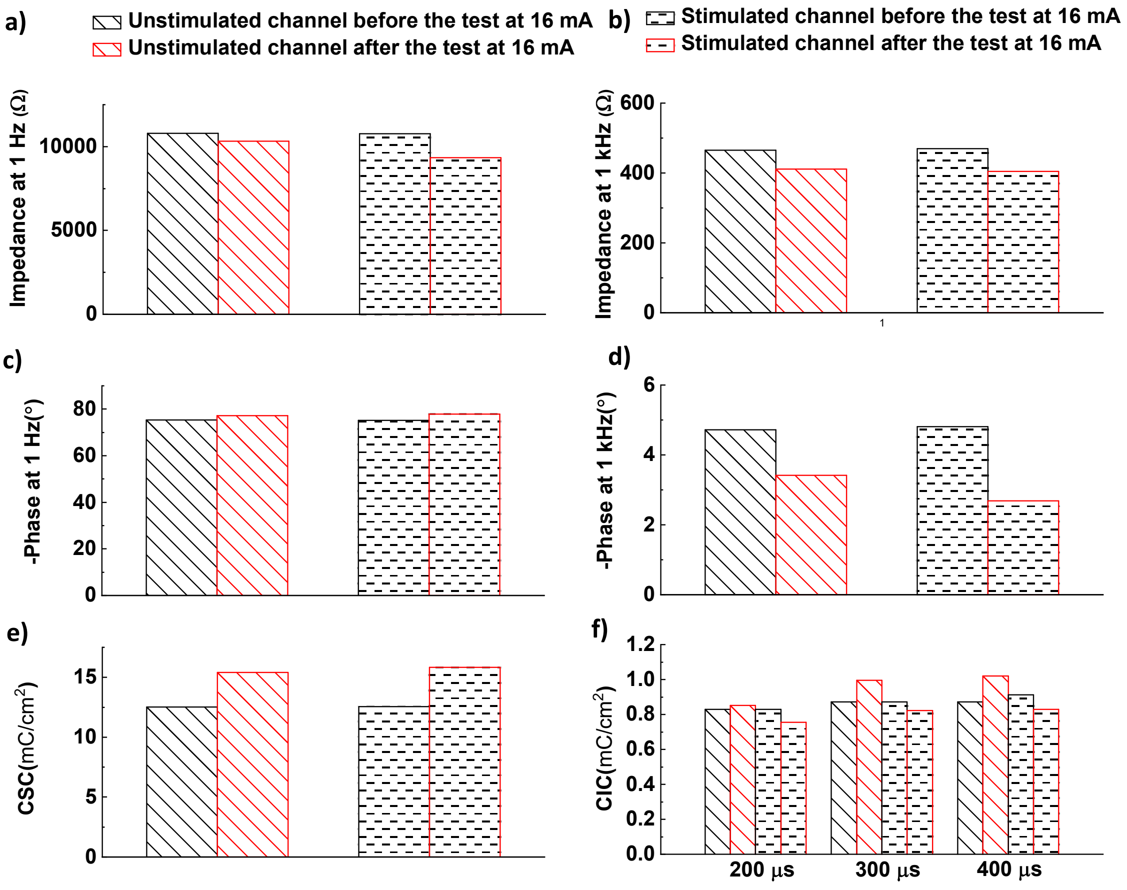


**Figure S27**. Electrochemical properties of *Flex* electrode 4 before and after the Stim-Stab test. a) 1-Hz impedance before and after the Stim-Stab test, b) $10^3$-Hz impedance before and after the Stim-Stab test, c) 1-Hz phase before and after the Stim-Stab test, d) $10^3$-Hz phase before and after the Stim-Stab test, e) CSC of *Flex* electrode 4 before and after the Stim-Stab test, f) CIC of *Flex* electrode 4 before and after the Stim-Stab test.

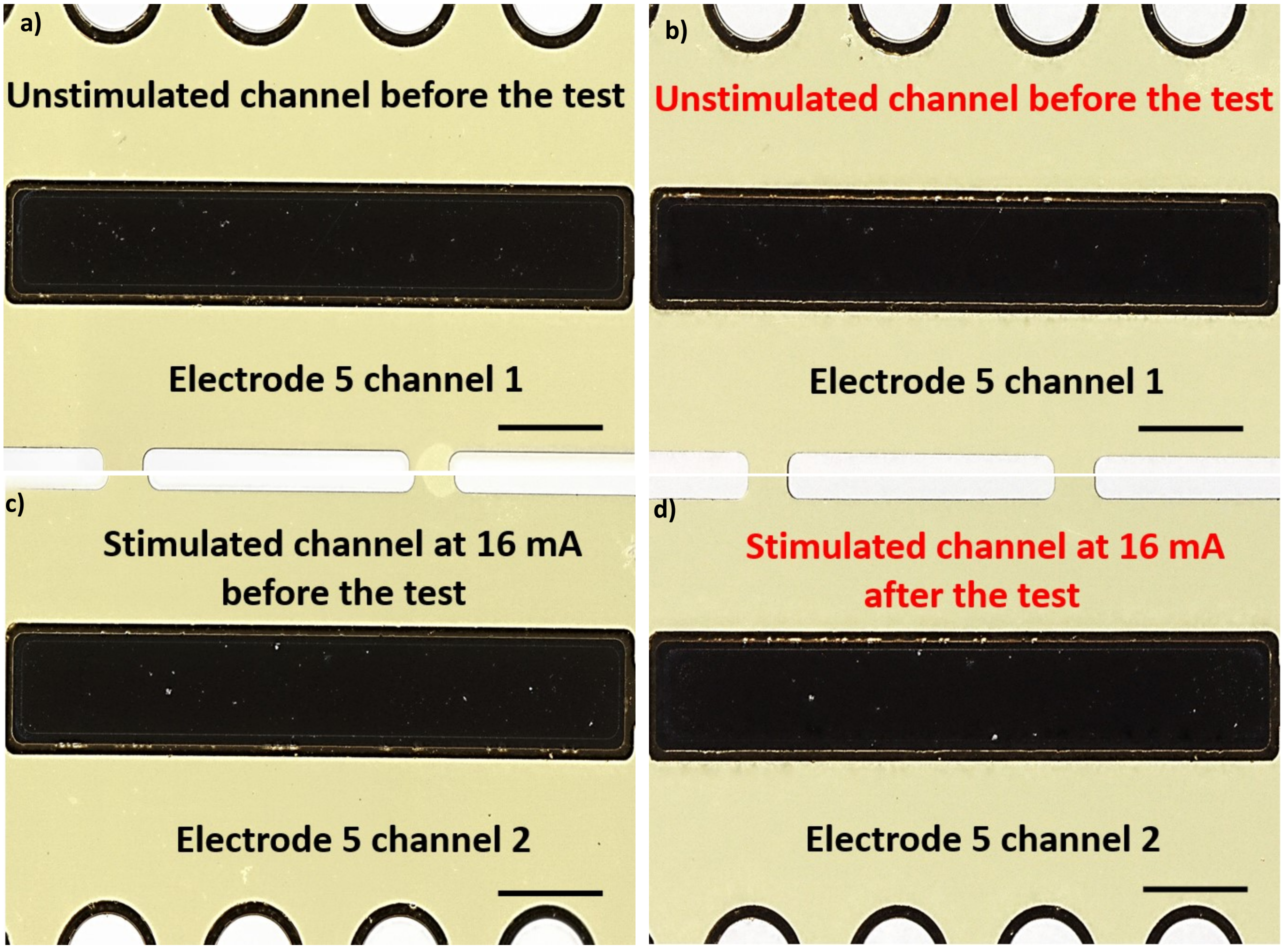


**Figure S28.** Optical image of *Flex* electrode 5 before and after the Stim-Stab test. a) the optical image of the unstimulated channel (electrode 5 channel 1) before the Stim-Stab test, b) the optical image of the unstimulated channel after the Stim-Stab test, c) the optical image of the stimulated channel (electrode 5 channel 2) before the Stim-Stab test, d) the optical image of the stimulated channel after the Stim-Stab test, scale bar=0.25 mm.

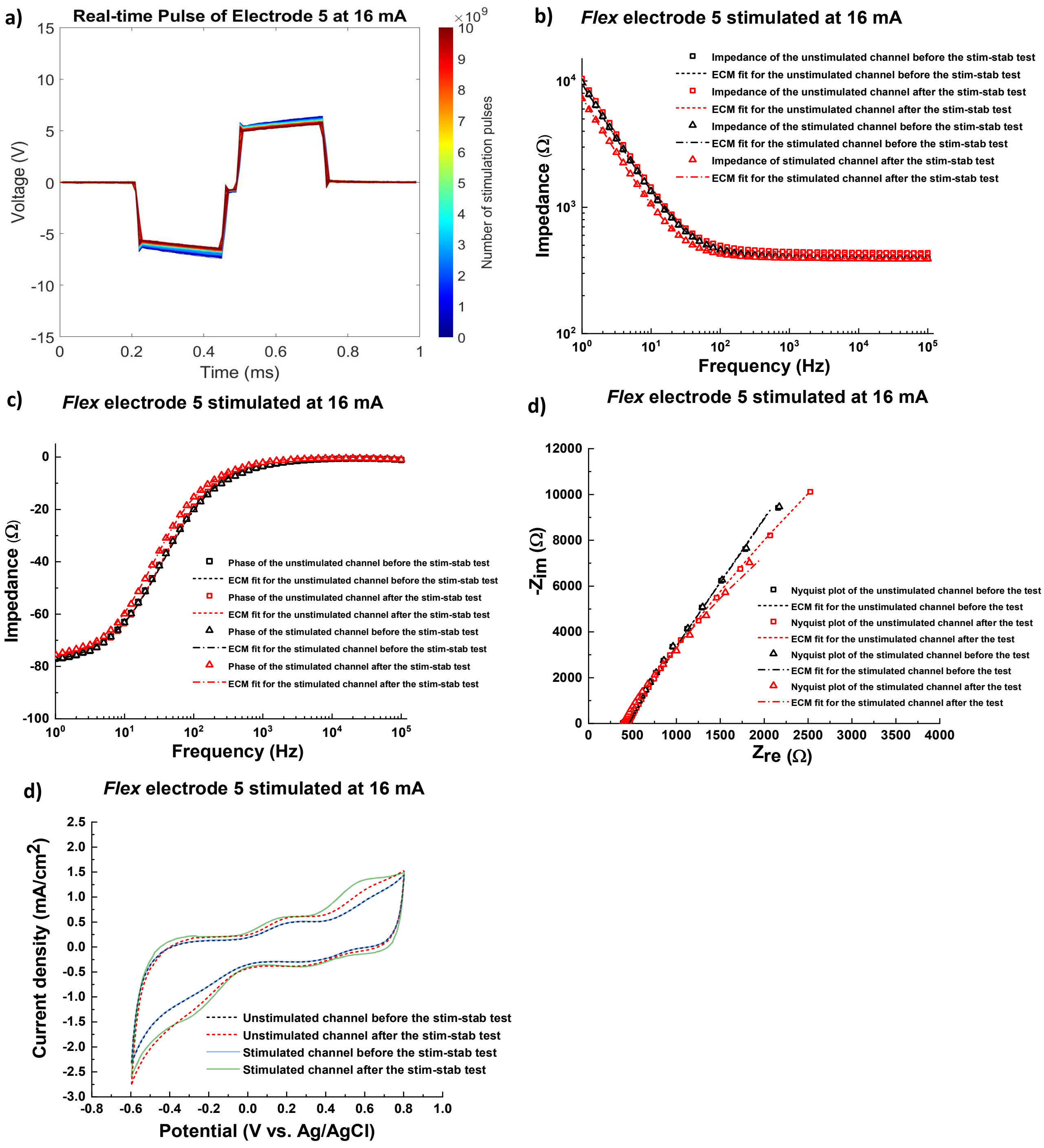


**Figure S29**. Real-time VT waveform, EIS and CV curves of *Flex* electrode 5. a) Real-time VT waveform of electrode 5 channel 2 stimulated at 16 mA as a function of the pulse number, b) impedance of electrode 5 before and after the Stim-Stab test, and the equivalent circuit model fit, c) phase of electrode 5 before and after the Stim-Stab test, and the equivalent circuit model fit, d) Nyquist plot of electrode 5 before and after the Stim-Stab test, and the equivalent circuit model fit, e) CV curves of electrode 5 before and after the Stim-Stab test.

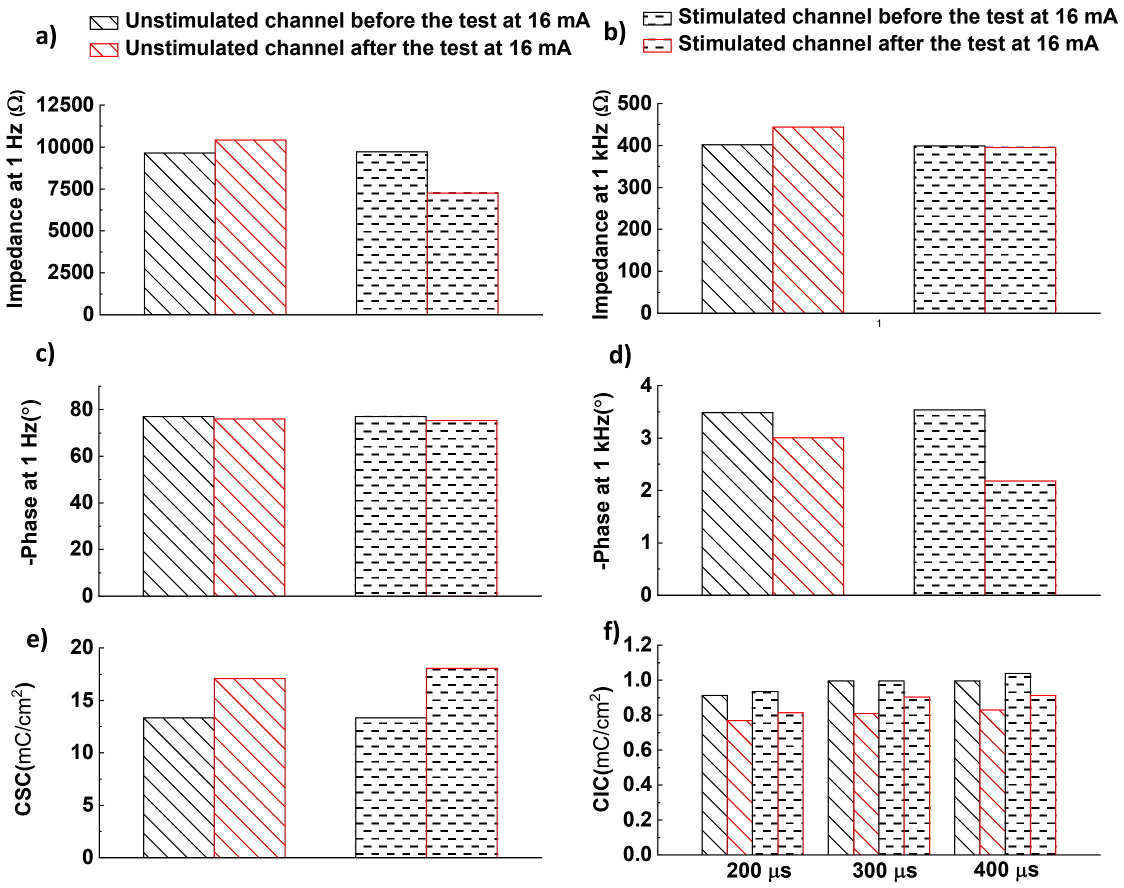


**Figure S30**. Electrochemical properties of *Flex* electrode 5 before and after the stim-stab test. a) 1-Hz impedance before and after the Stim-Stab test, b) $10^3$-Hz impedance before and after the Stim-Stab test, c) 1-Hz phase before and after the Stim-Stab test, d) $10^3$-Hz phase before and after the Stim-Stab test, e) CSC of *Flex* electrode 5 before and after the Stim-Stab test, f) CIC of *Flex* electrode 5 before and after the Stim-Stab test.

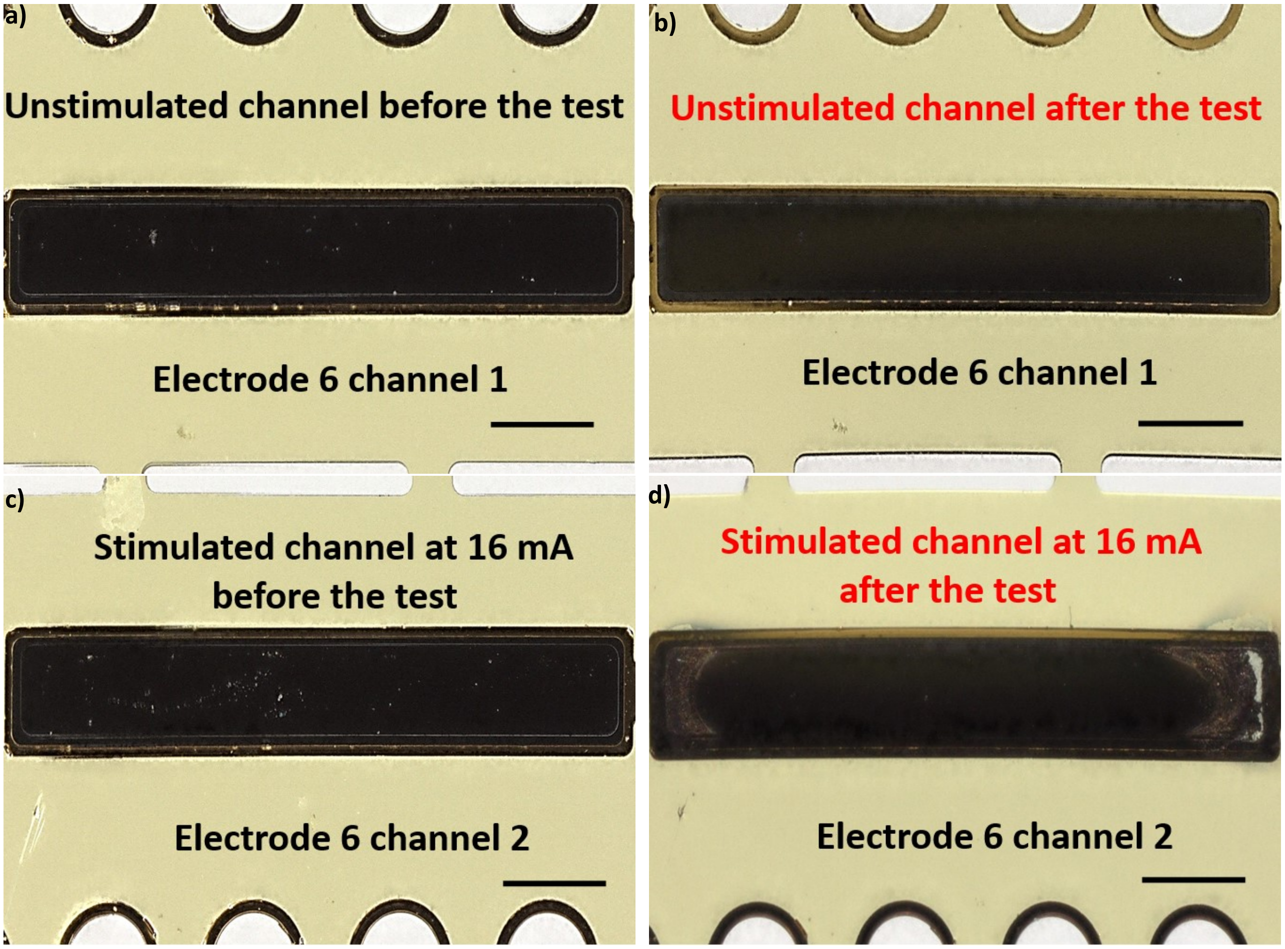


**Figure S31.** Optical image of *Flex* electrode 6 before and after the Stim-Stab test. a) the optical image of the unstimulated channel (electrode 6 channel 1) before the Stim-Stab test, b) the optical image of the unstimulated channel after the Stim-Stab test, c) the optical image of the stimulated channel (electrode 6 channel 2) before the Stim-Stab test, d) the optical image of the stimulated channel after the Stim-Stab test, scale bar=0.25 mm.

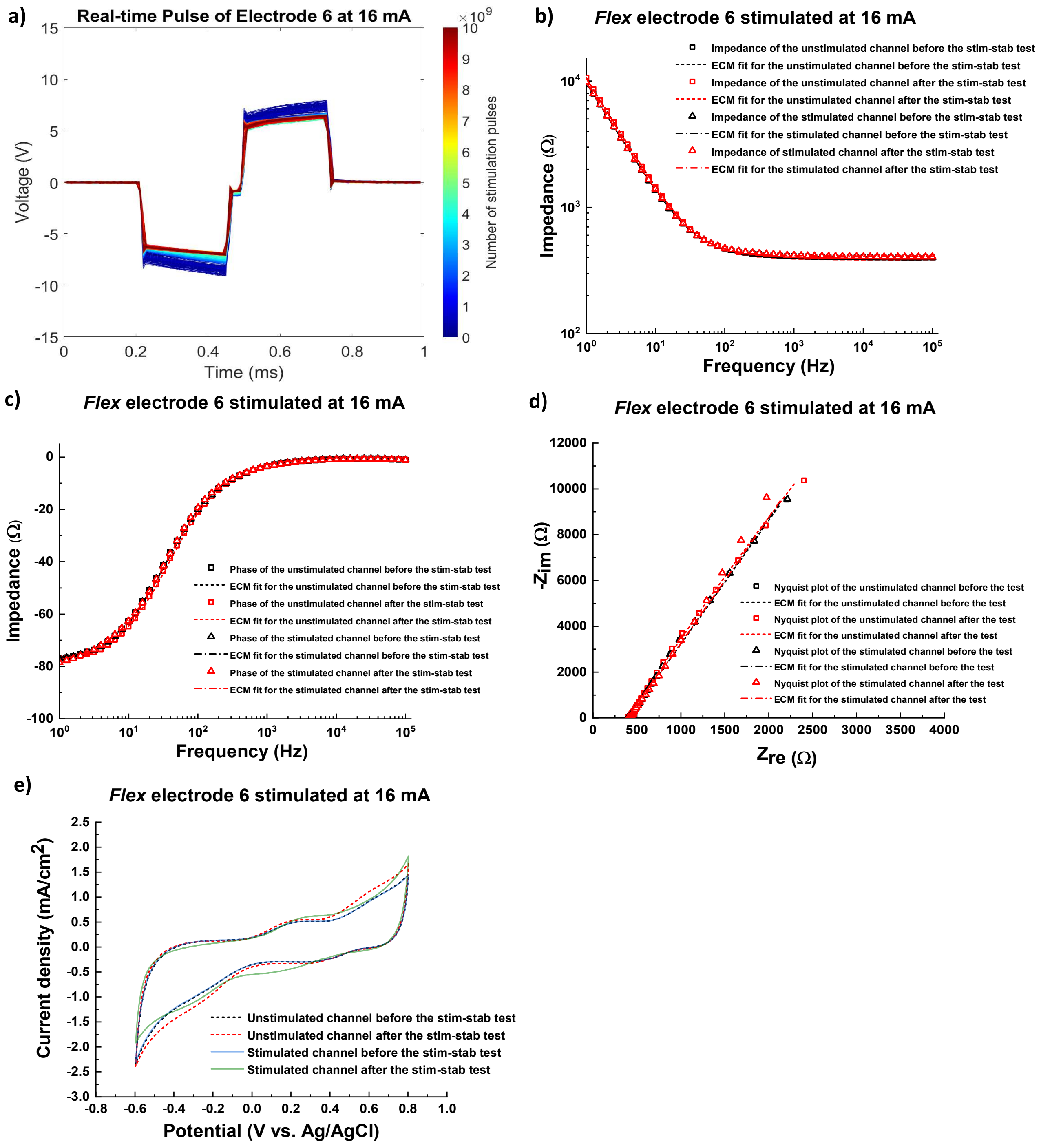


**Figure S32**. Real-time VT waveform, EIS and CV curves of *Flex* electrode 6. a) real-time VT waveform of electrode 6 channel 2 stimulated at 16 mA as a function of the pulse number, b) impedance of electrode 6 before and after the Stim-Stab test, and the equivalent circuit model fit, c) phase of electrode 6 before and after the Stim-Stab test, and the equivalent circuit model fit, d) Nyquist plot of electrode 6 before and after the Stim-Stab test, and the equivalent circuit model fit, e) CV curves of electrode 6 before and after the Stim-Stab test.

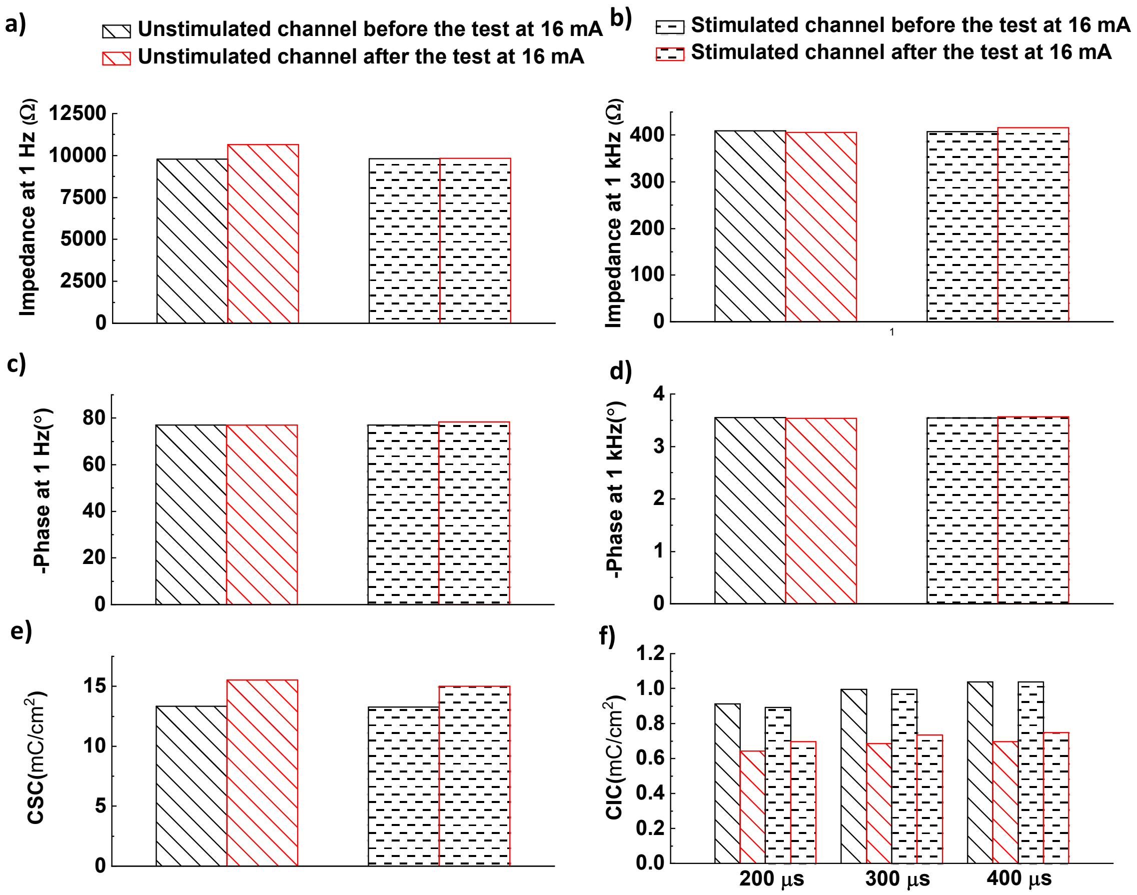


**Figure S33**. Electrochemical properties of *Flex* electrode 6 before and after the Stim-Stab test. a) 1-Hz impedance before and after the Stim-Stab test, b) $10^3$-Hz impedance before and after the Stim-Stab test, c) 1-Hz phase before and after the Stim-Stab test, d) $10^3$-Hz phase before and after the Stim-Stab test, e) CSC of *Flex* electrode 6 before and after the Stim-Stab test, f) CIC of *Flex* electrode 6 before and after the Stim-Stab test.

**Table S1.** Parameters of the ECM for unstimulated and stimulated electrode channels before and after the Stim-Stab test.

| 3-electrode configuration | Unstimulated channels before the Stim-Stab ( n = 6) | UnStimulated channels after the Stim-Stab (n = 6) | Electrode 5 channel 2 before the Stim-Stab | Electrode 5 channel 2 after the Stim-Stab | Electrode 6 channel 2 before the Stim-Stab | Electrode 6 channel 2 after the Stim-Stab | Electrode 3 channel 2 before the Stim-Stab | Electrode 3 channel 2 after the Stim-Stab |
|---|---|---|---|---|---|---|---|---|
| $R_s$ (Ω) | 446 ± 38 | 407 ± 16 | 394 | 392 | 403 | 410.1 | 455.9 | 358.5 |
| $R_{ct}$ (MΩ) | 3.1 ± 0.6 | 1.0 ± 0.6 | 2.3 | 0.1 | 3.8 | 0.5 | 3.4 | $2.276\times10^{-3}$ |
| $n$ | 0.855 ± 0.025 | 0.896 ±0.003 | 0.887 | 0.904 | 0.885 | 0.888 | 0.836 | 0.683 |
| $Q$ (pF) | 20.5 ± 0.4 | 19.7 ± 2.0 | 20.7 | 25.7 | 20.6 | 20.2 | 19.5 | 0.781 |
| $W$ (Ω/s$^{1/2}$) | - | - | - | - | - | - | - | $76.79\times10^{-6}$ |
